# Sexually dimorphic mechanisms of VGLUT-mediated protection from dopaminergic neurodegeneration

**DOI:** 10.1101/2023.10.02.560584

**Authors:** Silas A. Buck, Sophie A. Rubin, Tenzin Kunkhyen, Christoph D. Treiber, Xiangning Xue, Lief E. Fenno, Samuel J. Mabry, Varun R. Sundar, Zilu Yang, Divia Shah, Kyle D. Ketchesin, Darius D. Becker-Krail, Iaroslavna Vasylieva, Megan C. Smith, Florian J. Weisel, Wenjia Wang, M. Quincy Erickson-Oberg, Emma I. O’Leary, Eshan Aravind, Charu Ramakrishnan, Yoon Seok Kim, Yanying Wu, Matthias Quick, Jonathan A. Coleman, William A. MacDonald, Rania Elbakri, Abby L. Olsen, Emily M. Rocha, Gary W. Miller, Briana R. De Miranda, Michael J. Palladino, Brian D. McCabe, Kenneth N. Fish, Marianne L. Seney, Stephen Rayport, Susana Mingote, Karl Deisseroth, Thomas S. Hnasko, Rajeshwar Awatramani, Alan M. Watson, Scott Waddell, Claire E. J. Cheetham, Ryan W. Logan, Zachary Freyberg

**Author notes:** Correspondence (Z.F.). Department of Biology, University of Oxford, Oxford OX1 3RB, UK.

## Abstract

Parkinson’s disease (PD) targets some dopamine (DA) neurons more than others. Sex differences offer insights, with females more protected from DA neurodegeneration. The mammalian vesicular glutamate transporter VGLUT2 and *Drosophila* ortholog dVGLUT have been implicated as modulators of DA neuron resilience. However, the mechanisms by which VGLUT2/dVGLUT protects DA neurons remain unknown. We discovered DA neuron dVGLUT knockdown increased mitochondrial reactive oxygen species in a sexually dimorphic manner in response to depolarization or paraquat-induced stress, males being especially affected. DA neuron dVGLUT also reduced ATP biosynthetic burden during depolarization. RNA sequencing of VGLUT^+^ DA neurons in mice and flies identified candidate genes that we functionally screened to further dissect VGLUT-mediated DA neuron resilience across PD models. We discovered transcription factors modulating dVGLUT-dependent DA neuroprotection and identified dj-1β as a regulator of sex-specific DA neuron dVGLUT expression. Overall, VGLUT protects DA neurons from PD-associated degeneration by maintaining mitochondrial health.

## Introduction

Parkinson’s disease (PD) is an age-related neurodegenerative disorder which represents one of the fastest growing neurologic diseases today^1^. Midbrain dopamine (DA) neurons are specifically vulnerable to degeneration in PD^2,3^, though this degeneration is not uniform. DA neurons in the ventral tegmental area (VTA) are more protected from degeneration relative to neighboring DA neurons in the substantia nigra *pars compacta* (SNc)^2–9^. Yet, the mechanisms for these region-specific differences in resilience remain poorly understood.

Sex differences in DA neuron resilience provide important insights for uncovering new mechanisms of selective DA neuroprotection. PD affects more men than women and has an earlier age of symptomatic onset for men^10–18^. Moreover, while rodent PD models have traditionally focused on males^19^, in studies comparing both sexes, female rodents exhibit less SNc DA neuron degeneration and fewer motor symptoms, similar to humans^19–22^. Additional clues are offered by a distinct subpopulation of DA neurons that express the vesicular glutamate transporter 2 (VGLUT2) and co-transmit glutamate. The highest proportion of VGLUT2-expressing DA neurons is found in the medial VTA, with a smaller population in the lateral SNc or SN *pars lateralis*^23–28^ – DAergic midbrain regions that are relatively spared in PD patients as well as in primate and rodent PD models^2–7,9,29–32^. Indeed, VGLUT2-expressing (VGLUT2^+^) DA neurons are more likely to survive insults in preclinical PD models compared to DA neurons that do not express VGLUT2 (VGLUT2^−^)^8,33–35^. In rodent models of acute neurotoxicant-induced DA neurodegeneration, conditional knockout (cKO) of VGLUT2 in DA neurons renders these cells more vulnerable to degeneration by 6-hydroxydopamine (6-OHDA) or 1-methyl-4-phenyl-1,2,3,6-tetrahydropyridine (MPTP)^8,33^. Such results are also consistent with recent human postmortem data showing that VGLUT2-expressing DA neurons are more likely to survive in PD patients^35^. Importantly, these findings are well-conserved across evolution since, in flies, we discovered that the *Drosophila* ortholog of VGLUT2, *Drosophila* VGLUT (dVGLUT), mediates sex differences in DA neuron resilience. Diminishing dVGLUT expression in DA neurons reduces females’ greater resilience to age-related DA neuron loss^36^. This is in line with evidence that females express more VGLUT2 or dVGLUT in DA neurons compared to males – a finding conserved across flies, rodents, and humans^36^.

Expression of DA neuron VGLUT2 and dVGLUT is highly dynamic, particularly during development and in response to cell stress or injury. Almost all midbrain DA neurons express VGLUT2 during development^8^. However, by adulthood, most midbrain DA neurons repress or ‘turn off’ VGLUT2 expression^24–26^. In contrast, medial VTA and lateral SNc DA neurons maintain VGLUT2 expression throughout life. Significantly, in adult rodents, VGLUT2 expression re-emerges in surviving midbrain DA neurons in response to 6-OHDA or MPTP injury^8,33^. Similarly, in adult flies, DA neuron dVGLUT expression dynamically increases throughout aging, providing DA neuroprotection^36^. Together, findings across flies, rodents, and humans suggest that: 1) VGLUT (VGLUT2 in mammals and dVGLUT in *Drosophila*) expression in DA neurons represents an adaptive compensatory response to DA neuron insults to boost DA neuron resilience and prevent cell death; and 2) interfering with VGLUT2 expression may increase the extent and/or rate of DA neuron degeneration in PD. Yet, this also raises significant questions concerning the fundamental mechanisms responsible for VGLUT’s impact on DA neurons and the sex differences associated with DA neuron resilience in PD.

Here, we used a combination of imaging, genetics, and pharmacology in the genetically tractable *Drosophila* system to determine the mechanisms by which VGLUT confers resilience to PD-related DA neuron degeneration in males and females. We used the paraquat model of PD since paraquat is one of the most widely used pesticides globally^37^, and occupational paraquat exposure substantially increases PD risk^38^. Paraquat causes DA neurotoxicity by generating reactive oxygen species (ROS) that impair mitochondrial function^39–43^. As in mammals, paraquat exposure causes selective DA neuron degeneration in the fly brain^44–46^, which enabled us to identify conserved mechanisms of DA neuron resilience. We discovered that DA neuron dVGLUT modulated levels of mitochondrial ROS in response to paraquat exposure, with males specifically affected. Further, knockdown of dVGLUT in DA neurons increased ROS generation in response to neuronal depolarization, suggesting dVGLUT modulates mitochondrial stress related to increased ATP production. Consistent with this, we found that dVGLUT lessened ATP biosynthetic burden in DAergic terminals during firing. These results emphasize a critical role for DA neuron dVGLUT as a modulator of mitochondrial function. Finally, using comparative RNA sequencing (RNAseq) approaches in mice and flies, we identified differentially expressed (DE) genes enriched in VGLUT-expressing DA neurons that were evolutionarily conserved. A forward genetic screen of these DE genes for resilience across several PD models revealed genes primarily associated with mitochondrial metabolism and transcriptional regulation. Significantly, we identified the *PARK7* ortholog, *dj-1β*, as a novel regulator of sex-specific DA neuron dVGLUT expression. Together, our results indicate VGLUT is a central mediator of DA neuron resilience that protects DA neurons by maintaining mitochondrial health, and sexual dimorphism in this mechanism of neuroprotection may explain females’ greater resilience in PD.

## Results

### dVGLUT-expressing DA neurons localize to distinct cell clusters

To characterize dVGLUT-dependent mechanisms of DA neuron resilience, we first identified DA neuron populations that co-express dVGLUT throughout the adult *Drosophila* brain. Neurons that co-expressed *dVGLUT* alongside tyrosine hydroxylase (TH; encoded by the *ple* gene), the rate limiting enzyme in the production of DA, were labeled in whole wild-type fly brains via multiplex RNAscope fluorescent *in situ* hybridization (Figure 1). We focused on several DA neuron clusters including within the protocerebral anterior lateral (PAL), protocerebral anterior medial (PAM), and subesophageal ganglion (SOG) regions in the anterior aspect of the adult fly brain. In parallel, we also examined the protocerebral posterior medial 1-3 (PPM1-3) and protocerebral posterior lateral 1 and 2 (PPL1, PPL2) clusters in the posterior aspect of the brain^47^ (Figure 1A). We found dVGLUT co-expression in PPM1 DA neurons, as well as in a subset of PPL1 DA neurons, consistent with earlier findings^48^. Importantly, we discovered previously undescribed populations of TH^+^/dVGLUT^+^ DA neurons in the lateral SOG and within the tritocerebrum (T1), as well as in PPL2, including PPL2ab and PPL2c subgroups (Figure 1B). These data provide the first comprehensive map of DA neuron dVGLUT expression, establishing four unique clusters of TH^+^/dVGLUT^+^ DA neurons in the adult fly brain.

**Figure 1.**
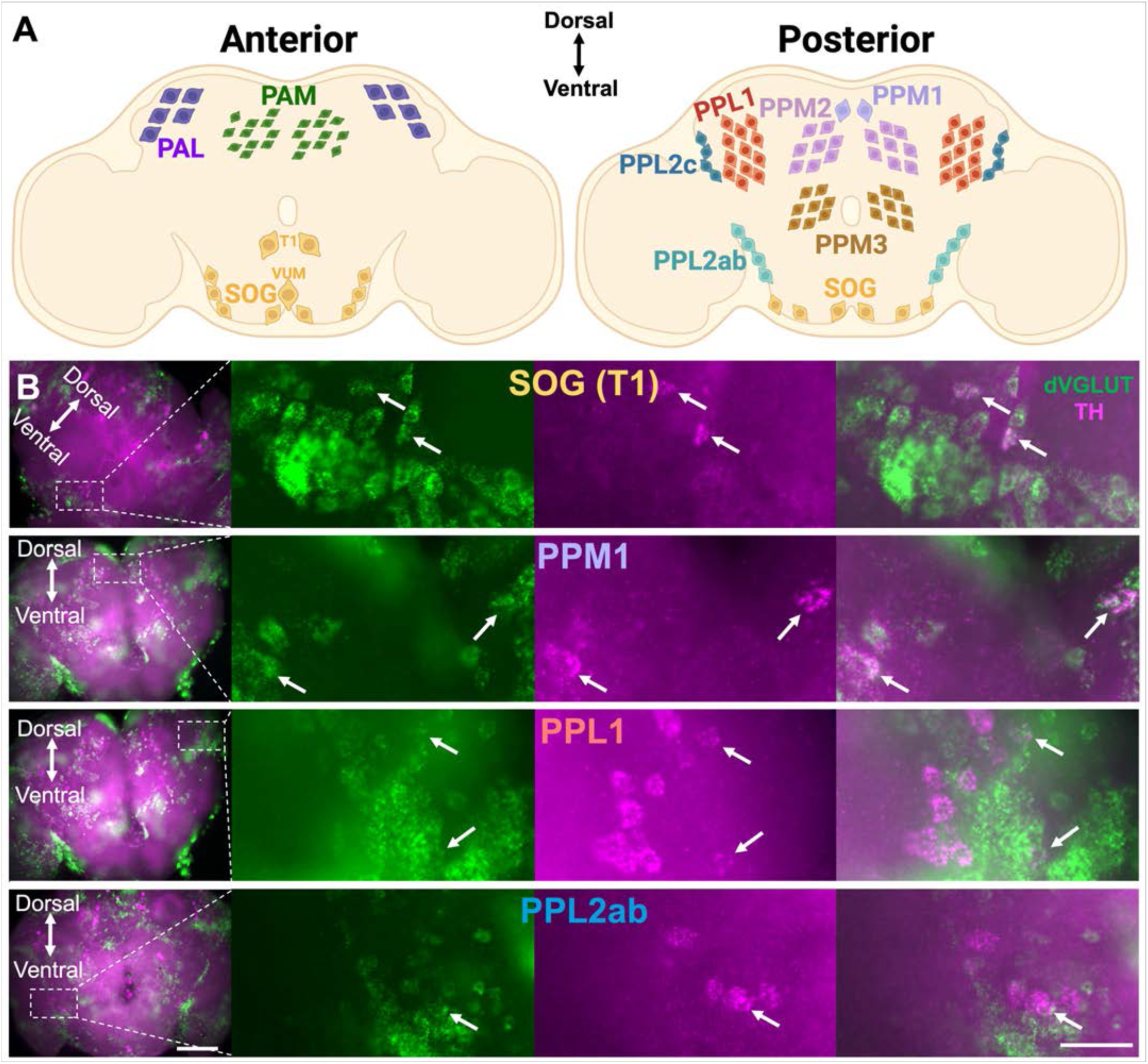
dVGLUT-expressing DA neurons localize to distinct cell clusters. **(A)** Illustrations of DA neuron clusters in the anterior and posterior regions of the adult *Drosophila* brain including: subesophageal ganglion (SOG), tritocerebrum (T1), protocerebral posterior medial 1 (PPM1), protocerebral posterior lateral 1 (PPL1), and protocerebral posterior lateral 2 (PPL2) clusters. **(B)** Representative multiplex RNAscope 4x images (left) and magnified 20x views of insets (3 right panels) of wild-type w^1118^ fly brains (14d post-eclosion), displaying *dVGLUT* (green) and *TH* (magenta) mRNA co-expression in several DA neuron clusters. TH^+^/dVGLUT^+^ cells were found in PPL2 subclusters comprised of both PPL2ab and PPL2c. 4x scale bar=100μm, 20x scale bar=25μm.

### Dynamic regulation of dVGLUT expression mediates sex-specific DA neuron resilience

To dissect dVGLUT’s roles in selective DA neuron resilience, we first determined the impacts of different cell stressors on DA neuron dVGLUT expression in males and females. We used an intersectional genetic luciferase reporter, which we previously validated^36^, to indirectly measure changes in DA neuron dVGLUT expression (Figure 2A). We tested our intersectional dVGLUT reporter using the paraquat model of PD^44^. When reduced, paraquat is specifically taken up into DA neurons by the DA transporter (DAT) where it induces oxidative stress, eventually leading to DA neuron degeneration^41–43^. Using this model, we asked: 1) whether DA neuron dVGLUT expression changes in response to paraquat exposure across time, 2) if advancing age modifies this potentially adaptive dVGLUT response, and 3) whether paraquat-induced alterations in DA neuron dVGLUT expression are sex-specific.

**Figure 2.**
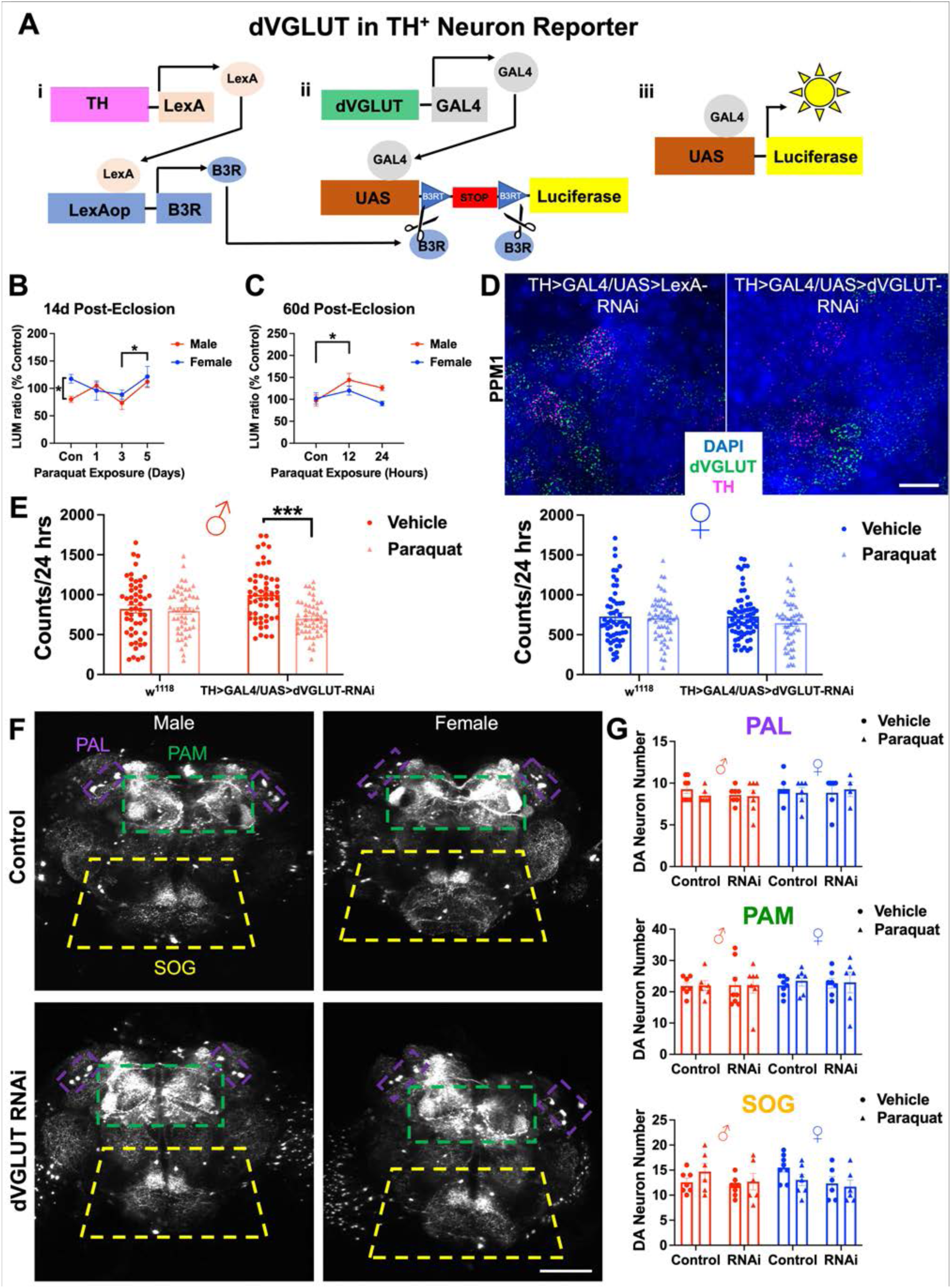
Dynamic regulation of dVGLUT expression mediates sex-specific DA neuron resilience. **(A)** Schematic of the intersectional luciferase reporter to estimate dVGLUT expression in DA neurons (dVGLUT>GAL4/LexAop>B3R;TH>LexA/UAS>B3RT-STOP-B3RT-Luciferase). Panel i: The *TH* promoter drives LexA to express B3 recombinase (B3R) in TH^+^ DA neurons. Panel ii: B3R excises a transcriptional stop cassette within UAS>Luciferase. Panel iii: This excision permits successful dVGLUT>GAL4-driven transactivation of UAS>Luciferase specifically in TH^+^/dVGLUT^+^ cells. **(B)** Among adult flies (14d post-eclosion), control (Con) females exhibited higher baseline DA neuron dVGLUT expression compared to males (p=0.048, Bonferroni multiple comparisons test). 5d exposure to 10mM paraquat significantly raised DA neuron dVGLUT expression compared to 3d exposure (p=0.047, Bonferroni multiple comparisons test). Luminescent DA neuron dVGLUT reporter data were normalized to % untreated male and female controls. N=4-8 homogenates of 5 brains per group. **(C)** Among older flies (60d post-eclosion), paraquat exposure (10mM, 12h) significantly raised DA neuron dVGLUT expression compared to the vehicle-treated Con group (p=0.039, Bonferroni multiple comparisons test). N=4-8 homogenates of 5 brains per group. Related analyses are shown in Figure S1. **(D)** Representative images showing *dVGLUT* (green) and *TH* (magenta) mRNA expression in TH-driven Control (TH>GAL4/UAS>LexA-RNAi) and dVGLUT RNAi (TH>GAL4/UAS>dVGLUT-RNAi) flies at 14d post-eclosion. Scale bar=10μm. Related analyses are shown in Table S1. **(E)** At day 2 of 10mM paraquat exposure, a genotype × paraquat interaction was observed in male flies (4d post-eclosion) (left panel; F_1,202_=10.3, p=0.0016, two-way ANOVA). Male TH-driven dVGLUT RNAi flies exhibited decreased locomotion in response to paraquat (p<0.001), but male w^1118^ wild-type control flies did not (p>0.05, Bonferroni multiple comparisons test). Female w^1118^ wild-type and dVGLUT RNAi flies were not significantly affected by paraquat (right panel; p>0.05, Bonferroni multiple comparisons test). N=50-67 flies per group. **(F)** Representative projection images of GFP-labeled DA neurons in TH-driven Control (TH>GAL4,UAS>GFP/UAS>LexA-RNAi) and dVGLUT RNAi (TH>GAL4,UAS>GFP/UAS>dVGLUT-RNAi) brains (14d post-eclosion). Images are from protocerebral anterior lateral (PAL, purple), protocerebral anterior medial (PAM, green) and subesophageal ganglion (SOG, yellow) clusters. Scale bar=100μm. **(G)** There was no effect of paraquat exposure (10mM, 5d) on DA neuron number (p>0.05, three-way ANOVA). There was a significant effect of dVGLUT RNAi on SOG DA neuron number (F_1,46_=4.95, p=0.031, three-way ANOVA). N=4-8 per group. *p<0.05, ***p<0.001; all data are plotted as Mean±SEM.

Examination of vehicle-treated flies revealed that DA neuron dVGLUT expression increased with age (p=0.0064; Figure S1A) and was significantly higher in 14d-old adult females than in males (p=0.048; Figure 2B), confirming earlier observations^36^. In contrast, sex differences in DA neuron dVGLUT expression were absent in aged flies (60d post-eclosion, p>0.05; Figure 2C). Duration of paraquat exposure also significantly modified DA neuron dVGLUT reporter expression. In younger flies, DA neuron dVGLUT expression was greater after 5d versus 3d of paraquat exposure (p=0.047; Figure 2B). In aged flies (60d post-eclosion), we identified elevated DA neuron dVGLUT expression even earlier compared to younger flies, with significantly increased DA neuron dVGLUT expression following 12h of paraquat exposure (p=0.039; Figure 2C). In parallel, we examined whether paraquat exposure also altered TH expression in dVGLUT^+^ DA neurons. We therefore developed an intersectional luciferase reporter to measure changes in TH expression specifically in TH^+^/dVGLUT^+^ DA neurons (Figure S1B) which we compared to a non-intersectional reporter of global TH expression in all TH-expressing DA neurons (Figure S1C). Besides providing an indirect measure of DA biosynthetic capacity, the global TH reporter also offers a proxy of total DA neuron number throughout the fly brain. Validation of the non-intersectional TH reporter demonstrated a strong luminescent signal with high signal-to-noise compared to undriven controls (Figure S1D). Using both TH reporters, we discovered that paraquat exposure did not alter TH expression in either TH^+^/dVGLUT^+^ DA neurons or in the larger DA neuron population in 14d-old males and females (p>0.05; Figure S1E). However, TH expression in aged 60d-old flies was increased in a sex-dependent manner (Figure S1F). Paraquat exposure significantly elevated TH expression in TH^+^/dVGLUT^+^ neurons of aged males compared to females (p=0.048) with similar sex differences evident in the broader DA neuron population (p=0.0002; Figure S1F). Notably, we found an overall decrease in global TH expression across time of exposure (p=0.033; Figure S1F), suggesting paraquat-induced DAergic degeneration. Finally, direct comparison of intersectional versus global TH reporters revealed that paraquat-induced decreases in TH expression were not evident in TH^+^/dVGLUT^+^ neurons but instead were mainly driven by expression changes across the broader DA neuron population. Together, these data suggest that DA neuron dVGLUT expression protects against paraquat-induced diminishment of DA neuron function or DA neuron loss, as indicated by decreased TH expression (Figure S1F).

To probe dVGLUT’s role more directly in DA neuron vulnerability to early paraquat exposure, we knocked down DA neuron dVGLUT expression using RNA interference (RNAi). We confirmed RNAi-mediated dVGLUT knockdown in DA neurons via multiplex RNAscope, finding a significant decrease (48.8-69.0%) in the percentage of TH^+^ DA neurons that co-express dVGLUT compared to controls (Figure 2D and Table S1), consistent with earlier work^36^. Moreover, there was a 31.6-51.4% reduction in dVGLUT mRNA grains in TH^+^ cells compared to all control lines tested, further validating the efficacy of RNAi-mediated dVGLUT knockdown in DA neurons (Figure 2D; Table S1). We then examined the impact of DA neuron dVGLUT knockdown on locomotion in response to early paraquat exposure (2d, 10mM). In males, we found a significant genotype × paraquat interaction (p=0.006), where paraquat decreased locomotion in dVGLUT RNAi flies (p<0.001), but not in wild-type controls (p>0.05; Figure 2E). In comparison, females did not exhibit paraquat-induced locomotor deficits in either control or RNAi flies (p>0.05; Figure 2E). Since RNAi-mediated knockdown does not entirely eliminate DA neuron dVGLUT expression, we posit that the higher levels of dVGLUT expressed in adult female DA neurons compared to males provide sufficient protection against paraquat-induced locomotor impairments.

Finally, we examined whether: 1) paraquat-induced alterations in DA-mediated behavior translated to changes in DA neuron number, and 2) the impact of DA neuron dVGLUT knockdown on these relationships. We focused on the three anterior DA neuron clusters in the PAL, PAM, and SOG brain regions since differences in DA neuron resilience have been reported in these clusters, including in *Drosophila* models of PD and age-related cell loss^36,49^. We found no significant DA neuron loss in any of the three clusters in response to 5d of paraquat exposure in 14d-old males and females (p>0.05; Figures 2F and 2G). These results suggest that at this timepoint, DA neurons are undergoing paraquat-induced stress but have not yet undergone degeneration, consistent with our TH reporter data. We also observed that DA neuron dVGLUT RNAi knockdown did not significantly alter DA neuron death at this early point in paraquat exposure in either sex (p>0.05; Figures 2F and 2G). Interestingly, a main effect of RNAi was observed, where DA neuron dVGLUT RNAi flies displayed fewer SOG DA neurons than control flies across sexes and treatment groups (Control: 14.0±0.6, RNAi: 12.1±0.6; p=0.031), indicating a potential developmental effect of dVGLUT RNAi on SOG DA neurons. Together, these results demonstrate that, as in mammalian PD models and clinical PD, dVGLUT confers protection to DA neurons in *Drosophila*. More specifically, loss of DA neuron dVGLUT expression renders males more vulnerable to DAergic insults, including paraquat in addition to aging^36^. The early time periods of paraquat exposure that we examined (≤5d) when DA neurons are stressed but still intact suggest dVGLUT-dependent mechanisms of neuroprotection occur during critical windows of vulnerability prior to later irreversible loss of DA neurons.

### DA neuron dVGLUT modulates sex differences in mitochondrial oxidative stress

Paraquat damages DA neurons through redox cycling^50^. This generates ROS that boosts oxidative stress, causing mitochondrial dysfunction and ultimately DA neuron loss^41^. Critically, mitochondrial ROS-mediated stress is a key aspect of PD pathogenesis^51,52^. We therefore hypothesized that dVGLUT mediates DA neuron resilience by moderating mitochondrial oxidative stress in response to paraquat. To test this, we employed the MitoTimer ROS biosensor, a mitochondrial matrix-localized dsRed mutant that shifts fluorescence from green to red when oxidized. Thus, measuring the red:green fluorescence ratio (red:green ratio) in DA neurons within whole living fly brains enables MitoTimer to serve as an *in vivo* reporter of accumulating ROS during mitochondrial stress^53,54^. We imaged TH-driven MitoTimer in DAergic projections to the SOG, which are known to originate from PPM1 DA neurons^55^ and local SOG DA neurons^55–57^ – two DA neuron clusters that are majority dVGLUT^+^ (Figure 1). For comparison, we also imaged DAergic projections to the ellipsoid body (EB) and fan-shaped body (FSB). These central complex structures are innervated mainly by dVGLUT^−^ PPM3 DA neurons^47^ (Figure 1) and play key roles in DA-mediated navigation and goal-oriented behaviors, resembling the functions played by the mammalian basal ganglia^58–61^. Control flies co-expressing either Luciferase or LexA RNAi alongside MitoTimer in DA neurons showed strong regional differences in baseline MitoTimer red:green ratio. Indeed, the mainly dVGLUT^+^ DAergic projections to the SOG possessed a lower red:green ratio compared to the dVGLUT^−^ DA neuron projections to the EB (p<0.001) and FSB (p<0.001; Figures S2A and S2B). This finding therefore correlated DA neuron dVGLUT expression with lower mitochondrial ROS in DAergic projections. Importantly, control females possessed a lower baseline red:green ratio than males across all regions (p=0.0071; Figures S2A and S2B), suggesting less mitochondrial ROS in female flies at baseline. When we exposed control flies to paraquat (5d, 10mM), we found a significant increase in MitoTimer red:green ratio in DAergic projections to the FSB across sexes (p=0.038), but no change was found in the SOG or EB (p>0.05; Figures S2C and S2D).

We next examined the impact of DA neuron dVGLUT knockdown on accumulation of mitochondrial ROS in DAergic projections in response to paraquat exposure (5d, 10mM). Projections to the SOG and EB regions did not show paraquat-induced changes in MitoTimer signal in either males or females (p>0.05; Figures S3A and S3B). In contrast, there was a significant paraquat-induced increase in the MitoTimer red:green ratio in DAergic projections to the FSB of male TH-driven dVGLUT RNAi flies that was absent in females (p=0.0049; Figures 3A and 3B). Since changes in the MitoTimer red:green ratio may also be attributed to differences in mitochondrial turnover, we confirmed our results were the product of elevated mitochondrial ROS by exposing flies to hydrogen peroxide (H_2_O_2_), another agent that induces mitochondrial oxidative stress via free radical formation^62,63^. In response to H_2_O_2_ exposure (3%, 2d), we similarly observed a significant increase in the MitoTimer red:green ratio in DAergic projections to the FSB in male dVGLUT RNAi flies (p<0.001; Figures S3C, S3D), replicating our paraquat findings. As with paraquat, females remained protected even when dVGLUT was knocked down in DA neurons (p>0.05; Figures S3C and S3D).

**Figure 3.**
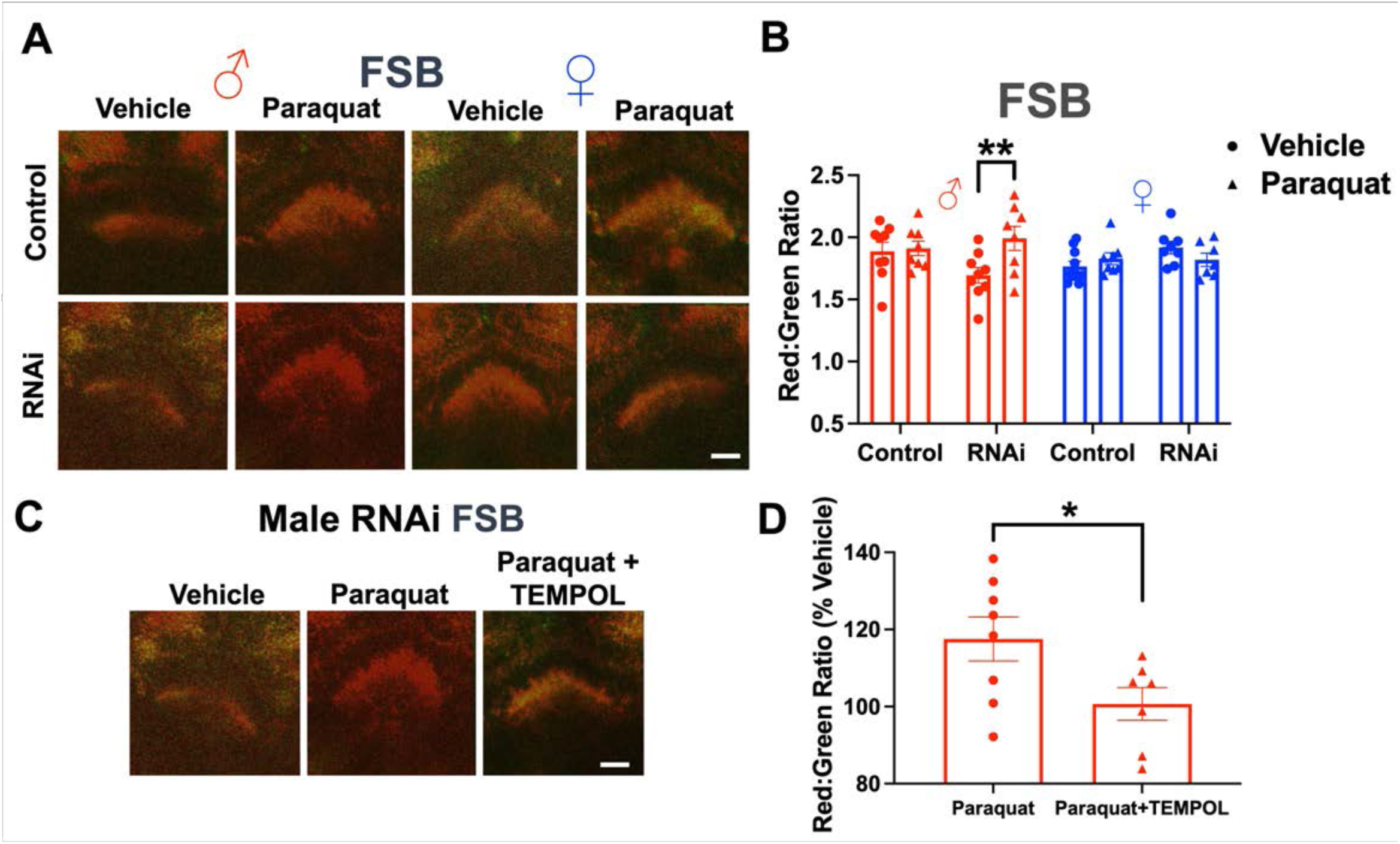
DA neuron dVGLUT modulates sex differences in mitochondrial oxidative stress. **(A)** Representative projection images of MitoTimer-labeled DAergic projections to the fan-shaped body (FSB) of male and female dVGLUT RNAi flies (TH>GAL4,UAS>MitoTimer/UAS>dVGLUT-RNAi) versus Control (TH>GAL4,UAS>MitoTimer/UAS>LexA RNAi) flies (14d post-eclosion). Green and red fluorescent channels were overlaid to generate final images. All groups were exposed to either paraquat (10mM, 5d) or vehicle control (0mM, 5d). Scale bar=25μm. **(B)** Paraquat exposure (10mM, 5d) significantly increased the MitoTimer red:green ratio in DAergic projections to the FSB of male dVGLUT RNAi flies compared to vehicle (paraquat × sex × RNAi interaction: F_1,59_=6.05, p=0.017, three-way ANOVA; male RNAi vehicle versus 10mM: p=0.0049, Bonferroni multiple comparisons test). N=7-10 per group. **(C)** Representative projection images of MitoTimer-labeled DAergic projections to the FSB of adult male dVGLUT RNAi flies. Flies were exposed to 5d of either 10 mM paraquat, 10 mM paraquat + antioxidant 3mM TEMPOL, or vehicle control. Scale bar=25μm. **(D)** Co-treatment with 3mM TEMPOL blocked paraquat-induced increases in the MitoTimer red:green ratio in male RNAi flies (p=0.038, unpaired t-test). N=7-8 per group. *p<0.05, **p<0.01; all data are plotted as Mean±SEM. See also Figures S2, S3 and S4.

We additionally determined if a chronic stressor like aging also leads to dVGLUT-mediated DA neuron resilience to mitochondrial oxidative stress/ROS. There was a significant interaction between brain region and age (p<0.001) on the MitoTimer red:green ratio (Figure S4A and S4B). DAergic projections to the SOG were more protected from age-related mitochondrial oxidative stress versus projections to the EB and FSB (Figures S4C), suggesting dVGLUT-expressing DAergic projections are resilient to age-associated mitochondrial ROS accumulation. However, we found no significant effects of sex or DA neuron dVGLUT RNAi (p>0.05). This suggests different and/or additional compensatory neuroprotective mechanisms capable of handling the chronic accumulation of mitochondrial ROS across aging compared to acute oxidative stress in response to paraquat.

Finally, we aimed to attenuate paraquat- and H_2_O_2_-specific increases in DA neuron mitochondrial oxidation by co-treating flies with the antioxidant 4-hydroxy-2,2,6,6-tetramethylpiperidin-1-oxyl (TEMPOL), a ROS metabolizer^64^. Exposing flies to TEMPOL (3mM, 5d) significantly lowered the MitoTimer red:green ratio in DA neurons across all sampled brain regions in control flies (p=0.0066; Figures S4D and S4E). Importantly, 3mM TEMPOL almost completely reversed the ∼20% increase in MitoTimer red:green ratio induced by paraquat or H_2_O_2_ in the FSB of male DA neuron dVGLUT RNAi flies, bringing the ratio back to levels observed with vehicle (Paraquat: p=0.038; Figures 3C and 3D; H_2_O_2_: p=0.0024, Figure S4F). In all, these findings demonstrate that 1) DA neuron dVGLUT expression is associated with baseline differences in mitochondrial ROS in DAergic projections, and 2) dVGLUT mediates male DA neuron vulnerability to PD-associated mitochondrial oxidative stress that occurs prior to DA neuron degeneration.

### dVGLUT modulates activity-driven intracellular ATP and mitochondrial ROS generation during DA neuron depolarization

Mitochondrial ROS is an intrinsic by-product of mitochondrial ATP synthesis, and increased ROS generation is observed during metabolically demanding states like neuronal depolarization^65,66^. We therefore hypothesized that dVGLUT expression impacts mitochondrial ROS levels in DA neurons via modulation of activity-driven mitochondrial ATP production. To test this, we examined the impact of DA neuron dVGLUT knockdown on activity-dependent ATP production in DA neurons via iATPSnFR, a genetically encoded fluorescent reporter of cytosolic ATP^67,68^. Both male and female dVGLUT RNAi flies exhibited significant increases in iATPSnFR fluorescence (iATPSnFr peak Λ1F/F*_i_*) within DAergic projections to the SOG in response to potassium chloride (KCl)-induced depolarization (p=0.024; Figures 4A, 4B and S5A). These data indicate a role for DA neuron dVGLUT in modulating activity-dependent biosynthesis and/or accumulation of intracellular ATP. There was no significant effect of dVGLUT RNAi on activity-driven iATPSnFR fluorescence changes in the EB or FSB (p>0.05; Figures S5B and S5C). Interestingly, regional differences in iATPSnFR signal for each genotype/sex group were strongly correlated to baseline MitoTimer red:green ratio in the same DAergic projections (r^2^=0.998, p=0.034; Figure 4C), further suggesting a relationship between activity-driven ATP synthesis and mitochondrial ROS generation.

**Figure 4.**
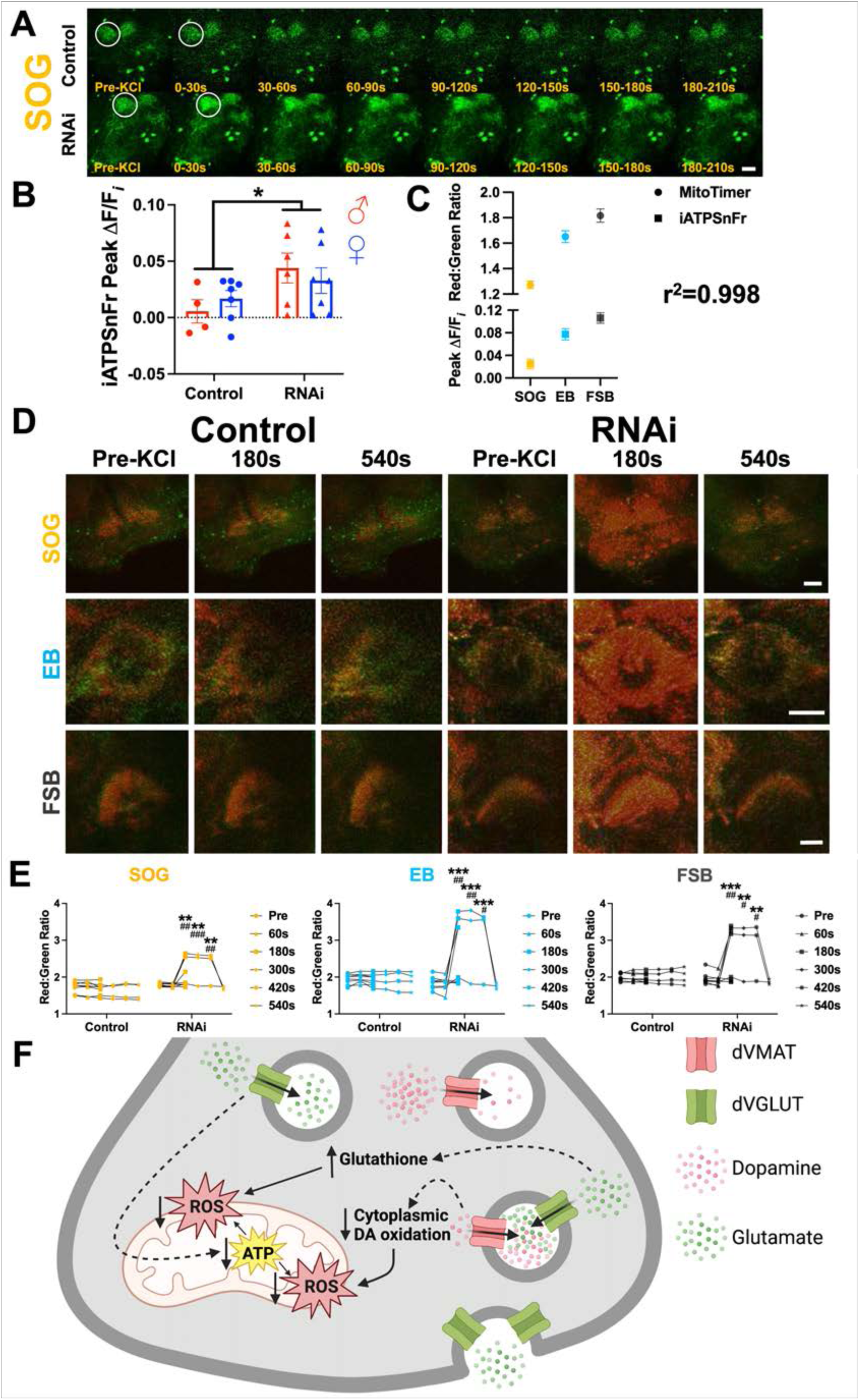
dVGLUT modulates activity-driven intracellular ATP and mitochondrial ROS generation in DA neurons. **(A)** Representative images of TH-driven iATPSnFr fluorescence (F) in DAergic projections to the SOG of adult (14d post-eclosion) *LexA* RNAi (Control, full genotype: UAS>iATPSnFr/w^1118^;TH>GAL4/UAS>LexA-RNAi) and *dVGLUT* RNAi (RNAi, full genotype: UAS>iATPSnFr/w^1118^;TH>GAL4/UAS>dVGLUT-RNAi) flies before and after 40mM KCl stimulation. White circles highlight regions that feature differences in intracellular DA neuron ATP levels during KCl-induced depolarization. Scale bar=25μm. **(B)** Quantification showed increased peak iATPSnFr fluorescence (Peak ΔF/F*_i_*) in SOG DAergic projections of RNAi flies (main effect of RNAi: F_1,20_=5.93, p=0.024, two-way ANOVA). N=4-7 per group, *p<0.05. Additional analyses are shown in Figure S5. **(C)** Regional differences in DA neuron baseline MitoTimer red:green ratio and iATPSnFr peak ΔF/F after KCl-induced depolarization were strongly correlated (r^2^=0.998, p=0.034). N=4 genotype/sex groups compared for correlation analysis. **(D)** Representative images of TH-driven MitoTimer labeling in DAergic projections to the SOG, EB, and FSB of Control and RNAi flies before and after 40mM KCl treatment. Scale bars=25μm. **(E)** DA neuron dVGLUT RNAi knockdown increased mitochondrial ROS acutely in response to 40mM KCl stimulation compared to Control flies (KCl × RNAi interaction, SOG: F_5,45_=6.38, p<0.001; EB: F_5,43_=6.63, p<0.001; FSB: F_5,43_=6.08, p<0.001, two-way ANOVA). **p<0.01, ***p<0.001 compared to baseline (pre-KCl), ^#^p<0.05, ^##^p<0.01, ^###^p<0.001 compared to Control flies (Bonferroni multiple comparisons test). N=8-9 per group. **(F)** Schematic showing that in DA neurons, dVGLUT decreases ATP levels during depolarization and decreases mitochondrial ROS both during depolarization and in response to paraquat. dVGLUT’s modulation of intracellular ATP and mitochondrial ROS accumulation may be due to its impacts on: 1) cytoplasmic glutamate availability, enabling efficient glutathione production to prevent toxic ROS accumulation, and 2) reducing levels of cytoplasmic DA available for oxidation that contribute to ROS generation. All data are plotted as Mean±SEM.

We next examined the impact of KCl-induced depolarization on dVGLUT-mediated DA neuron mitochondrial ROS in our *ex vivo* whole brain preparations. DA neuron dVGLUT RNAi dramatically increased MitoTimer red:green ratio during depolarization in the SOG (p<0.001), EB (p<0.001), and FSB (p<0.001; Figures 4D and 4E). In contrast, there was no discernable effect of KCl in controls across all assayed brain regions (p>0.05). Interestingly, we also observed a rapid return to baseline within ∼9 min of KCl treatment in the DA neuron dVGLUT RNAi flies, suggesting that mitochondrial ROS could still be cleared in the DA neurons, albeit in a diminished or altered manner in response to decreased dVGLUT expression (Figure 4E). Together, our findings demonstrate a novel role for dVGLUT in DA neuron resilience by mediating mitochondrial oxidative stress. We propose a model where dVGLUT decreases DA neuron mitochondrial ROS production either during heightened cell activity that places greater stress on the mitochondrial machinery of ATP generation (*i.e*., respiratory complexes) or following exposure to mitochondrial stressors like paraquat and H_2_O_2_ (Figure 4F).

### RNAseq reveals differentially expressed genes in VGLUT-expressing DA neurons conserved across flies and mice

To further dissect the mechanisms by which VGLUT confers resilience to DA neurons, we identified genes that are differentially expressed (DE) in TH^+^/VGLUT^+^ DA neurons relative to VGLUT^−^ DA neurons. Since VGLUT’s neuroprotective properties are shared between *Drosophila* and mammals, we employed a comparative approach to identify conserved DE genes in VGLUT-expressing DA neurons. In flies, we analyzed our recent 10x single-cell RNA sequencing (scRNAseq) of the fly DA system in adult males and females^69^. scRNAseq identified a total pool of 2,642 monoaminergic neurons that express the *Drosophila* ortholog of the vesicular monoamine transporter (*dVMAT*). Among dVMAT^+^ cells, we found 1,727 DA neurons based on co-expression of the dopamine transporter (*DAT*) (Figure 5A). Annotation of dVMAT^+^/DAT^+^/TH^+^ DA neurons via Uniform Manifold Approximation and Projection (UMAP) plots revealed multiple subclusters of dVGLUT^+^ DAergic cells, consistent with RNAscope findings described above (Figure 1) and previously^48^.

**Figure 5.**
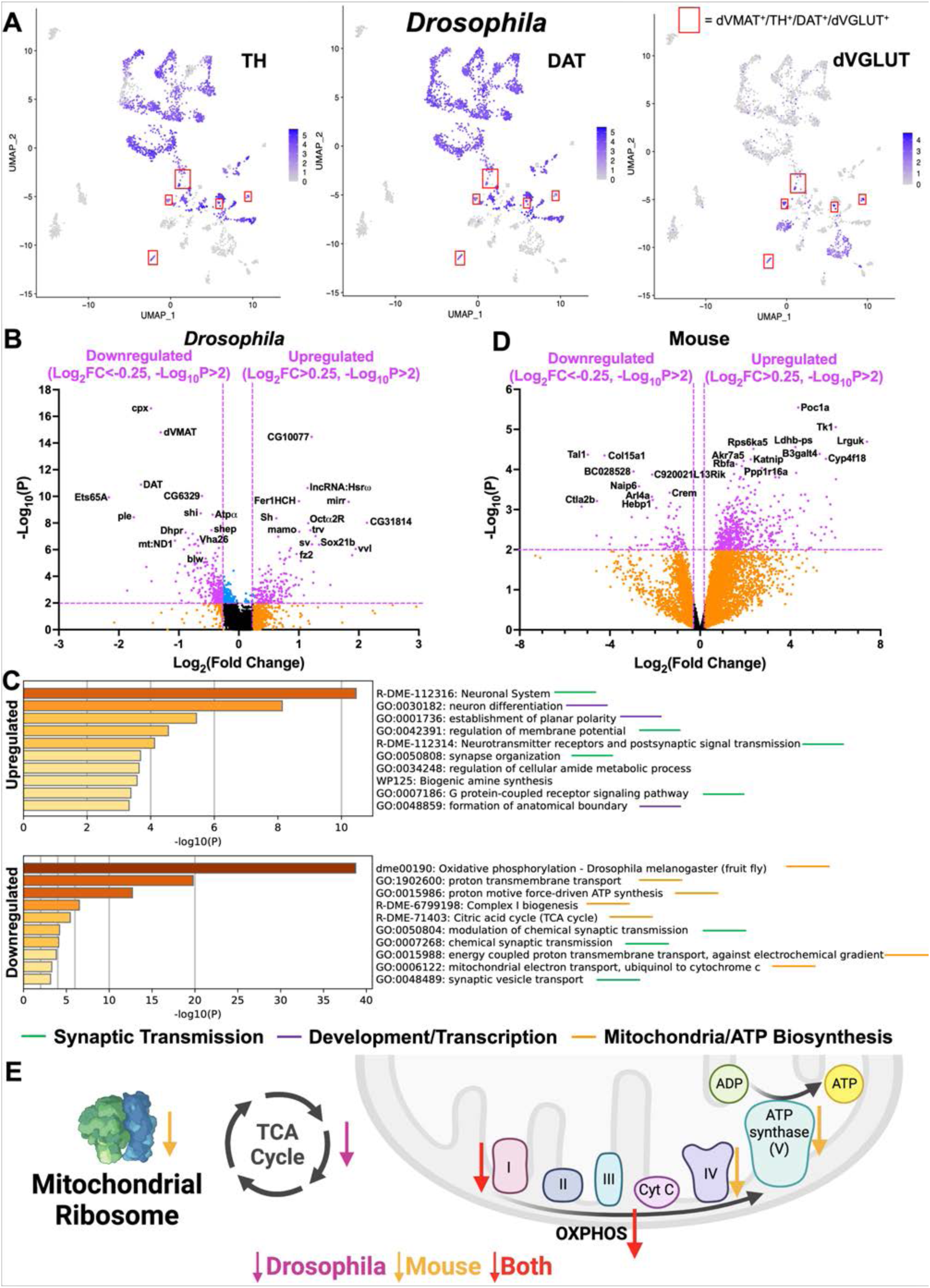
RNAseq reveals differentially expressed genes in VGLUT-expressing DA neurons conserved between flies and mice. **(A)** Annotations of adult *Drosophila* brain 10X single-cell RNAseq (scRNAseq) data via Uniform Manifold Approximation and Projection (UMAP) plots identified discrete clusters of dVMAT^+^ DA neurons. Expression of *TH*, *DAT* and *dVGLUT* across all 2,642 dVMAT-expressing cells is represented, with highest expression shown in purple and lowest/no expression shown in gray. Specific subpopulations of dVMAT^+^/TH^+^/DAT^+^/dVGLUT^+^ cells are highlighted by red boxes. **(B)** Volcano plot showing differential expression of genes in dVGLUT^+^ DA neurons compared to dVGLUT^−^ DA neurons. **(C)** Gene ontology (GO) analysis of *Drosophila* scRNAseq data identifying top pathways based on differential gene expression in dVGLUT^+^ DA neurons. **(D)** Volcano plot showing differential expression of genes in mouse VGLUT2^+^ DA neurons compared to all DAT^+^ DA neurons. Additional analyses relating to VGLUT2^+^ DA neurons are shown in Figures S6 and S7 and Videos S1 and S2. **(E)** Schematic summarizing the mitochondrial findings from the comparative fly and mouse RNAseq studies. These data revealed overall downregulation of genes implicated in mitochondrial processes in both *Drosophila* dVGLUT^+^ and mouse VGLUT2^+^ DA neurons. This included downregulation of mitochondrial ribosomal genes in mouse and overall downregulation of TCA cycle genes in flies. Among the mitochondrial respiratory complexes in VGLUT2^+^ DA, Complex I gene expression was downregulated in both mice and flies, while Complex IV and ATP synthase gene expression were specifically downregulated in mice.

DE gene analyses showed 160 genes were significantly upregulated in dVGLUT^+^ DA neurons [log_2_ fold change (FC)>0.25, -log_10_(P)>2] and 263 genes that were downregulated [log_2_FC<-0.25, -log_10_(P)>2] in dVGLUT^+^ DA neurons (Figure 5B and Table S2). Gene ontology (GO) pathway analysis revealed that the top gene categories associated with DE genes in dVGLUT^+^ DA neurons were development/gene transcription, synaptic neurotransmission, and mitochondrial metabolism/ATP biosynthesis (Figure 5C). Upregulated genes included several encoding transcription factors implicated in nervous system development and patterning such as *mirror* (*mirr*), *ventral veins lacking* (*vvl*), *Sox21b,* and *shaven* (*sv*). Other upregulated genes were implicated in synapse development and remodeling like the Wnt receptor *frizzled 2* (*fz2*), or in neurotransmission such as the α2 octopamine receptor (*Octα2R*). Significantly, many of the genes downregulated in dVGLUT^+^ DA neurons were associated with mitochondrial function and ATP production such as *mt:ND1* and *bellwether* (*blw*). Interestingly, the machinery of DA biosynthesis and packaging was also downregulated in dVGLUT^+^ DA neurons compared to dVGLUT^−^ DA neurons, including *TH* (*ple*), *dVMAT*, and *DAT* (Figure 5B). Since dVGLUT^+^ DA neurons possess enhanced activity-dependent vesicular DA loading and release^70^, decreased expression of DA biosynthesis/packaging and bioenergetics may represent a homeostatic response to prevent over-secretion of DA as well as to mitigate DA-driven ROS generation during periods of heightened activity. Moreover, we found greater upregulation of synaptic transmission pathway genes in male flies and stronger downregulation of oxidative phosphorylation pathway genes in females (Figure S6A). Based on our data connecting mitochondrial genes and function to dVGLUT^+^ DA neurons, we conducted additional analyses using MitoCarta^71^ to identify mitochondria-specific pathways. We found that genes involved in oxidative phosphorylation and mitochondrial respiratory complex I were significantly downregulated in dVGLUT^+^ DA neurons, and this downregulation was stronger in females (Figures S6B and S6C). These results suggest that dVGLUT^+^ DA neurons demonstrate a distinct transcriptional profile compared to dVGLUT^−^ DA neurons, including downregulated expression of genes implicated in mitochondrial function and ATP synthesis, mirroring the decreased mitochondrial ROS accumulation and ATP production observed in dVGLUT^+^ DAergic projections in the SOG.

To translate our *Drosophila* RNAseq findings to mammals, we employed an intersectional genetic strategy to selectively label VGLUT2-expressing DA neurons (TH^+^/VGLUT2^+^) in mouse midbrain for RNAseq (see Methods). Using the INTRSECT2.0 system^72^, we virally labeled TH^+^/VGLUT2^+^ cells with EYFP alongside mCherry-labeled TH^+^/VGLUT2^−^ DA neurons. We developed a workflow for accurately identifying labeled cell bodies and their projections throughout whole cleared mouse brains imaged by ribbon scanning confocal microscopy (RSCM) (see Methods). Following assembly and rendering of the 3D RSCM volumetric data, we aligned the whole brain volumes to the Allen Mouse Brain Atlas (CCFV3)^73^. Cell bodies were detected and classified using a neural network-based approach^74^; cell projections were identified following binarization^75^. We found EYFP-labelled TH^+^/VGLUT2^+^ cell bodies in the medial VTA and lateral SNc. Further, we observed local projections in the midbrain alongside projections to the striatum, consistent with recent reports that TH^+^/VGLUT2^+^ DA neurons project to the medial nucleus accumbens shell and tail of striatum^23,76^. We also observed TH^+^/VGLUT2^+^ DA neuron projections to the isocortex and hippocampal formation (Figure S7, Movie S1 and Movie S2), which is also in line with previous findings^23,76^. By comparison, there were substantially more mCherry-labeled TH^+^/VGLUT2^−^ cell bodies in midbrain with extensive projections throughout striatum, hippocampus, and isocortex (Figure S7, Movie S1 and Movie S2).

We next isolated EYFP-labeled TH^+^/VGLUT2^+^ neurons from midbrains of INTRSECT2.0-labeled brains for sequencing via bulk RNAseq. To maximize the number of available cells for comparison, we conducted bulk RNAseq of the total midbrain DAT^+^ DA neuron population in DAT-Cre::TdTomato mice (see Methods). Altogether, we found 516 genes upregulated and 84 genes downregulated in mouse VGLUT2^+^ DA neurons compared to all DAT^+^ DA neurons (Figure 5D and Table S3). GO pathway analyses of the mouse RNAseq data revealed overrepresentation of genes in TH^+^/VGLUT2^+^ neurons associated with mitochondrial metabolism/ATP biosynthesis, development/gene transcription, and autophagy/apoptosis (Figure S6D). As in the fly, *VMAT2* (encoded by the *Slc18a2* gene) expression was downregulated in mouse TH^+^/VGLUT2^+^ neurons (Table S3). Mitochondria-specific pathway analysis in mouse revealed strong downregulation of genes involved in mitochondrial function as in *Drosophila* (Figure S6E). This included downregulation of respiratory complex I and oxidative phosphorylation genes (Figures S6E, Table S3), suggesting an important role for mitochondrial function in dVGLUT/VGLUT2-associated DA neuron resilience (Figure 5E). Further, the greater downregulation of mitochondrial gene expression in female fly dVGLUT^+^ DA neurons compared to males raises the question of whether these differences contribute to or even drive VGLUT-mediated sex differences in DA neuron resilience.

### Forward genetic screens identify modulators of VGLUT-associated DA neuron resilience

We conducted a forward genetic RNAi screen in *Drosophila* to identify which of the DE genes identified in our fly and mouse RNAseq data mediate sexually dimorphic DA neuron resilience. Starting with 423 DE genes from our fly TH^+^/dVGLUT^+^ neuron RNAseq dataset, we narrowed our list to 187 gene candidates for screening (see Methods, Figure 6A and Table S4). We focused on DE gene candidates with 1) known roles in DA neurons and/or PD, 2) the greatest expression differences between dVGLUT^+^ and dVGLUT^−^ DA neurons, and 3) the strongest sex differences in expression. We removed candidates that did not possess mammalian gene orthologs or did not have appropriate publicly available RNAi lines (Table S4, see Methods). Finally, we included all significant DE genes that were conserved across fly and mouse RNAseq datasets, as well as fly orthologs of mouse transcription factors identified as upstream regulators of DE transcripts (Figures 6A and S6F).

**Figure 6.**
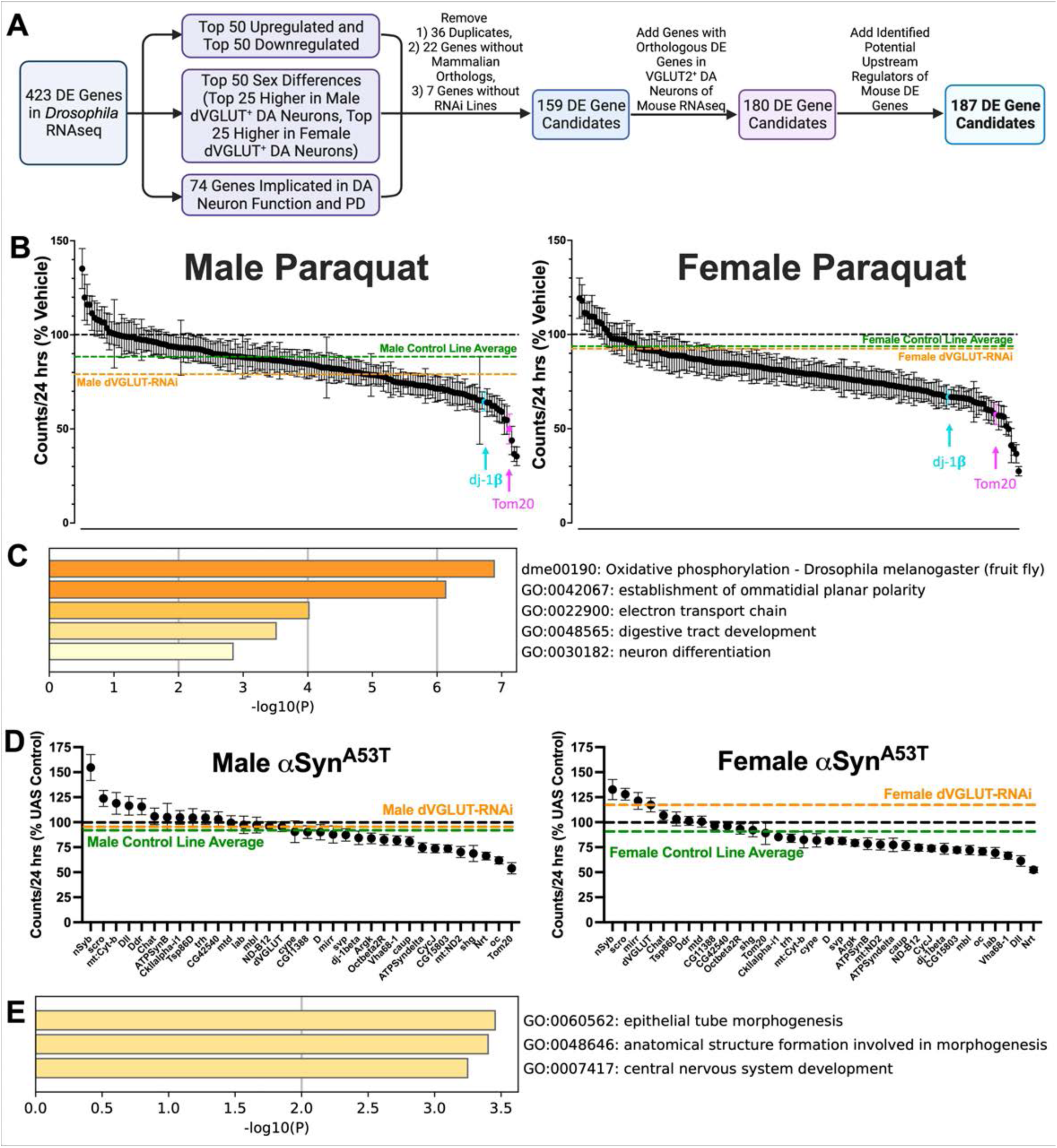
Forward genetic screens identify modulators of VGLUT-associated DA neuron resilience. **(A)** Flow chart depicting the process by which candidate genes were selected for forward genetic screening. **(B) Primary paraquat screen:** 187 candidate genes differentially expressed in TH^+^/VGLUT^+^ DA neurons were knocked down via RNAi. Adult (2d post-eclosion) male and female TH-driven RNAi flies were exposed to either paraquat (10mM, 4d) or vehicle control. Paraquat-induced alterations in locomotion over days 2-4 of exposure are plotted as % locomotion relative to each candidate’s vehicle-treated control. For reference, TH-driven dVGLUT RNAi flies (orange dashed lines) were tested alongside candidates. Average paraquat locomotion of 5 Bloomington *Drosophila* Stock Center (BDSC)/Harvard Transgenic RNAi Project (TRiP) control lines tested are represented by green dashed lines. Two screen hits that are orthologs of human genes with known roles in PD are highlighted (*i.e.*, *dj-1β*, *Tom20*). N=3-37 flies per group; mean N=18 flies per group. **(C)** GO pathway analysis of paraquat screen candidate hits. **(D) Secondary αSyn^A53T^ screen:** The top 32 candidate genes from the primary screen were knocked down via TH-driven RNAi alongside TH-driven αSyn^A53T^ expression. αSyn^A53T^-induced changes in locomotion (at 4-6d post-eclosion) are plotted as % UAS Control. N=12-39 flies per group; mean N=21 flies per group. **(E)** GO pathway analysis of candidate hits from the αSyn^A53T^ secondary screen. Each candidate’s results in the screens along with control line results are shown in Table S5. Further analyses of candidate genes are included in Figure S8. All data are plotted as Mean±SEM.

We established our primary screen to determine whether candidates modified locomotor response to paraquat-induced insult in adult male and female flies (Figure 6B). We obtained RNAi lines for all candidates from either the Bloomington *Drosophila* Stock Center (BDSC)/Harvard Transgenic RNAi Project (TRiP)^77^ or Vienna Drosophila Resource Center (VDRC). These lines were crossed with TH>GAL4 flies to create DA neuron-specific candidate knockdown (Table S4). We measured locomotion in these flies during early paraquat exposure (days 2-4) – a period when locomotion is affected but prior to significant DA neuron loss (see Figure 2). This enabled us to identify candidates during a critical window of DA neuron vulnerability to neurotoxicant stress.

We first validated our findings demonstrating increased vulnerability of male TH-driven dVGLUT RNAi flies to paraquat (see Figure 2E) using the smaller sample sizes necessary for a higher-throughput screen. We found that ∼16 flies/group/sex were sufficient to detect increased locomotor vulnerability to paraquat (Figure S8A). We also confirmed greater locomotor vulnerability in male control flies to paraquat compared to females (Figures 6B, S8B). In response to TH-driven RNAi of the 187 candidates, 11 lines were lethal or had significant lifespan deficits (*e.g.*, *Vha26*, *mamo*), precluding further analysis (Table S4). We found a normal distribution of paraquat’s effects on locomotion among the remaining candidates, including some candidates which exhibited greater decreases in locomotion compared to either control or dVGLUT RNAi lines (Figure 6B, Table S5). Importantly, we discovered that DA neuron knockdown of *dj-1β* and *Tom20* produced some of the strongest paraquat-induced locomotor impairments in both males and females. Interestingly, the human orthologs of *dj-1β* and *Tom20*, *DJ-1* (*PARK7*) and *TOM20* respectively, have documented roles in PD pathogenesis^78,79^. These results validated our screen as an effective means of identifying modulators of DA neuron vulnerability relevant to PD. Importantly, our data provide a possible cellular locus of VGLUT^+^ DA neurons for human PD pathology.

To determine screen hits, we adjusted for differences in locomotion and vulnerability of female flies attributable to the source of RNAi vector [BDSC/TRiP lines versus VDRC lines] (Figure S8C; see Methods). Post-correction, we selected all candidate genes that produced: 1) a significant effect of paraquat in both males and females, or 2) a significant sex difference in percent change in locomotion in response to paraquat (Table S5); significance was indicated by p<0.05 and a false-discovery rate (FDR) q<0.05. We identified 18 candidates shared between males and females (*dj-1β*, *Argk*, *oc*, *shg*, *CycJ*, *CkIIalpha-i1*, *ATPSynB*, *cype*, *svp*, *Tom20*, *trh*, *lab*, *Nrt*, *scro*, *CG42540*, *ATPsyndelta*, *Vha68-1*, *CG15803*) (Table S5). We also discovered 14 hits with sex differences in response to paraquat (Table S5). Knockdown of *mirr*, *Dd*r, *Tsp86D*, and *mt:ND2* elevated male vulnerability to paraquat-induced locomotor impairment compared to females (*i.e*., greater percent decrease in locomotion caused by paraquat treatment in males compared to females), while knockdown of *ChAT*, *nSyb*, *Dll*, *D*, *ND-B12*, *caup*, *Octbeta2R*, *mtd*, *mbl*, and *mt:Cyt-b* induced greater female vulnerability compared to males. Consistent with the RNAseq data, hits were associated with mitochondrial function (*dj-1β*, *Argk*, *ATPSynB*, *cype*, *Tom20*, *ATPsyndelta*, *ND-B12*, *mt:Cyt-b*, *mt:ND2*) and transcriptional regulation/development, particularly of the central nervous system (*oc*, *shg*, *CycJ*, *CkIIalpha-i1*, *svp*, *trh*, *lab*, *Nrt*, *scro*, *mirr*, *dll*, *D*, *caup*, *mbl*). This was confirmed by GO pathway analysis of the combined top candidates (Figure 6C).

To validate candidates from our initial screen, we developed a secondary screen based upon the human α-synuclein A53T (αSyn^A53T^) fly PD model. TH-driven overexpression of αSyn^A53T^ induces pathological α-synuclein aggregation which leads to early locomotor deficits and DA neurodegeneration^80,81^. When we tested the effect of DA neuron αSyn^A53T^ expression on age-related decreases in locomotion, we did not see an exacerbation of age-related locomotor degeneration in male or female controls (Figure S8D). Moreover, though dVGLUT RNAi males were more vulnerable to age-related locomotor impairment as previously described^36^, DA neuron αSyn^A53T^ overexpression did not exacerbate this reduction in locomotion (p>0.05; Figure S8D). In contrast, females co-expressing DA neuron dVGLUT RNAi and αSyn^A53T^ displayed a significant locomotor reduction compared to female dVGLUT RNAi flies without αSyn^A53T^ overexpression (p=0.021; Figure S8D). These results suggest a sex-specific interaction between dVGLUT and αSyn^A53T^ in mediating DA neuron vulnerability in females and indicate that the female DAergic system may require more “hits” than males to become vulnerable. Further, our data demonstrate the ability of the αSyn^A53T^ model to detect potential mediators of PD-related vulnerability.

For the secondary screen of the 32 top hits, we overexpressed αSyn^A53T^ alongside candidate gene RNAi in DA neurons and measured locomotion at 4-6d post-eclosion. We found 19 candidates (after correction for RNAi source; Figure S8E) with either a significant decrease in locomotion in both sexes or a significant sex difference in response to αSyn^A53T^ (Figure 6D). After removing lines with FDR q>0.05, 13 candidates remained: *Nrt*, *CycJ*, *CG15803*, *Argk*, *shg*, *mt:ND2*, *mt:Cyt-b*, *Tom20*, *oc*, *Dll*, *mirr*, *CkIIalpha-i1*, and *trh* (Table S5). Interestingly, while some candidates have roles in mitochondrial function (*mt:ND2*, *mt:Cyt-b*, *Tom20*), many have developmental roles, serving as transcription factors in central nervous system development (*oc*, *Dll*, *mirr*, *trh*) or mediating cell adhesion in development (*Shg*, *Nrt*) (Figures 6E, S8F).

### Top candidate transcription factors impact DA neuron numbers in a region- and sex-specific manner

Based on the primary and secondary screens, we selected 6 top candidates for further testing. We focused on *oc*, *Dll*, *mirr*, *trh*, and *caup*, since they are transcription factors, which were overrepresented among the screen’s top hits, and they display very distinct expression patterns in specific DA neuron clusters (Figure 7A). We additionally selected *dj-1β* given its established roles in mediating DA neuron vulnerability^82,83^. We discovered that TH-driven knockdown of *mirr*, *trh*, *Dll*, and *oc* significantly impacted DA neuron numbers in region- and sex-specific manners (Figures 7B, 7C, Table S6); there was no significant effect of DA neuron *caup* or *dj-1β* knockdown on DA neuron numbers (p>0.05). DA neuron-specific RNAi knockdown of either *mirr* or *Dll* diminished DA neuron numbers in some discrete DAergic cell clusters (PPM1/2) but increased neuron numbers in others (PAL, PPL2). Knockdown of DA neuron *trh* and *oc* also produced region-specific decreases in DA neuron numbers (Figures 7B, 7C). Strikingly, *oc* knockdown almost eliminated all DA neurons in the PPL2ab subcluster (p<0.001), indicating a critical role for *oc* among PPL2 DA neurons^84^. Altogether, these data provide evidence that transcription factors DE in TH^+^/VGLUT^+^ neurons play important roles in the regulation of DA neuron development and/or survival in select brain regions of adult males and females.

**Figure 7.**
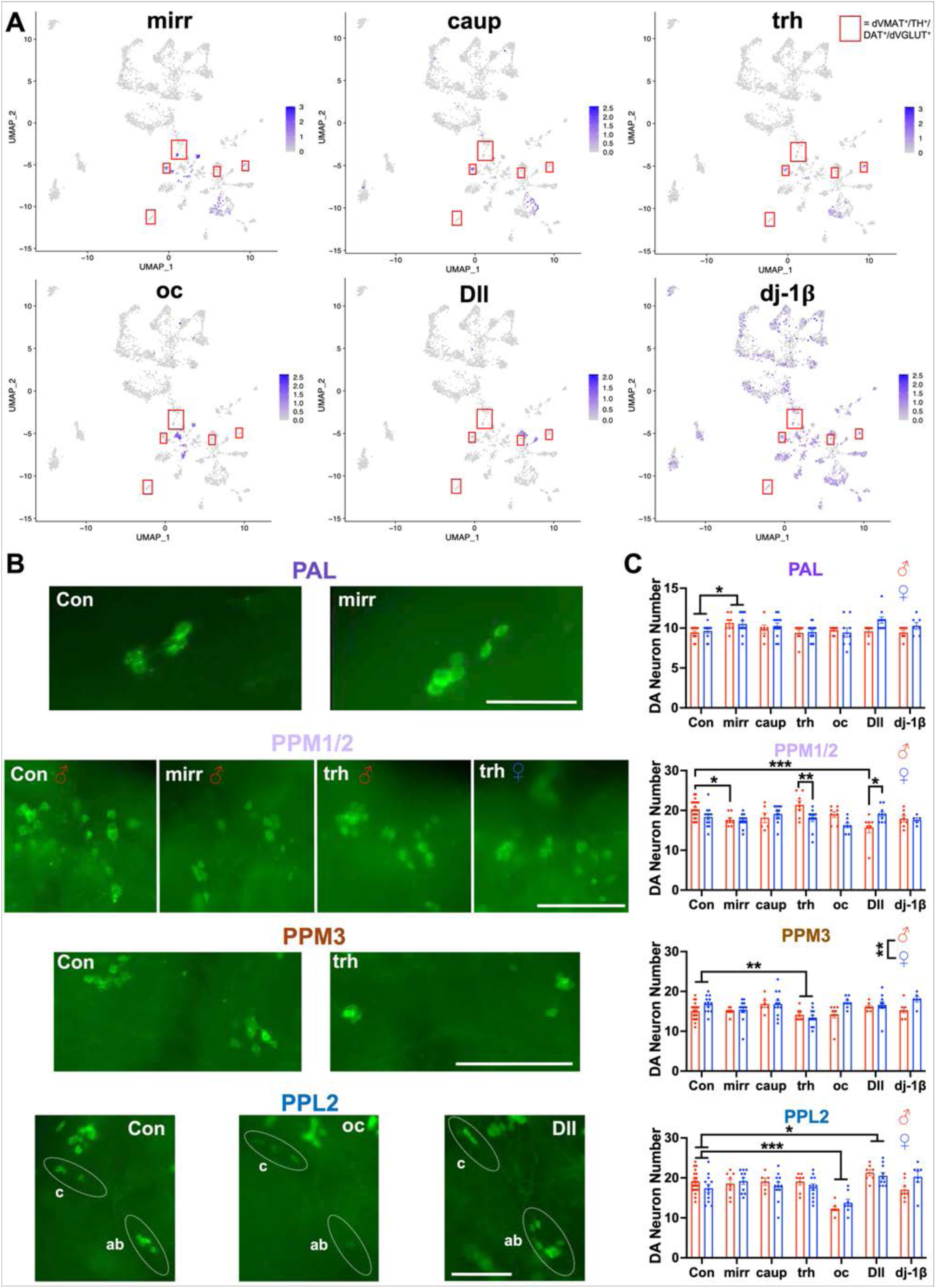
Top candidate transcription factors impact DA neuron numbers in a region- and sex-specific manner. **(A)** UMAP plots showing expression of selected top candidates from primary and secondary genetic screens in dVMAT-expressing cells. Highest expression levels are shown in purple and lowest/no expression is shown in gray. dVMAT^+^/TH^+^/DAT^+^/dVGLUT^+^ cells are highlighted by red boxes. **(B)** Representative images of GFP-labeled DA neurons demonstrating the impact of TH-driven RNAi knockdown of top gene candidates and *LexA* RNAi control (Con) on DA neuron number (14d post-eclosion) in the PAL, PPL2, PPM1/2 and PPM3 cell clusters. PAL, PPL2 scale bars=100μm, PPM1/2, PPM3 scale bars=250μm. **(C)** TH-driven RNAi of candidate genes *mirr*, *trh*, *oc*, and *Dll* differentially altered GFP-labeled DA neuron numbers in region-specific and sex-specific manners compared to controls. N=5-21 fly brains per group. *p<0.05, **p<0.01, ***p<0.001, two-way ANOVA with Bonferroni multiple comparisons test. All data are plotted as Mean±SEM.

### dj-1β modulates sex differences in DA neuron dVGLUT expression

Given the critical roles played by dj-1β in oxidative stress in PD^85^, and the upregulation of dVGLUT in response to paraquat-induced stress, we examined whether dj-1β modulates DA neuron dVGLUT expression with or without paraquat-induced oxidative stress. We performed RNAscope of green fluorescent protein (GFP)-labeled DA neurons to quantify the impact of *dj-1β* knockdown on *dVGLUT* mRNA expression in DA neurons of adult males and females. In control flies, higher percentages of TH^+^ DA neurons expressing dVGLUT were found in PPL1 and SOG DA neurons compared to other DAergic clusters (p=0.0041 for main effect of region; Figures 8A, 8B). Additionally, paraquat exposure across 10d did not significantly alter the proportion of DA neurons expressing dVGLUT in controls (p>0.05; Table S6). Importantly, in response to TH-driven *dj-1β* knockdown, we observed a significant increase in the proportion of dVGLUT-expressing DA neurons within the PPL2 of male flies (p=0.018); we observed no significant effect in other brain regions (p>0.05; Table S6). Females were not affected (p>0.05), and the increase was eliminated by paraquat exposure (p=0.016; Figures 8C, 8D). These results demonstrate an important relationship between dj-1β and dVGLUT in the overall mechanisms of resilience to DA neuron insults. We posit that the loss of dj-1β, a key chaperone responsible for modulating mitochondrial oxidative stress, induces compensatory dVGLUT upregulation within DA neurons, enabling continued DA neuron survival.

**Figure 8.**
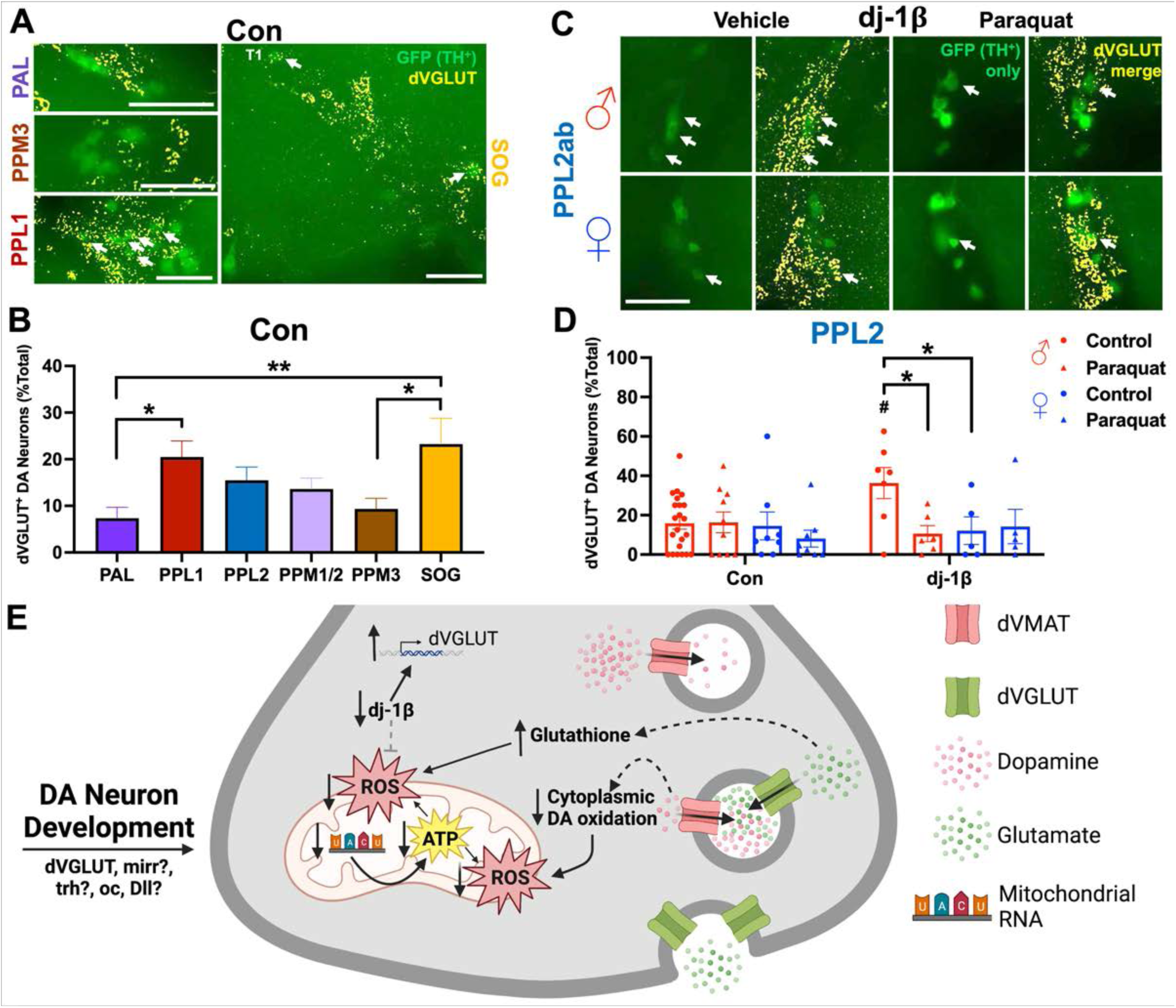
*dj-β* modulates sex differences in DA neuron dVGLUT expression. **(A)** Representative images of DA neuron *dVGLUT* mRNA expression (in yellow); TH^+^ DA neurons are GFP-labeled (in green) in adult *LexA* RNAi control (Con) flies. DA neuron dVGLUT expression is shown in the PPL1 and SOG (both T1 and lateral SOG dVGLUT-expressing DA neurons), incontrast to the PAL and PPM3 DA neuron clusters. Scale bars=100μm. **(B)** Quantification of DA neuron dVGLUT expression showed higher percentages of dVGLUT^+^ DA neurons in the PPL1 and SOG compared to other DA neuron clusters (F_5,168_=3.59, p=0.0041, one-way ANOVA). N=25-31 fly brains per group. **(C)** Representative images of DA neuron dVGLUT mRNA expression in response to TH-driven *dj-1β* RNAi (dj-1β); images focus on the PPL2ab cluster of GFP-labeled DA neurons from *dj*-*1β* RNAi flies exposed to 10mM paraquat (10d) or vehicle control. **(D)** Quantification in the PPL2 DA neuron cluster showed a paraquat × sex × genotype interaction (F_1,63_=4.29, p=0.043, three-way ANOVA) where TH-driven *dj-1β* knockdown in males boosted the percentage of DA neurons expressing dVGLUT compared to male Con flies (p=0.018, Bonferroni multiple comparisons test). N=5-22 fly brains per group. **(E)** Schematic summarizing mechanisms of dVGLUT-mediated DA neuron resilience. dVGLUT expression protects DA neurons by decreasing ATP production and mitochondrial ROS levels, which is reflected by lower mitochondrial gene expression in dVGLUT^+^ DA neurons. Mitochondrial oxidative stress vulnerability may also be lower in dVGLUT-expressing DA neurons due to increased glutathione and decreased cytoplasmic DA oxidation. Additionally, vulnerability may be impacted by differential expression of transcription factors that impact DA neuron development and/or differentiation. Finally, during periods of cell stress, DA neuron dVGLUT expression is raised as part of a neuroprotective response to accumulating mitochondrial ROS. This increase in dVGLUT expression is triggered by the concomitant loss of dj-1β during periods of cell stress/insult, particularly in males. *p<0.05, **p<0.01, ^#^p<0.05 compared to Con Male receiving Control food. All data are plotted as Mean±SEM.

## Discussion

In this study, we show, for the first time, fundamental mechanisms by which VGLUT confers selective resilience to DA neurons. We demonstrate that dVGLUT expression is dynamically upregulated in DA neurons in response to neurotoxicant exposure or aging. We also find that dVGLUT-expressing DA neurons display: 1) decreased mitochondrial gene expression, 2) decreased baseline mitochondrial ROS, and 3) decreased depolarization-induced cytoplasmic ATP levels. Consistent with this, DA neuron dVGLUT expression reduces mitochondrial ROS during cell stress and diminishes activity-dependent increases in intracellular ATP and mitochondrial ROS. Just as importantly, we identified sex differences, where males are more vulnerable to *dVGLUT* knockdown in DA neurons, and females display higher DA neuron dVGLUT expression along with greater mitochondrial resilience. Finally, by conducting a forward genetic screen, we identified novel modulators of DA neuron resilience and dVGLUT expression, particularly genes primarily associated with mitochondrial metabolism and transcriptional regulation. Among these genes, we discovered that *dj-1β*, a gene associated with familial early-onset PD and PD pathogenesis, is a sex-specific regulator of DA neuron dVGLUT expression. Collectively, our findings demonstrate new VGLUT-mediated mechanisms of DA neuron resilience in males and females.

As in the mammalian brain, there is increasing recognition of neurotransmitter co-transmission in *Drosophila*, including in neurons that co-release DA alongside glutamate^36,70,86–91^. In this study, we comprehensively mapped dVGLUT expression throughout the fly DA system for the first time. We confirmed the presence of dVGLUT^+^ DA neurons within the PPM1 and PPL1 brain regions^48^, which is consistent with work showing dVGLUT protein expression in DAergic PPL1 projections to the PPL1-γ2α’1 (MB-MV1) region^70^. We also identified previously undescribed dVGLUT-expressing DA neurons in the SOG and PPL2 clusters. The identification of such discrete subpopulations of dVGLUT^+^ DA neurons provides a unique model for the dissection of region- and cell-specific mechanisms underlying dVGLUT-mediated resilience. Additionally, while the adult fly brain does not possess anatomic structures identical to those found in the mammalian midbrain (*i.e.*, VTA and SNc), there are similarities in the organization of TH^+^/VGLUT^+^ DA neurons between flies and mammals. In both cases, VGLUT^+^ DA neurons are organized into several well-defined clusters that project to regions strongly implicated in positive and negative reinforcement as well as learning and memory^76,92–97^. We therefore posit that there may be evolutionarily conserved mechanisms underlying the development, organization, and/or survival of VGLUT^+^ DA neurons.

Our intersectional genetic dVGLUT reporter revealed upregulation of DA neuron dVGLUT in response to paraquat exposure, consistent with previously observed increases across aging^36^. Significantly, we and others also described dynamic DA neuron VGLUT2 upregulation and/or VGLUT2-associated resilience in 6-OHDA, MPTP, rotenone, and αSyn pre-formed fibril (PFF) preclinical PD models^8,33,35,98^. Likewise, the proportion of surviving SNc DA neurons expressing VGLUT2 was ∼3-fold higher in male PD patients compared to control subjects^35^. Overall, our results demonstrate for the first time in *Drosophila* PD models that dVGLUT upregulation is neuroprotective for DA neurons, highlighting the importance of regulatory mechanisms for VGLUT expression in DA neurons.

Aging is the greatest risk factor for neurodegenerative disorders including PD^99^, and we indeed observed stronger DA neurotoxicity of paraquat in aged flies. Importantly, dVGLUT upregulation was observed in DA neurons of aged flies in response to paraquat exposure. Moreover, the dVGLUT-expressing SOG DAergic projection region was resilient to age-related increases in mitochondrial ROS. While we did not observe age-related changes in mitochondrial ROS accumulation in SOG projections in response to dVGLUT RNAi knockdown, this may be due to residual dVGLUT expression exerting neuroprotective effects. Nevertheless, considering previous findings of dVGLUT RNAi effects on age-related DA neurodegeneration^36^, we contend that VGLUT functions as an important modulator of age-related mitochondrial dysfunction. Future studies should investigate the intersection between aging, PD model vulnerability, and VGLUT-mediated resilience.

We found significant sex differences in dVGLUT-associated DA neuron resilience. Specifically, we found higher DA neuron dVGLUT expression in females, and female flies were more resilient to paraquat, in line with previous work^36,46^. These sex differences fit with both preclinical PD models and clinical PD. PD affects more men than women and has an earlier age of onset for men^10–17^, and rodent PD model studies find less SNc DA neurodegeneration and milder motor symptoms in females^19–21^. Consistent with this, we showed decreased baseline mitochondrial oxidation in female DAergic synapses. Indeed, previous reports show higher baseline expression of antioxidant enzymes and associated protection from mitochondrial damage in females in various PD models^100–102^. We therefore hypothesize that sex differences in dVGLUT-mediated mitochondrial oxidative stress vulnerability underlie this sexual dimorphism. However, while females were resistant to dVGLUT RNAi-induced vulnerability to paraquat, DA neuron *dVGLUT* knockdown increased female vulnerability to αSyn^A53T^ overexpression, highlighting that *dVGLUT* mediates female resilience to some models of DA neurodegeneration^36^. Though the mechanisms behind sex differences in PD are unknown, there are several potential candidates with sexually dimorphic expression that may interact with VGLUT to protect DA neurons in PD. In mammals, estrogen reduces mitochondrial oxidative stress in PD models^22^. Conversely, the *sex determining region of the Y chromosome* (*SRY*) is upregulated in both animal and cell PD models, and its inhibition confers DA neuroprotection^103^. We propose that these sex-specific factors may impact DA neuron VGLUT2 expression and resilience. Further work is needed to investigate the impacts of estrogen and *SRY* in the context of VGLUT2-mediated DA neuroprotection in PD.

We found that DA neuron *dVGLUT* knockdown exacerbates paraquat-induced mitochondrial oxidative stress in DAergic projection regions specifically in males. Importantly, this increase in oxidative stress occurs at a stage of paraquat exposure that does not induce DA neuron degeneration. This suggests that dVGLUT exerts a protective role early in the neurodegenerative process, which is important for the creation of successful PD interventions. Relatedly, we found that dVGLUT RNAi increases both DA neuron ATP levels and DA neuron mitochondrial oxidation during depolarization. ATP production leads to ROS accumulation in axonal mitochondria^65^, consistent with our time-course of ATP and ROS accumulation in DAergic terminals in response to depolarization. dVGLUT may directly impact mitochondrial ATP production, which in turn increases vulnerability to oxidative stress. Interestingly, the increased depolarization-dependent DA neuron ROS accumulation in flies with *dVGLUT* knockdown was transient. Since *dVGLUT* knockdown was not total, this suggests that residual dVGLUT and/or other ROS-mediating factors may require more time to diminish activity-induced increases in DA neuron mitochondrial ROS compared to control flies. Further, we observed a regional correlation between baseline DA neuron mitochondrial oxidative stress and ATP during depolarization. DAergic projections to the SOG, which contain dVGLUT^+^ inputs from local SOG and PPM1 DA neurons^55–57^, had the lowest mitochondrial oxidation and ATP levels. These data are consistent with mammals, as resilient VTA DA neurons (which have higher levels of VGLUT2 co-expression), have not only greater resilience to PD and mitochondrial toxin models of PD, but also 1) lower mitochondrial density, 2) lower baseline mitochondrial ROS, and 3) lower baseline oxidative phosphorylation/ATP production^104^. However, it is important to note that other factors, like calbindin, also differ in expression between selectively vulnerable versus resilient midbrain DA neuron subregions^23,105^. We therefore posit that calbindin may additionally contribute to regional differences in mitochondrial vulnerability to oxidative stress via its calcium buffering capacity^106–108^.

How might dVGLUT decrease ATP burden during depolarization in DA neurons? Recent work has shown that VGLUT anion channel activity decreases ATP consumption by accelerating vesicular glutamate accumulation and reducing ATP-driven H^+^ transport^109^. This may be relevant to DA neurons given our previous work demonstrating VGLUT’s key roles in activity-dependent synaptic vesicle pH regulation^70^. During depolarization, VGLUT mediates hyperacidification by enhancing glutamate and chloride transport. Such activity-induced hyperacidification is driven by the vesicular Vacuolar-type ATPase (V-ATPase) which consumes more ATP to transport more H^+^ into the vesicle lumen, lowering the vesicular pH. This intraluminal hyperacidification facilitates the tuning of vesicular DA content^70^, since the proton gradient (Λ1pH) provides the main driving force for synaptic vesicle DA loading and retention^110^. Conversely, by decreasing VGLUT expression in DA neurons, we postulate that there is a diminished conduit for vesicular anion flux which would force the V-ATPase to work harder to maintain vesicular pH. The resulting increase in rates of ATP hydrolysis by the V-ATPase would elevate ATP demand and raise the overall metabolic burden of DA neuron mitochondria. Such a scenario would explain the higher levels of cytoplasmic ATP and mitochondrial ROS observed in response to DA neuron *dVGLUT* knockdown.

We conducted scRNAseq in *Drosophila* and bulk RNAseq in mice to further define the mechanisms of VGLUT-mediated DA neuron resilience. Specifically, we aimed to identify additional genes that function in tandem with VGLUT to protect DA neurons during periods of stress or injury. Our comparative sequencing approaches revealed numerous DE genes conserved across both fly and mouse VGLUT^+^ DA neurons, suggesting that the expression profiles of these cells are evolutionarily conserved. Notably, we further validated our fly scRNAseq data against a recent scRNAseq study of adult *Drosophila* DA neuron clusters which independently confirmed our DE genes in dVGLUT^+^ DA neurons and the directionality of their expression relative to dVGLUT^−^ DAergic cells (*i.e*., upregulated versus downregulated genes)^48^. Strikingly, in both datasets, mitochondrial genes, particularly genes implicated in respiratory complex I function/structure and oxidative phosphorylation, were downregulated in VGLUT-expressing DA neurons. This is in line with our findings showing the SOG, a dVGLUT-expressing DAergic projection region, had less mitochondrial ROS and lower ATP levels during activity compared to other regions. These measures were increased in response to DA neuron dVGLUT knockdown, giving further credence to a mitochondrial mechanism of VGLUT-mediated DA neuron resilience. Our data are also consistent with findings in mammalian brain showing increased mitochondrial density in vulnerable SNc DA neurons compared to resilient (and VGLUT2^+^) VTA DA neurons^104^. We hypothesize that diminished expression of genes associated with mitochondrial metabolism in VGLUT^+^ DA neurons is related to these cells’ ability to fire at higher frequencies (>20 Hz) compared to other DAergic cells^111–113^. Therefore, the ability to tightly regulate mitochondrial metabolism at the gene expression level may prevent the generation of toxic ROS in response to elevated neuronal firing rates. Just as importantly, downregulating mitochondrial genes during periods of stress or injury when energy demand is raised may further shield mitochondria from oxidative stress, increasing the likelihood of cell survival. Consistent with this, downregulation of respiratory complex I subunits confers protection against cisplatin-mediated ROS neurotoxicity^114^. Interestingly, we also discovered sex differences in expression of genes associated with oxidative phosphorylation and mitochondrial function between male and female flies, which aligns with evidence of sex differences in mitochondrial structure and function between males and females^115^. Further work is clearly needed to identify the mechanisms responsible for these sex differences at the mitochondrial level as this may provide additional targets for boosting DA neuron resilience.

We conducted a forward genetic screen of 187 genes identified from our RNAseq data to identify novel modulators of dVGLUT-associated DA neuron vulnerability in paraquat and αSyn^A53T^ PD models. Consistent with the RNAseq data, top gene hits were implicated in mitochondrial function and transcriptional regulation. Since we sought to identify modulators of dVGLUT-associated DA neuron vulnerability, we focused on transcription factors that may dynamically regulate dVGLUT expression during stress. We selected transcription factors with exclusive expression in dVGLUT^+^ DA neuron subclusters; we avoided candidates with broader expression patterns to better isolate effects on DA neuron vulnerability that were more likely to be dVGLUT-mediated. Importantly, 4 out of 5 of the screened transcription factors, *mirr*, *trh*, *oc*, and *Dll,* impacted DA neuron number upon knockdown, indicating regulation of DA neuron development and/or survival. *mirr* is a *Drosophila* ortholog of Iroquois homeobox transcription factors, which are expressed during mouse DA neuron development^116^, downregulated in rat VTA dVGLUT^−^ DA neurons^117^, and upregulated in PD patient pluripotent stem cell-derived DA neurons^118^. *trh* is the *Drosophila* ortholog of neuronal PAS domain protein 1 (*NPAS1*) and *NPAS3* in mammals. *NPAS3* expression is downregulated in DA neurons during early stages of PD^119^, while *NPAS1* represses *TH* expression during development^120^. *Dll* is an ortholog of NK-like distal-less homeobox transcription factors closely related to Hox-like genes. In *C. elegans*, *Dll* ortholog *ceh-43* controls both induction and maintenance of terminal DAergic fate^121^, while mouse *Dll* orthologs, *Dlx5/6*, regulate development and function of hypothalamic DAergic neurons^122^. Finally, *oc* is the fly ortholog of *Otx2*, which plays critical roles in mammalian DA neuron development^123^. Analogous to *oc*’s impact on fly PPL2 DA neurons, *Otx2* is required for mammalian VTA DA neuron development^124–126^. *Otx2* also shows decreased expression with age^127^ and tau deficiency^128^, and confers DA neuron protection from PD models^126,129,130^. Importantly, a subtype of VGLUT2^+^ DA neurons in the medial VTA expresses *Otx2*^23^. Based on these findings, we hypothesize that *dVGLUT/VGLUT2* and *oc/Otx2* are likely expressed together during development and work together to protect VTA DA neurons from PD. The precise relationship between *oc* and *dVGLUT* in DA neuron development will be tested in future studies.

In parallel to our transcription factor studies, we found that in the PPL2 DA neuron cluster, TH-driven knockdown of *dj-1β* increased DA neuron *dVGLUT* expression, and this effect was especially evident in male flies. Similar to our observations with *dVGLUT*, *DJ-1* (*PARK7*), the mammalian ortholog of *dj-1β,* mediates paraquat’s effect on mitochondrial oxidative stress^131^ as well as on respiratory complex I activity^85^. This observation raises the question: how does *DJ-1* impact DA neuron *dVGLU*T expression? DJ-1 is a molecular chaperone that modulates aggregation of proteins associated with neurodegeneration, including αSyn and tau^132,133^, and loss-of-function mutations are associated with early-onset familial PD^78,134^. Likewise, loss of DJ-1 in flies and mammals leads to mitochondrial stress and dysfunction while elevating ROS generation^131,135,136^. We therefore speculate that the loss of DA neuron *dj-1β* expression induces mitochondrial oxidative stress, which triggers dVGLUT upregulation as part of a compensatory neuroprotective mechanism. At present, we cannot rule out that *dj-1β* directly impacts *dVGLUT* transcription, since DJ-1 additionally functions as a regulator of gene transcription, including of inflammatory cytokines to reduce neuroinflammation in DA neurons^137^. It is therefore possible that *dj-1β*-induced transcriptional modulation may also impact dVGLUT transcription either directly or indirectly.

Based on our findings, we present multiple lines of evidence demonstrating a mitochondrial mechanism of dVGLUT-mediated DA neuron resilience (Figure 8E). In DA neurons expressing dVGLUT, mitochondrial gene expression is downregulated, and this expression is lower in females, which correlates with increased DA neuron dVGLUT expression in females. This decreased mitochondrial gene expression in dVGLUT-expressing DA neurons is associated with lower ATP levels during depolarization, lower baseline mitochondrial ROS, and greater female resilience to paraquat-induced mitochondrial ROS. We show that dVGLUT protects DA neurons by directly impacting baseline ATP production during depolarization and depolarization- and toxin-induced mitochondrial ROS accumulation. Further, DA neuron *dVGLUT* expression is increased in response to *dj-1β* knockdown. This may occur to compensate for ROS accumulation that is observed in DJ-1 depletion models^135^ and thus explain why VGLUT2 expression is associated with DA neuron survival in human PD patients^35^. We identified transcription factors DE in dVGLUT-expressing DA neurons that mediate DA neuron development and differentiation. The combination of these genes and dVGLUT may explain differences in mitochondrial gene expression and vulnerability between DA neuron subtypes. Finally, the capacity of VGLUT-expressing DA neurons to handle glutamate may offer additional neuroprotection by enhancing glutathione biosynthesis^138^ and reducing cytoplasmic DA oxidation^139^. Together, we demonstrate a comprehensive mechanism whereby dVGLUT protects DA neurons from the paraquat model of PD by decreasing mitochondrial ATP and ROS production, thereby improving mitochondrial health. Ultimately, manipulating VGLUT2 may offer a novel, effective therapeutic target in PD for both men and women.

## Methods

### Drug treatments

All drugs were purchased from Sigma-Aldrich (St. Louis, MO). Drugs were diluted to their final respective concentrations in molten cornmeal-molasses media: paraquat dichloride (10mM), hydrogen peroxide (H_2_O_2_, 3%), 4-hydroxy-2,2,6,6-tetramethylpiperidin-1-oxyl (TEMPOL; 3mM). Food containing drug or vehicle controls were then poured into standard fly vials and given 1 hour (h) to solidify prior to addition of flies.

### Drosophila studies

#### Drosophila strains

All *Drosophila melanogaster* strains were maintained on standard cornmeal-molasses media at 24°C, 60-70% humidity under a 12h light:12h dark cycle in humidity- and temperature-controlled incubators. Unless otherwise noted, fly stocks were obtained from the Bloomington *Drosophila* Stock Center (BDSC). Expression drivers included TH>GAL4^140^ (gift of Dr. Serge Birman, Université Aix-Marseille, Marseille, France), TH>LexA^141^, dVGlut>GAL4 (BDSC #60312), and dVGlut>LexA (BDSC #84442)^142,143^. For RNAi-mediated *dVGLUT* knockdown, we employed UAS>Vglut-RNAi^HMS^ (abbreviated dVGLUT-RNAi, BDSC #40845)^36,77,144,145^. Controls included the wild-type w^1118^ strain (BDSC #5905), TH>GAL4/w^1118^ (TH driver-only), UAS>dVGLUT-RNAi/w^1118^ (undriven dVGLUT RNAi) to control for genetic background, vector, and insertion site. We also employed TH-driven expression of UAS>Luciferase (BDSC #35788) and UAS>LexA-RNAi (BDSC #67946)^77^ as controls in RNAi knockdown studies. For imaging experiments, DA neurons were labeled with green fluorescent protein (GFP) by recombining TH>GAL4 with 20xUAS>6xGFP (abbreviated as UAS>GFP; BDSC #52262)^146^. Reporter lines included UAS>MitoTimer (BDSC #57323)^54^ to measure mitochondrial oxidative stress and UAS>iATPSnFr (gift of Drs. Kevin Mann and Thomas Clandinin, Stanford University)^68^, a sensor of cytosolic ATP. We used UAS>αSyn^A53T^ (BDSC #8148) for GAL4-driven expression of the A53T mutant form of human α-synuclein (αSyn^A53T^). For intersectional luciferase reporters of DA neuron *dVGLUT* or *TH* expression, we employed LexAop>B3 recombinase (LexAop>B3R) and UAS>B3RT-STOP-B3RT-Luciferase strains as described previously^36^. A complete list of the RNAi lines used for forward genetic screening is provided in Table S4.

### Multiplex fluorescent *in situ* hybridization

Multiplex RNAscope-based fluorescent *in situ* hybridization in *Drosophila* brain was performed as previously described^147^. Briefly, adult central brains were rapidly dissected from male and female flies at 14 days (d) post-eclosion in ice-cold Schneider’s insect medium and fixed in 2% paraformaldehyde (in PBS, 25°C). Following washes in PBS with 0.5% Triton X-100 (PBST), brains were dehydrated in a graded series of ethanol dilutions and shaken in 100% ethanol (4°C, overnight). Brains were subsequently rehydrated with a series of ethanol solutions and incubated with 5% acetic acid (4°C) to enhance probe penetration. Following PBST washes, brains were fixed again in 2% paraformaldehyde (in PBS, 25°C) and placed into 1% NaBH_4_ (4°C). After a final PBST wash, brains were transferred onto Superfrost Plus slides (Thermo Fisher Scientific, Waltham, MA) and air-dried. Multiplex fluorescent *in situ* hybridization via RNAscope was subsequently performed according to manufacturer’s instructions (Advanced Cell Diagnostics, Hayward, CA) to detect mRNA expression of *TH* (*ple* gene, Cat. No. 536401-C2) and *dVGLUT* (*vglut* gene, Cat. No. 424011). Briefly, samples underwent: 1) protease treatment, 2) probe hybridization (2h, 40°C), and 3) probe amplification (40°C). Samples were then counterstained with DAPI and coverslipped; DAPI counterstain was excluded for GFP-labeled brains. *dVGLUT* and *TH* mRNAs were detected with fluorescent Alexa 488 and Atto 550 dyes, respectively; alternatively, Atto 550 labeled *dVGLUT* mRNA in GFP-labeled brains.

### Confocal microscopy of *Drosophila* brain fluorescent *in situ* hybridization

Images were acquired with an Olympus IX81 inverted microscope equipped with an Olympus spinning disk confocal unit (Olympus, Center Valley, PA), a Hamamatsu EM-CCD digital camera (Hamamatsu, Bridgewater, NJ) and a high-precision BioPrecision2 XYZ motorized stage with linear XYZ encoders (Ludl Electronic Products Ltd, Hawthorne, NJ). A 20x (0.8 NA) air objective (Olympus) was employed for manual quantification of GFP-labeled DA neuron numbers and *dVGLUT* mRNA expression in candidate RNAi lines. A 60x (1.4 NA) SC oil immersion objective (Olympus) was used for quantification of RNAi-mediated *dVGLUT* knockdown at randomly selected image sites across the fly brain. For these 60x images, image sites were systematically and randomly selected across the extent of each *Drosophila* brain using a grid of 100μm^2^ frames spaced by 200μm. 3D z-stacks were acquired in all imaging studies: 2048×2048 pixels for 20x images, and 1024×1024 pixels for 60x images; 0.2μm z-steps). Z-stacks were collected using optimal exposure settings (*i.e.*, those that yielded the greatest dynamic range with no saturated pixels), with differences in exposures normalized during image processing. Image collection was controlled by Slidebook 6.0 (Intelligent Imaging Innovations, Inc., Denver, CO). Analyses were conducted while blinded to genotype, sex, and treatment conditions.

### Confocal image analysis

Imaging data was initially analyzed via Slidebook and Matlab (MathWorks, Natick, MA) software. First, a Gaussian channel was made for each channel to quantify mRNA expression by calculating a difference of Gaussians using sigma values of 0.7 and 2. Then, 3D image stacks were separated every 20 images (summed to 4μm in the z-axis). Intensity values were averaged within each Gaussian channel in every image set to create 2D projection images of DAPI-stained nuclei (60x images) or GFP-labeled cells (20x images) alongside RNAscope-labeled TH and dVGLUT mRNA grains. To quantify mRNA expression within the respective TH and dVGLUT channels, 2D projection images were separated into quantitative TIFF files of each Gaussian channel and imported into the HALO image analysis platform (version 3.5, Indica Labs, Albuquerque, NM) equipped with a fluorescent *in situ* hybridization module. DAPI- or GFP-labeled cells and fluorescently-labeled mRNA grains from the dVGLUT and TH channels were quantified via HALO software based on the following thresholding criteria: any object 5-50μm^2^ for DAPI, 50-300μm^2^ for GFP, and 0.03-0.15μm^2^ for *TH* and *dVGLUT* mRNA grains. To determine the minimum number of mRNA grains associated with a DAPI-stained nucleus or GFP-labeled cell to be considered positive, we tested different thresholds of mRNA grain numbers above background levels (*i.e.*, the number of mRNA grains expressed in a typical cell volume). For 60x DAPI-stained images, thresholds of 3 *TH* mRNA grains and 8 *dVGLUT* mRNA grains were selected for quantifying positive cells; for 20x GFP-labeled images, 4 *dVGLUT* mRNA grains were selected as the threshold. All mRNA grains within 1μm of DAPI-stained nuclei were assigned to the respective cell; the 1μm border was reduced whenever necessary to prevent overlap between neighboring cells. *TH* and *dVGLUT* expression was quantified either as percent of all TH^+^ cells that were dVGLUT^+^, or as the average number of *dVGLUT* grains/TH^+^ cell. Experimenters were blinded to sex, genotype, and treatment until analyses were completed.

### Intersectional luciferase reporter assays

#### Construction of TH^+^/dVGLUT^+^ intersectional luciferase reporters

To measure changes in DA neuron dVGLUT expression, we assembled an intersectional genetic luciferase reporter: dVGLUT>GAL4/LexOP>B3R;TH>lexAop/UAS>B3RT-STOP-B3RT-Luciferase. Changes in TH expression specific to dVGLUT-expressing DA neurons were measured using a similarly constructed luciferase reporter: dVGLUT>LexA/lexAOP>B3R;TH>GAL4/UAS>B3RT-STOP-B3RT-Luciferase. For both intersectional reporter lines, B3R-mediated excision results in luciferase expression under the control of either dVGLUT or TH promoters specifically in TH^+^/dVGLUT^+^ neurons. We additionally generated TH>GAL4/UAS>Luciferase flies to measure luciferase expression more broadly in all TH-expressing DA neurons.

#### Luciferase assay

Assays were conducted as previously described^36^. Briefly, adult intersectional DA neuron dVGLUT and TH luciferase reporter flies were fed either drug- or vehicle-treated food with brains extracted at 14d or 60d post-eclosion to analyze drug-related changes in luciferase reporter activity. In parallel, a separate cohort was aged to 2d or 60d post-eclosion to analyze age-related changes in the activity of the respective luciferase reporters. Brains were rapidly dissected from decapitated flies in PBS with additional removal of associated cuticle and connective tissues; each sample comprised 5 male or 5 female fly brains. Luciferase assays were performed using the Dual-Luciferase Reporter Assay System (Promega, Madison, WI) per manufacturer instructions. Brains were placed in passive lysis buffer and homogenized with a pestle. Samples were then shaken at room temperature (45min, 1200rpm) and stored at -80°C until the assay was performed (within one month of brain collection). Once thawed, sample lysates were vortexed and diluted to ensure that luminescent signal was not oversaturated. Sample lysates were then added into a clear F-bottom 96-well plate in triplicate followed by addition of Luciferase Assay Reagent II. The resulting firefly luciferase-mediated luminescence was immediately read in a Pherastar FSX plate reader (BMG Labtech, Ortenberg, Germany). Firefly luciferase reactions were terminated with addition of Stop & Glo reagent; the presence of a fixed amount of *Renilla* luciferase in the buffer provided an internal control^148^. Triplicates for each sample were averaged, and results were reported as a luminescence ratio of firefly luciferase luminescence divided by *Renilla* luciferase luminescence (LUM ratio).

### Drosophila behavior

#### Paraquat behavior

To determine the impacts of 10mM paraquat on locomotor behavior, we used the TriKinetics *Drosophila* Activity Monitoring (DAM) system (TriKinetics, Waltham, MA) as described earlier^145^. Newly eclosed male and female flies were collected daily at fixed, regular times to ensure precise age determination. Flies (2d post-eclosion) were transferred individually into activity tubes containing standard cornmeal-molasses media in the presence or absence of paraquat. Tubes were then placed into DAM2 activity monitors under a 12h light:12h dark cycle. Flies were allowed to acclimate for 24h followed by continuous 24h activity monitoring for movement. Locomotor activity was measured as the number of times (counts) a fly crossed the infrared beam running through the middle of each activity tube per 24h period. Activity data from the flies were collected for either an individual day (day 2) or across several days (days 2-4) in the monitors and averaged over the three days to provide a mean number of counts/24h.

#### αSyn^A53T^ behavior

To identify the effect of DA neuron αSyn^A53T^ expression on locomotion, TH>GAL4 was recombined with UAS>αSyn^A53T^. Recombined TH>GAL4,αSyn^A53T^ flies were then crossed with candidate RNAi lines. The resulting adult offspring co-expressing the respective RNAi alongside αSyn^A53T^ (aged either 2d or 16d post-eclosion) were loaded into DAM2 activity monitors containing vehicle food. Locomotion was then recorded during days 2-4 in the activity monitors (days 4-6 and days 18-20 post-eclosion).

### *Ex vivo* multiphoton *Drosophila* brain imaging

#### Dissection and microscopy

Isolated *ex vivo* whole adult *Drosophila* brain preparations were obtained by rapid brain removal and microdissection in adult hemolymph-like saline (AHL, in mM: 108 NaCl, 5 KCl, 2 CaCl_2_, 8.2 MgCl_2_, 1 NaH_2_PO_4_, 10 sucrose, 5 trehalose, 5 HEPES, 4 NaHCO_3_; pH 7.4, 265 mOsm). Brains were immobilized via pinning onto slabs of thin Sylgard with fine tungsten wire as described previously^145,149^. Brain preparations were imaged while bathed in AHL on a Bergamo II resonant-scanning two-photon microscope (Thorlabs, Newton, NJ) using a 20x (1.0 NA) water immersion objective lens (Olympus) and an Insight X3 laser (Spectra-Physics Inc., Milpitas, CA). Less than 50mW mean power was delivered through the objective with both gain and pockels cell settings for power modulation fixed for all imaging setups unless otherwise stated. The tunable output of the laser was mode-locked at 920nm for imaging of GFP, iATPSnFR, or unoxidized MitoTimer, while the fixed 1045nm output imaged oxidized MitoTimer. Fluorescent emissions were collected using a 525/50nm full-width half-maximum (FWHM) bandpass emission filter for GFP, unoxidized MitoTimer, and iATPSnFr (green channel) and a 607/70nm FWHM bandpass filter for oxidized MitoTimer (red channel). Imaging parameters were optimized to avoid pixel saturation. Data acquisition was performed with ThorImage software (versions 4.0/4.1; Thorlabs). Z-stacks of optical sections were acquired through the depth of the fly brain in 2μm z-steps with 8x frame averaging, except for iATPSnFr studies, which were collected in 3μm z-steps with no averaging. Images were 16-bit, 1024×1024 pixels (pixel size 0.64μm), except for KCl studies which were 512×512 pixels (pixel size 1.28μm, 15.1 frames/s). Average z-projections of the image stacks were generated using the Fiji/ImageJ image processing software package (National Institutes of Health, Bethesda, MD)^150^.

#### Multiphoton image analyses

**GFP-labeled DA neuron quantitation:** GFP-labeled DAergic cell bodies were identified throughout z-stacks from brains of adult male and female flies with the following genotypes: TH>GAL4,UAS>GFP/UAS>LexA-RNAi and TH>GAL4,UAS>GFP/UAS>dVGLUT-RNAi. Analyses focused on DA neuron clusters across several brain regions including the PAL, PAM, and SOG regions, as previously described^36^. For DAergic cell body counts, comparable results were independently obtained from n≥3 blinded experimenters; the median of the results across reviewers was used for analysis. **MitoTimer analyses:** For analysis of the MitoTimer biosensor, mean pixel fluorescence intensity from green and red emission channels was collected in the SOG, EB, and FSB brain regions. Levels of MitoTimer oxidation were expressed as the ratio of MitoTimer red to green fluorescence (red:green ratio). MitoTimer-labeled DAergic projection areas were analyzed throughout z-stacks in adult male and female flies. Genotypes tested included: TH>GAL4,UAS>MitoTimer/UAS>dVGLUT-RNAi, TH>GAL4,UAS>MitoTimer/UAS>Luciferase, and TH>GAL4,UAS>MitoTimer/UAS>LexA-RNAi. Effects of DA neuron depolarization on mitochondrial ROS production were assessed by continuously perfusing brains expressing TH-driven MitoTimer with 40mM KCl (in AHL, adjusted for ionic osmolarity). Z-stacks of red and green channels were acquired before and during constant perfusion with AHL containing 40mM KCl.

Mean pixel fluorescence intensity of red and green channels was quantified across timepoints. **iATPSnFR studies:** iATPSnFr-labeled DAergic projection areas (SOG, EB and FSB) were analyzed throughout brains of adult male and female flies (genotypes tested: UAS>iATPSnFr/w^1118^;TH>GAL4/UAS>dVGLUT-RNAi and UAS>iATPSnFr/w^1118^;TH>GAL4/UAS>LexA-RNAi). For studies of KCl-induced DA neuron depolarization, z-stacks were collected before and during perfusion with AHL containing 40mM KCl. The mean pixel fluorescent intensity value pre-KCl (F*_initial_* or F*_i_*) was subtracted from each post-KCl timepoint (F) and then divided by F*_i_* to provide the change in iATPSnFR fluorescence relative to the baseline measurement: (F - F*_i_*)/F*_i_*, abbreviated as Λ1F/F*_i_*. All multiphoton image analyses were conducted using Fiji/ImageJ software.

### *Drosophila* single-cell RNA sequencing

Central brains from adult male and female *Drosophila* were dissected as previously described^151^. Single cells were encapsulated using the Chromium Single Cell 3′ Reagent Kit v3 (10x Genomics, Pleasanton, CA) according to manufacturer instructions. Barcoded mRNAs were extracted, followed by amplification and sequencing of the cDNA libraries on an Illumina HiSeq4000 platform (Illumina, San Diego, CA).

Experimental data from two published^151,152^ and two unpublished experiments were combined and aligned to the *Drosophila melanogaster* reference genome (DMel6.25)^153^. Two-dimensional reduction and clustering were performed using the Seurat v3 R package^154^. Monoaminergic cells were extracted based on the expression levels of the *Drosophila* vesicular monoamine transporter (*Vmat*). Extracted cells were subsequently re-clustered using parameters that were optimized for the reduced transcriptional heterogeneity of the selected cells. Experimental- and replicate-based variation was regressed out, resulting in equal representation of all replicates in each cell cluster. Candidate marker genes were plotted to identify cell clusters of interest. A subpopulation of DA/glutamate neurons was identified based on co-expression of dVGLUT alongside DA neuron markers (dVMAT^+^/TH^+^/DAT^+^/dVGLUT^+^). Thresholds for each gene were set manually.

## Mouse studies

### Animal husbandry

Animals were housed and handled in accordance with all appropriate NIH guidelines through the University of Pittsburgh Institutional Animal Care and Use Committee. We abided by all appropriate animal care guidelines including ARRIVE (Animal Research: Reporting of *In Vivo* Experiments) guidelines for reporting animal research^155^. Mice were housed in cages with a 12h light:12h dark cycle and had access to food and water *ad libitum*. All efforts were made to ameliorate animal suffering.

### Mice

7-11-wk-old male and female mice were acquired in part from The Jackson Laboratory (Bar Harbor, ME): B6.SJL-Slc6a3^*tm1.1(cre)Bkmn*^/J (DAT^IRES*cre*^; stock #006660), *Slc17a6^tm^*^2^(cre)*^Lowl^*/J (VGLUT2^IRES*cre*^; stock #016963), and B6.Cg-*Gt(ROSA)26Sor^tm^*^9^(CAG–tdTomato)*^Hze^*/J (Ai9 tdTomato; stock #007909). All mice were congenic with the C57BL/6J background. VGLUT2-expressing DA neurons were labeled in double heterozygous VGLUT2^IRES*cre*^;TH^2A-flpo^ mice; construction and validation of these mice was described previously^23,28^. The overall DA neuron population was labeled in double heterozygous DAT^IRES*cre*^;Ai9 tdTomato mice.

### Viral vectors and stereotaxic surgeries

VGLUT2^IRES*cre*^;TH^2A-flpo^ mice (14 males, 12 females) were anesthetized with isoflurane and placed onto a stereotaxic apparatus (Kopf Instruments, Tujunga, CA). Following disinfection with betadine and 70% isopropyl alcohol, an incision exposed the skull for viral injections into the VTA using the following stereotaxic coordinates (from bregma) that varied according to mouse weight (g): 12-16g, -3.0mm anterior/posterior (AP), -4.1mm dorsal/ventral (DV), 0.6mm lateral (L); 17-24g, -3.3mm AP, -4.3mm DV, 0.6mm L; >25g, -3.44mm AP, -4.5mm DV, 0.6mm L. Injections of INTRSECT2.0 viruses were made bilaterally and included AAV8-EF1a-Cre^ON^,Flp^ON^-EYFP (1μL, titer 5.01×10^12^ vg/mL), in combination with either AAV8-EF1a-Cre^OFF^Flp^ON^-mCherry (1μL, titer 1.30×10^13^ vg/mL) or AAV8-EF1a-Cre^OFF^Flp^ON^-oScarlet (1μL, titer 1.15×10^12^ vg/mL). The injection rate was 0.1μL/min, and the needle was left in place for 5 min to minimize backflow along the injection tract, then withdrawn. Injected mice were allowed to recover for ∼5 wks post-injection to ensure adequate AAV-mediated fluorescent label expression.

### Mouse brain clearing

Brains were rapidly extracted from INTRSECT2.0 AAV-injected mice ∼5 wks post-injection and incubated overnight in 4% paraformaldehyde in PBS at 4°C. Brains were subsequently cleared using the CUBIC approach^156,157^ by immersion in 50% CUBIC R1 for 1d, followed by solution changes to 100% CUBIC R1 on days 2, 3, 7 and 10. Brains were then washed with PBS before immersion in CUBIC R2 (No TEA) for 2d until optically clear.

### Ribbon scanning confocal microscopy of mouse brain

Whole cleared INTRSECT2.0-labeled mouse brains were mounted in CUBIC R2 (No TEA) and imaged via ribbon scanning confocal microscopy (RSCM) on a Caliber RS-G4 microscope (Caliber I.D., Andover, MA) as described previously^158^. The microscope was equipped with a Nikon CFI90 20XC glycerol objective (1.0 NA), and scans were completed with a lateral pixel resolution of 0.274μm. Laser power was adjusted linearly between 15-30% for brain imaging with volumes acquired using 10.67μm z-steps.

### Whole mouse brain imaging analyses

3D RSCM volumetric data was assembled by first stitching composites using a dedicated computational cluster. Composites were then assembled into the imaris (.ims) file format for volumetric rendering via Imaris software (version 9.0; Bitplane AG, Zurich, Switzerland). Imaris files were the primary source used for all downstream processing pipelines. For brain volume registration, images first underwent contrast stretching and background removal using the Accelerated Pixel and Object Classification plugin for napari^159^. This was followed by image registration to the Allen Mouse Brain Common Coordinate Framework (CCFv3)^73^ using the brainreg pipeline^160^. Cell body detection within the 3D volumes of INTRSECT2.0-labeled brains was conducted using cellfinder software, a constituent of the BrainGlobe package of computational neuroanatomy tools^161^. This was followed by classification of cell candidates using ResNet-50 neural network architecture^74^ implemented in the fastAI deep learning library^162^. Binarization was completed using the ilastik software toolkit^75^ for annotation and training. The inference was done on a dedicated dask cluster^163^. Soma density and percentage of neuron projection-positive pixels were calculated based on CCFv3 structure volumes.

### Data access

Imaris files of brain imaging data will be deposited in the Brain Image Library (BIL) optical microscopy data archive and made available for public access. Links to complete volumetric datasets used for this study will be included here once available.

### Mouse brain fluorescent-activated cell sorting (FACS) and bulk RNA sequencing

#### Mice

Heterozygous male and female INTRSECT2.0 VGLUT2^IRES*cre*^;TH^2A-flpo^ mice (∼5 wks post-viral injection) and DAT^IRES*cre*^;Ai9 tdTomato were used.

#### Tissue extraction and neuron dissociation

Brains were rapidly extracted and rinsed with chilled artificial cerebrospinal fluid (ACSF). Midbrain regions (Bregma -2.88 to -3.88mm) were separated from each brain using a stainless-steel mouse brain matrix (1mm) and single edge blades (on ice), followed by transfer into conical tubes containing chilled ACSF for dissociation. To ensure sufficient cDNA yield, midbrain sections were pooled, such that each sample contained 2 midbrains of the same sex (final n=7 male, 6 female INTRSECT2.0 VGLUT2^IRES*cre*^;TH^2A-flpo^ samples; n=2 male, 3 female DAT^IRES*cre*^;Ai9 tdTomato samples). Midbrain samples were dissociated using the Miltenyi Neural Dissociation kit (Miltenyi Biotec, Bergisch Gladbach, Germany) according to manufacturer’s instructions, in combination with a Miltenyi gentleMACS Octo Dissociator with Heaters, Miltenyi gentleMACS C Tubes, and Miltenyi MACS SmartStrainer (70μm). After straining, cells were resuspended in DPBS supplemented with BSA for fluorescent-activated cell sorting (FACS).

#### FACS cell sorting

Individual cell suspensions were sorted by the Unified Flow Core FACS facility at the University of Pittsburgh using a BD FACSAria II (BD Biosciences, Franklin Lakes, NJ). Cells were sorted for viability using Ghost Dye Violet 450 (Tonbo Biosciences, San Diego, CA), by cell size, by fluorescence: EYFP and mCherry/oScarlet (for cells from INTRSECT2.0 mice) or tdTomato (for cells from DAT^IRES*cre*^;Ai9 tdTomato mice). We ensured that we did not collect TH^+^ monocytes^164^ by specifically collecting either EYFP^+^/CD45^−^/CD11b^−^, mCherry^+^ or oScarlet^+^/CD45^−^/CD11b^−^, or tdTomato^+^/CD45^−^/CD11b^−^ cells. Antibodies for the sorting included an anti-CD45 antibody (RA3-6B2) conjugated to Alexa Fluor 647 (Thermo Fisher Scientific) as described previously^165^ as well as an anti-CD11b antibody which was grown, purified, and conjugated to Pacific Blue dye in the laboratory of Dr. Mark Shlomchik (University of Pittsburgh). Appropriately labeled neurons were sorted directly into prechilled 96-well PCR plates; 2000 cells/well were sorted in triplicate for each sample.

#### Bulk RNA sequencing

RNA extractions, cDNA generation, and library preparation were performed by the University of Pittsburgh Health Science Sequencing Core at the UPMC Children’s Hospital of Pittsburgh. RNA was extracted using the Qiagen RNeasy Plus Micro extraction kit (Qiagen, Redwood City, CA) following manufacturer’s instructions. RNA was assessed for quality on an Agilent Fragment Analyzer 5300 using the High Sensitivity RNA kit (Agilent, Santa Clara, CA). The mean RNA integrity number (RIN) for samples was 8, indicating excellent quality for RNA sequencing. 4.5μL RNA was used from each sample for cDNA generation using the Takara Smart-Seq v4 Ultra Low Input RNA kit (Takara Bio USA, Ann Arbor, MI) following manufacturer’s instructions, with 15 cycles of cDNA amplification. Smart-Seq cDNA was assessed for quality on an Agilent Fragment Analyzer 5300 using the high sensitivity NGS kit (Agilent). All samples passed QC with full length cDNA (mean concentration of 13ng/μL; primary peaks ∼2000 bp; absence of short length cDNA with bimodal peaks including second peak at ∼300 bp). Library preparation was performed using 1ng of cDNA input with the Illumina Nextera XT kit (Illumina) and UDI indexes (Illumina) added using 12 PCR cycles. Libraries were assessed using an Agilent High Sensitivity NGS kit, and then normalized and pooled by calculating the nM concentration based on the fragment size and the concentration. Prior to sequencing, library pools were quantified by quantitative polymerase chain reaction (qPCR) on the LightCycler 480 using the KAPA qPCR quantification kit (KAPA Biosystems, Boston, MA). Libraries were sequenced on a NovaSeq 6000 system (Illumina) at UPMC Genome Center on an S2 100 cycle flow cell, 2 x 50 bp, for an average of ∼40 million reads/sample.

#### RNA sequencing analyses

FastQC (version 0.11.7) determined the per base sequence quality, with a mean score of 36 across samples. Paired-end reads were preprocessed, and adapters and low-quality bases were removed using trimmomatic (version 0.38). Trimmed reads were mapped to *Mus musculus* Ensembl GRCm38 using HISAT2 (version 2.1.0), with a mean overall alignment rate of 92.32% across samples. After mapping, a total of 46,080 Ensembl transcripts (including the EYFP and tdTomato transcripts used to label the desired respective cell populations) were filtered to remove low expression transcripts, retaining transcripts with at least one count per million (CPM) in half of the samples. After the filtering, 11,133 remained for DE analysis. RNAseq data were analyzed via DESeq2^166^ using neuron type and sex as the main effects. Principal component analysis was performed to assess sample heterogeneity and remove outliers using R package ggplot and the R function prcomp. Two samples had a first or second principal component outside three standard deviations of the mean and were removed from analyses.

### RNA sequencing analysis across *Drosophila* and mice

Differential expression (DE) was assessed in *Drosophila* using Seurat^167,168^ and in mice using limma^169^. Transcripts with p<0.01 (-log_10_(P)>2) and log_2_ fold change (FC) >0.25 or <-0.25 were considered DE in dVGLUT^+^ DA neurons compared to dVGLUT^−^ DA neurons for *Drosophila* and in INTRSECT2.0 EYFP^+^/VGLUT2^+^ DA neurons compared to DAT;CreTd-Tomato^+^/VGLUT2^−^ DA neurons for mice. DE gene lists for *Drosophila* and mice were entered into Metascape^170^ and assessed for pathway overrepresentation (Gene Ontology, Kyoto Encyclopedia of Genes and Genomes, Reactome, WikiPathways, CORUM) with expressed transcripts as background. DE gene lists were also compared via MitoCarta version 3.0^71^ to assess mitochondria-specific pathway overrepresentation via threshold-free gene set enrichment analysis (GSEA) using R package fgsea^171^, which captures cumulative evidence of pathway enrichment across the entire gene list. Mouse DE gene sets (male and female combined analysis DE genes, DE genes from male-only analysis, and DE genes from female-only analysis) were analyzed using Hypergeometric Optimization of Motif Enrichment (HOMER) to predict transcription factors that may be upstream regulators of DE transcripts; genes were analyzed for both known and *de novo* motifs^172^.

### Differentially expressed gene candidate selection for screening

Of the 423 DE genes in the *Drosophila* RNAseq dataset, we narrowed down the list of candidates first to 224 genes by selecting the following discrete groups: 1) the top 50 upregulated genes (top 25 by magnitude expression change and top 25 by fold change), 2) the top 50 downregulated genes, 3) the top 50 genes displaying sex differences in dVGLUT^+^ DA neurons (top 25 genes upregulated in males compared to females and top 25 upregulated in females) and 4) 74 genes of interest to DA neuron function and PD. Next, these 224 genes were narrowed down to 159 after removing 36 duplicates and 22 genes without mammalian orthologs. All candidate gene and control UAS>RNAi lines were obtained from either BDSC, Vienna Drosophila Resource Center (VDRC), or the Shigen National Institute of Genetics (NIG-FLY) (all lines listed in Table S4). We used only VALIUM20 vector-derived fly stocks from the Harvard Transgenic RNAi Project (TRiP)^77^ or flies from the VDRC KK collection^173^. 7 gene candidates did not have available RNAi lines from the VALIUM20 or KK collections and were therefore removed. Comparisons with mouse RNAseq data revealed 21 additional genes orthologous to *Drosophila* DE genes that were also upregulated or downregulated in the mouse RNAseq dataset (Table S3), yielding a list of 180 candidate genes. Finally, we performed Hypergeometric Optimization of Motif Enrichment (HOMER) analysis for known and *de novo* motifs in the mouse RNAseq dataset and identified transcription factors that may be upstream regulators of DE transcripts. We selected 7 transcription factors of interest, including 2 factors whose orthologs were already found to be DE genes in the *Drosophila* dataset, and added them to the list of candidates to yield a final total of 187 candidates to be screened for vulnerability to PD models.

### RNAi screens

For RNAi screen experiments, candidate RNAi lines were crossed with TH>GAL4 to drive candidate gene knockdown in DA neurons. In secondary and tertiary screens, RNAi lines were crossed with recombined TH>GAL4,UAS>αSyn^A53T^ or TH>GAL4,UAS>GFP flies to co-express the αSyn^A53T^ mutant or GFP, respectively, alongside the respective candidate gene RNAi in DA neurons. For all experiments utilizing drug treatments, flies were randomly assigned to vehicle or drug treatment groups.

#### Primary paraquat screen

TH>GAL4-driven candidate UAS>RNAi flies (2d post-eclosion) were placed into tubes containing either vehicle or 10mM paraquat-supplemented food. After 24h acclimation, locomotion was monitored throughout days 2-4 of drug exposure. Locomotion of each paraquat-exposed control and candidate RNAi line was displayed as percent of the locomotion of each line’s vehicle-treated animals. To adjust for significant sex differences in locomotion and vulnerability in female versus male flies attributable to the RNAi repository source, female VDRC RNAi fly locomotion counts were multiplied by 1.093 before analysis.

#### Secondary αSyn^A53T^ screen

TH>GAL4,UAS>αSyn^A53T^/UAS>RNAi candidate flies (2d post-eclosion) were placed into tubes containing vehicle food, and locomotion of each candidate was monitored during days 2-4 in DAM2 *Drosophila* activity monitors (TriKinetics) (days 4-6 post-eclosion). Locomotion of each control and candidate RNAi line was displayed as percent of the locomotion of each line without αSyn^A53T^ expression. To adjust for significant sex differences in locomotion and vulnerability in female flies compared to males attributable to the RNAi repository source, female VDRC RNAi fly locomotion counts were multiplied by 1.072 before analysis.

#### Tertiary DA neuron dVGLUT expression screen

DA neuron number and *dVGLUT* mRNA expression via RNAscope were analyzed within the brains of candidate gene RNAi lines. We analyzed GFP-labeled DA neurons in *LexA*, *mirr*, *caup*, *trh*, *oc*, *Dll* and *dj-1β* RNAi flies. To create these fly strains, the recombined TH>GAL4,UAS>GFP line was crossed with each UAS>RNAi line. *dVGLUT* mRNA expression in GFP-labeled DA neurons was quantified in *LexA* and *dj-1β*

RNAi lines. Brains were dissected at 14d post-eclosion, after 10d of exposure to either vehicle or 10mM paraquat food.

### Statistical analyses

Luciferase assays were analyzed via two-way analysis of variance (ANOVA) with age or treatment and sex or reporter line as between-subjects factors. In multiplex RNAscope studies, DA neuron dVGLUT expression differences were analyzed via one-way ANOVA. For RNAscope studies examining the impacts of genetic screen candidates on DA neuron dVGLUT expression, three-way ANOVA was employed with treatment, sex, and genotype as between-subjects factors. In behavioral studies, locomotion differences were analyzed using two-way ANOVA with treatment and genotype as between-subjects factors, or by three-way ANOVA with treatment, sex, and genotype as between-subjects factors. GFP-labeled DA neuron counts were analyzed using three-way ANOVA with treatment, sex, and genotype as between-subjects factors. MitoTimer red:green ratio results were analyzed using three-way ANOVA with treatment, age, genotype, sex and/or region as between-subjects factors. TEMPOL-treated dVGLUT-RNAi MitoTimer data in males were analyzed using an unpaired Student’s t-test, and KCl-treated MitoTimer results were analyzed using a mixed effects two-way ANOVA (to accommodate inclusion of animals with missing values) with timepoint as a within-subjects variable and genotype as a between-subjects variable. iATPSnFr responses were analyzed using two-way ANOVA with sex and genotype as between-subjects factors. A two-tailed Pearson correlation coefficient between MitoTimer red:green ratio and iATPSnFr peak fluorescence across DAergic regions was calculated. All significant main effects and interactions were followed up with Bonferroni post-hoc analyses, except for RNAi screen locomotion, which was followed up via Fisher’s LSD test with Benjamini, Krieger and Yekutieli two-stage linear step-up False Discovery Rate correction (significance defined as q<0.05). Statistical significance was defined as p<0.05, apart from RNA sequencing thresholding (p<0.01). Unless otherwise stated, statistical functions were completed using GraphPad Prism (version 9.3; GraphPad Software, San Diego, CA).

## Acknowledgments

We thank Drs. Edward Burton, Teresa Hastings, David Krantz, Mark Shlomchik, Kevin Mann, and Matthew Lycas for helpful advice and discussions. We also gratefully acknowledge assistance with movie assembly from Mary Brady. Additional schematic figures were created with BioRender.com. This work was supported by the National Institutes of Health F31NS118811 (S.A.B.), R01ES034037 (Z.F.), R21AG068607 (Z.F.), R21DA052419 (Z.F., R.W.L.), R21AA028800 (Z.F., R.W.L.), T32MH019986 (S.J.M.), R01MH117128 (S.R., S.M.), R01MH096985 (K.N.F.), R01HL150432 (R.W.L.), and R01HL150432-S1 (R.W.L.). C.D.T., Y.W., and S.W. were supported by funds from an ERC Advanced Grant (789274), a Wellcome Principal Research Fellowship in Basic Biomedical Sciences (200846), and a Wellcome Discovery Award (225192) to S.W. This research was also supported in part by the University of Pittsburgh Center for Research Computing, RRID:SCR_022735, through the resources provided. Specifically, this work used the HTC cluster, which is supported by NIH award number S10OD028483.

## Author contributions

S.A.B. and Z.F. conceived the idea and designed the study. S.A.B., S.A.R., T.K., C.D.T., L.E.F., S.J.M., V.R.S., Z.Y., D.S., K.D.K., D.D.B.K., I.V., M.C.S., F.J.W., M.Q.E.O., E.I.O., E.A., C.R., Y.S.K., Y.W., W.A.M., R.E., S.M., and C.E.J.C. performed the experiments. S.A.B., C.D.T., X.X., W.W., S.W., and R.W.L. performed the transcriptomic analyses. I.V., M.C.S., E.A., and A.M.W. performed the mouse brain imaging and associated analyses. S.A.B., S.A.R., T.K., C.D.T., X.X., S.J.M., I.V., W.W., W.A.M., R.E., A.M.W., R.W.L., and Z.F. analyzed the data. S.A.B. and Z.F. wrote the manuscript with contributions from the co-authors.

## Inclusion and diversity

We support inclusive, diverse, and equitable conduct of research.

---

**Figure S1.**
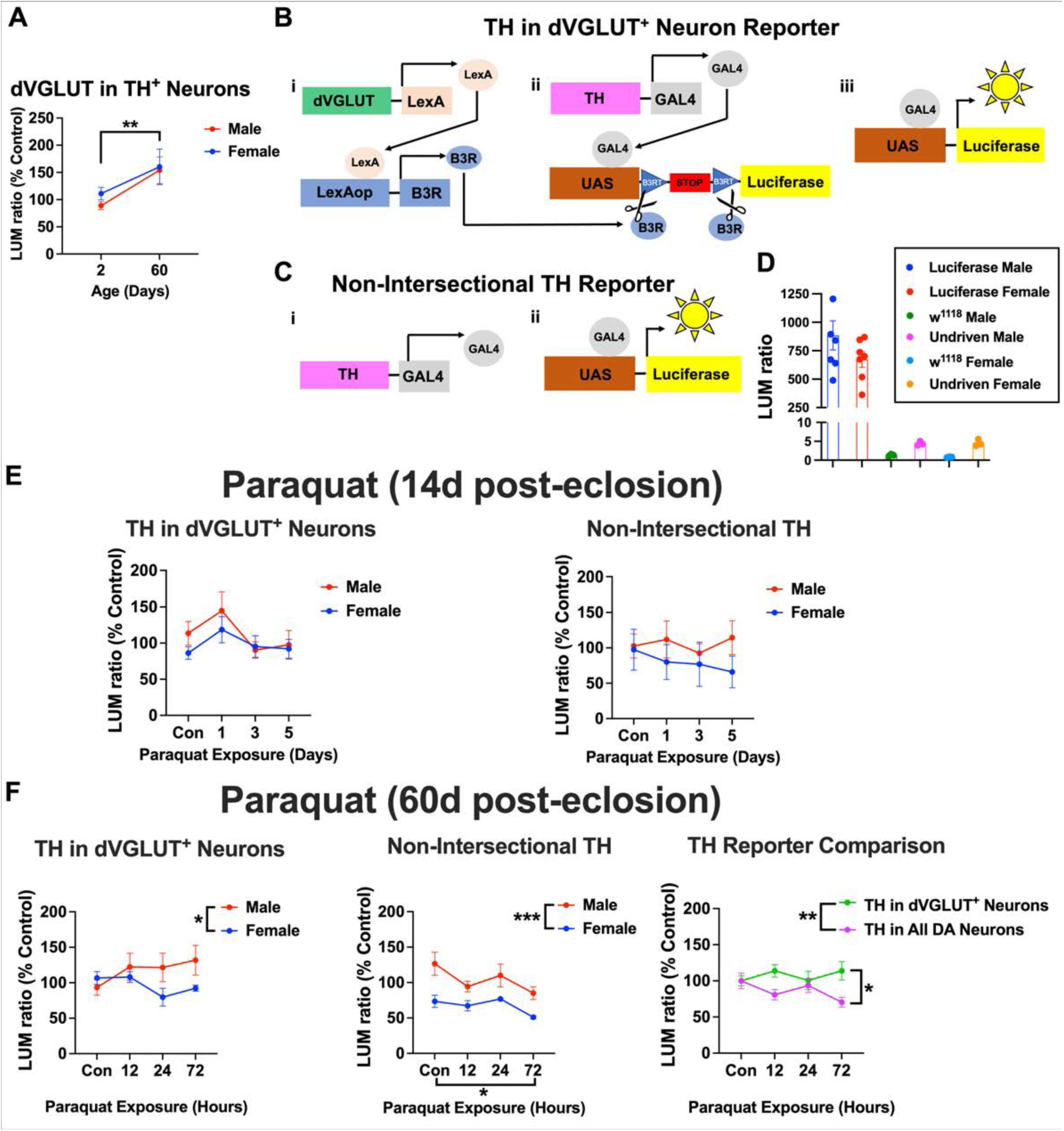
Comparison of intersectional and non-intersectional global reporters of TH expression, related to Figure 2. **(A)** There was an age-related increase in DA neuron dVGLUT expression in older male and female flies (60d post-eclosion) compared to young flies (2d post-eclosion) (Main effect of age: F_1,20_=9.27, p=0.0064, two-way ANOVA). Luminescent DA neuron dVGLUT reporter data were normalized to % young male and female controls. N=5-7 homogenates of 5 brains per group. **(B)** Schematic detailing an intersectional luciferase reporter of TH expression in TH^+^/dVGLUT^+^ DA neurons (dVGLUT>LexA/LexAop>B3R;TH>GAL4/UAS>B3RT-STOP-B3RT-Luciferase). Panel i: The dVGLUT promoter drives expression of the LexA transcriptional activator to selectively express B3 recombinase (B3R) in dVGLUT^+^ glutamatergic neurons. Panel ii: In parallel, the TH promoter drives expression of the GAL4 transcriptional activator in TH^+^ DA neurons. Panels iii: In the subpopulation of TH^+^/dVGLUT^+^ neurons where both B3R and GAL4 are co-expressed, B3R excises a transcriptional stop cassette within UAS>Luciferase. This enables GAL4 transactivation to express luciferase, generating measurable luminescence. **(C)** Schematic of the non-intersectional global luciferase reporter of TH expression across the overall population of DA neurons (TH>GAL4/UAS>Luciferase). **(D)** Quantification of luminescence produced by the non-intersectional global TH reporter showed robust luminescence when driven in TH^+^ DA neurons. There was negligible luciferase activity in wild-type w^1118^ and undriven (UAS>Luciferase only) male and female control flies. **(E)** In adult flies (14d post-eclosion), 10mM paraquat exposure did not significantly alter TH expression in male or female TH^+^/dVGLUT^+^ neurons using the intersectional TH reporter (p>0.05, two-way ANOVA; left panel) or across the overall DA neuron population via the non-intersectional global TH reporter (p>0.05, two-way ANOVA; right panel). N=6-9 homogenates per individual reporter; each homogenate was composed of 5 brains. **(F)** In aged flies (60d post-eclosion), 10mM paraquat-exposed males exhibited higher TH expression in TH^+^/dVGLUT^+^ neurons compared to females using the intersectional TH reporter (main effect of sex: F_1,47_=4.11, p=0.048, two-way ANOVA; left panel). In the broader DA neuron population, there was a similar sex difference between males and females throughout exposure groups (F_1,44_=16.44, p=0.0002, two-way ANOVA; middle panel) and paraquat exposure decreased TH expression (F_3,44_=3.19, p=0.033, two-way ANOVA; Bonferroni multiple comparisons test at 72h versus Con: p=0.015). Overall, DA neuron dVGLUT expression protected against paraquat-induced decreases in TH expression (main effect of reporter: F_1,99_=7.66, p=0.0067, two-way ANOVA; Bonferroni multiple comparisons test of intersectional TH reporter versus non-intersectional TH reporter at 72h: p=0.019; right panel). N=4-10 homogenates per group for individual reporters, N=8-20 homogenates for TH reporter comparison; each homogenate was composed of 5 brains. *p<0.05, **p<0.01, ***p<0.001; all data are plotted as Mean±SEM.

**Figure S2.**
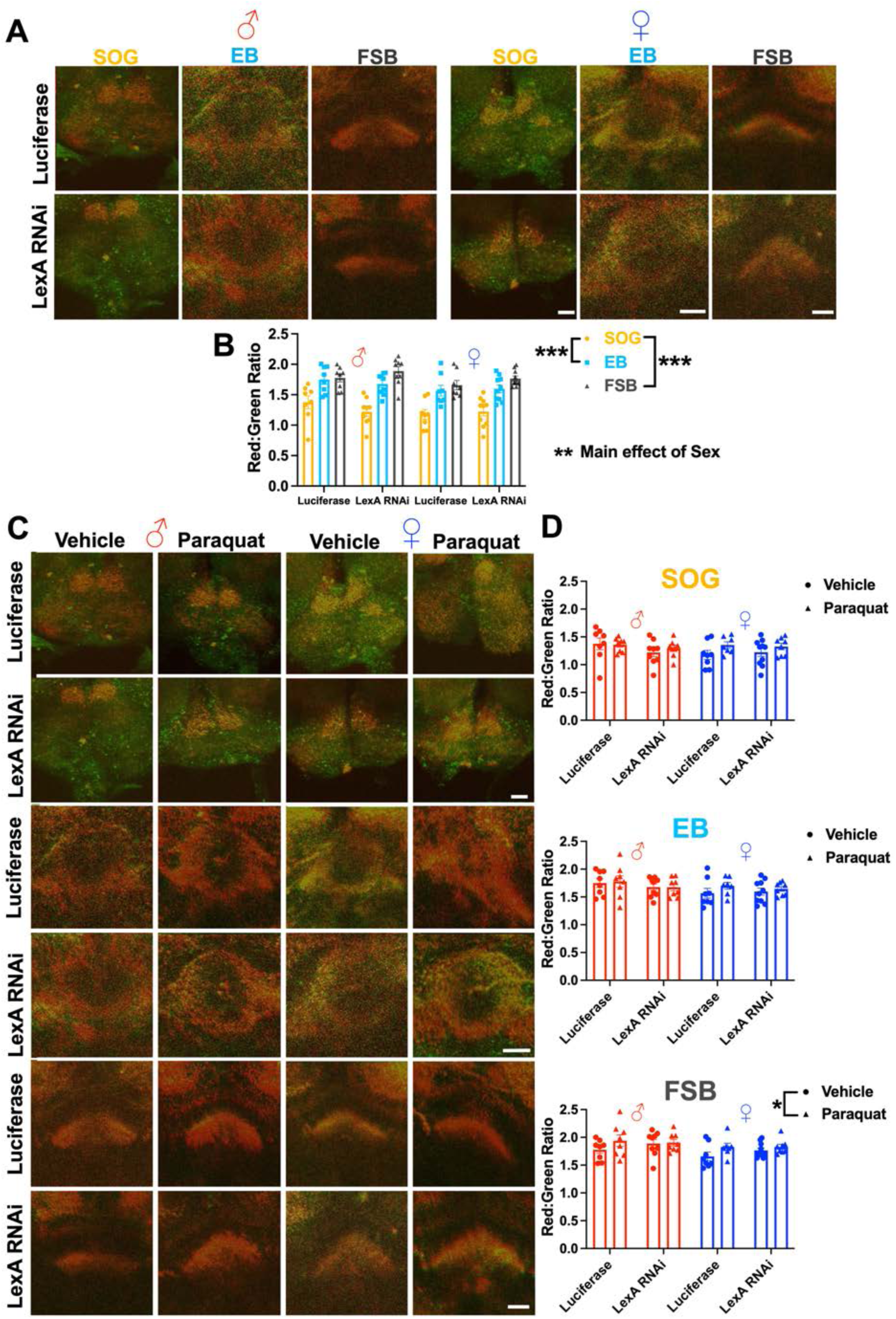
Control line does not impact sex-specific and region-specific differences in mitochondrial oxidative stress, related to Figure 3. **(A)** Representative projection images of MitoTimer-labeled DAergic projections to the SOG, EB and FSB regions in adult (14d post-eclosion) male and female brains (with green and red channels overlaid) from the following control strains: TH>GAL4,UAS>MitoTimer/UAS>Luciferase (Luciferase) and TH>GAL4,UAS>MitoTimer/UAS>LexA-RNAi (LexA RNAi) flies. Scale bars=25μm. **(B)** There were significant differences in baseline MitoTimer red:green ratios based on brain region (main effect of region: F_2,93_=55.61, p<0.001, three-way ANOVA) and sex (main effect of sex: F_1,93_=7.59, p=0.0071, three-way ANOVA) in the TH-driven Luciferase and *LexA* RNAi controls. However, there were no effects of the respective control lines on the red:green ratios (p>0.05, three-way ANOVA). N=8-10 per group. **(C)** Representative projection images of MitoTimer-labeled DAergic projections to the SOG, EB and FSB labeled by MitoTimer (with green and red channels overlaid) from adult male and female Luciferase and *LexA* RNAi control flies exposed to either paraquat (10mM, 5d) or vehicle (0mM, 5d). Scale bars=25μm. **(D)** Quantification of MitoTimer red:green ratios demonstrated a main effect of paraquat in DAergic projections to the FSB (F_1,58_=4.52, p=0.038, three-way ANOVA), but no significant effects of TH-driven Luciferase or *LexA* RNAi controls or interaction between controls and paraquat in any brain region (p>0.05, three-way ANOVA). N=7-10 per group. *p<0.05, **p<0.05, ***p<0.001; all data are plotted as Mean±SEM.

**Figure S3.**
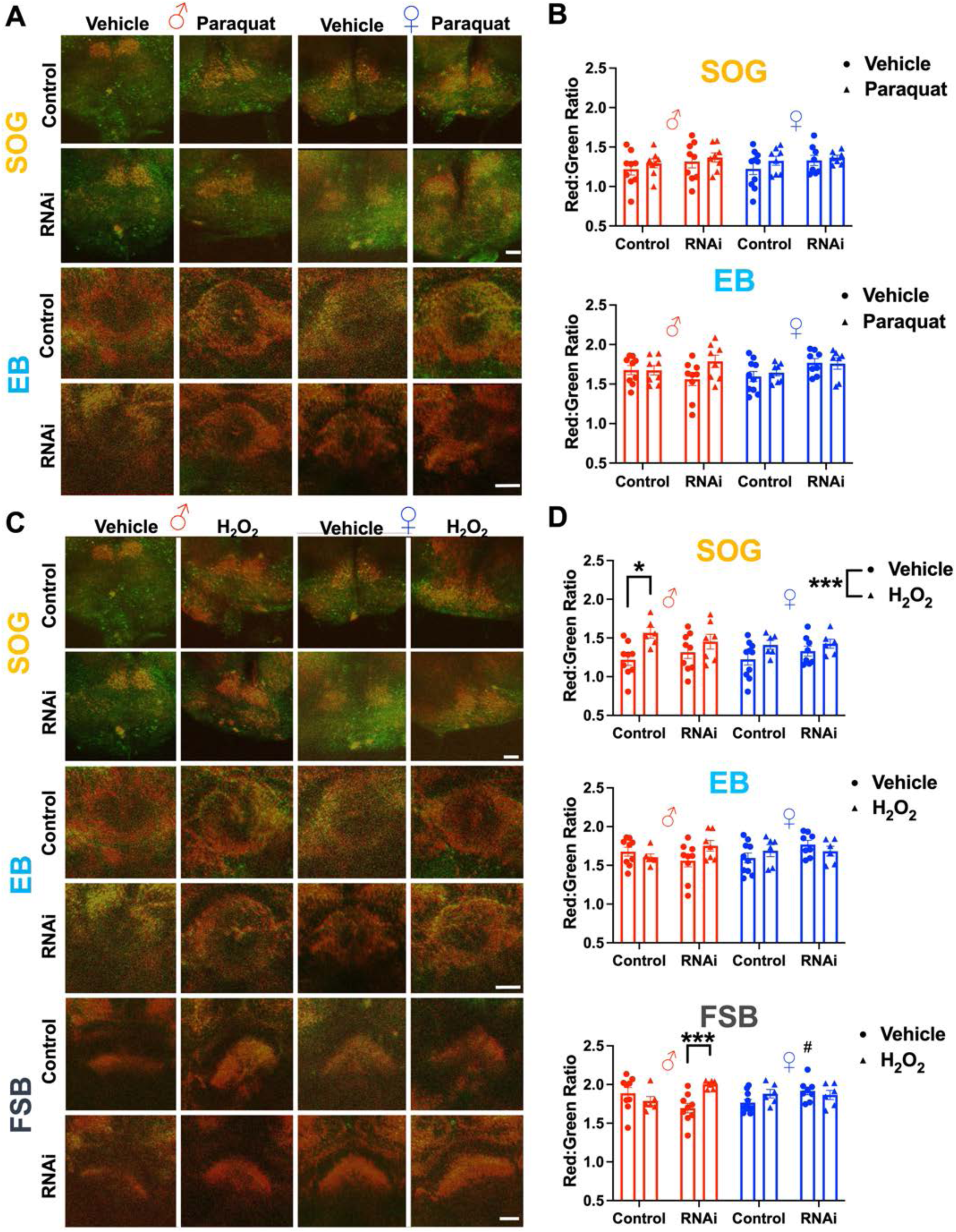
Region-specific impacts of paraquat and H_2_O_2_ on dVGLUT-mediated mitochondrial oxidative stress in DAergic projections, related to Figure 3. **(A)** Representative projection images of MitoTimer-labeled DAergic projections to the SOG and EB in adult (14d post-eclosion) dVGLUT RNAi (RNAi) versus *LexA* RNAi (Control) male and female flies; green and red fluorescent channels are overlaid. Flies were exposed to either paraquat (10mM, 5d) or vehicle control (0mM, 5d). Scale bars=25μm. **(B)** Quantification of MitoTimer red:green ratios in DAergic projections to the SOG or EB revealed no significant changes in response to paraquat (10mM) versus the vehicle control (0mM) in either males or females (p>0.05, three-way ANOVA). N=7-10 per group. **(C)** Representative projection images of MitoTimer-labeled DAergic projections to the SOG, EB and FSB from adult male and female RNAi or Control flies exposed to either H_2_O_2_ (3%, 2d) or vehicle (0%, 2d). Scale bars=25μm. **(D)** Quantification showed a main effect of H_2_O_2_ on the MitoTimer red:green ratio in DAergic projections to the SOG (F_1,53_=12.38, p<0.001, three-way ANOVA). There was also a significant increase in the MitoTimer red:green ratio in male RNAi flies after 3% H_2_O_2_ exposure in DAergic projections to the FSB (H_2_O_2_ × sex × RNAi interaction: F_1,53_=12.84, p<0.001, three-way ANOVA; p<0.001, Bonferroni multiple comparisons test). N=6-10 per group. *p<0.05, ***p<0.001, ^#^p<0.05 versus male RNAi control; all data are plotted as Mean±SEM.

**Figure S4.**
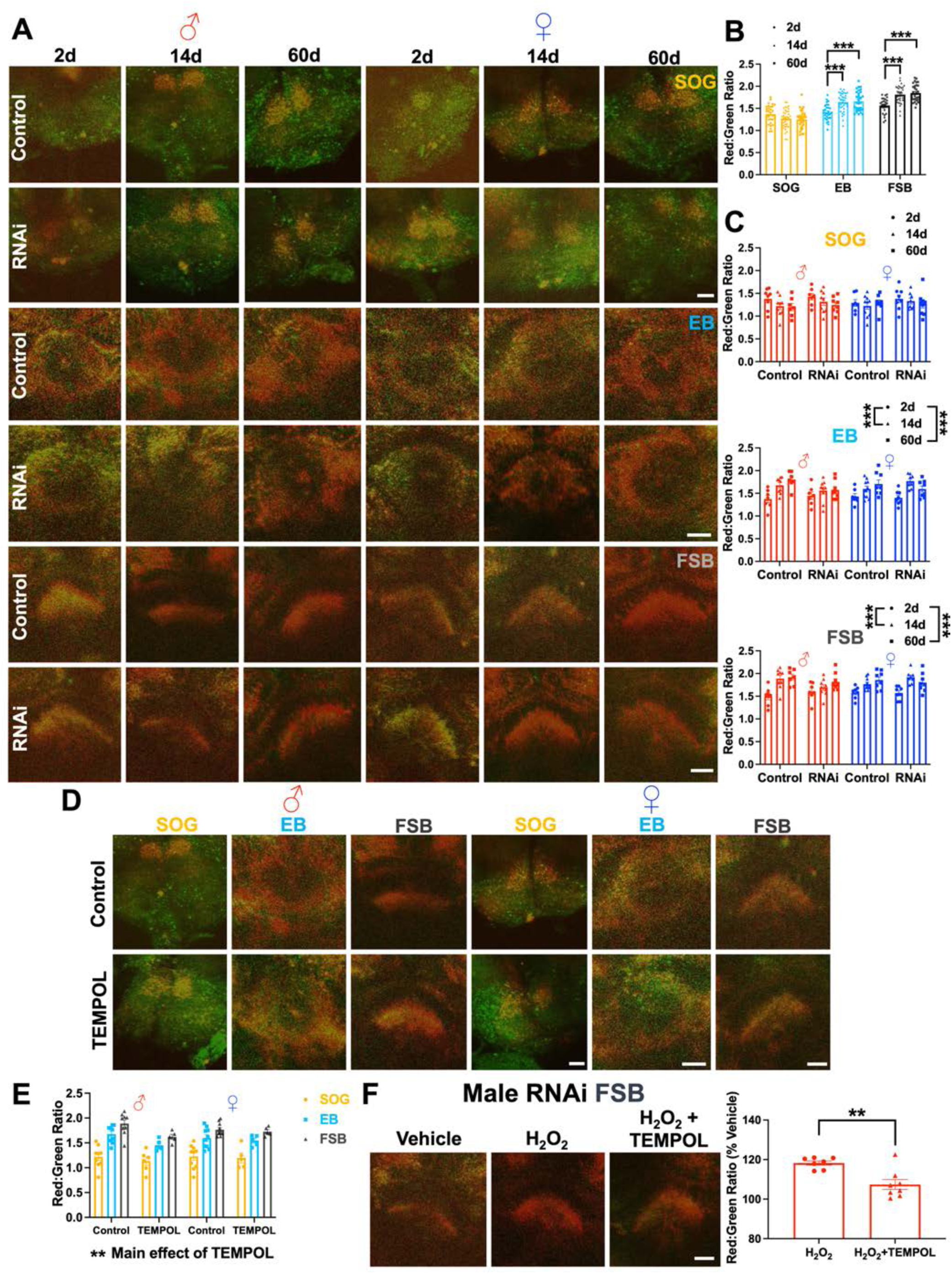
Impacts of aging and TEMPOL on mitochondrial oxidative stress in DA neurons, related to Figure 3. **(A)** Representative projection images of MitoTimer-labeled DAergic projections to the SOG, EB and FSB (with green and red channels overlaid) in brains of male and female DA neuron dVGLUT RNAi (RNAi) and *LexA* RNAi (Control) flies across aging (2d, 14d, 60d post-eclosion). Scale bars=25μm. **(B)** Quantification of the MitoTimer red:green ratio showed regional differences that significantly increased with age (age × brain region interaction: F_4,281_=11.5, p<0.001, two-way ANOVA). Bars display merged data of both sexes and genotypes. **(C)** Age-related increases in MitoTimer red:green ratio were observed in the EB (main effect of age: F_2,84_=15.51, p<0.001, three-way ANOVA) and FSB (main effect of age: F_2,84_=22.34, p<0.001, three-way ANOVA), but not in the SOG (p>0.05, three-way ANOVA). N=7-10 per group. **(D)** Representative projection images of MitoTimer-labeled DAergic projections to the SOG, EB and FSB (with green and red channels overlaid) from adult control flies (14d post-eclosion) exposed to either TEMPOL (3mM, 5d) or vehicle (0mM, 5d). Scale bars=25μm. **(E)** TEMPOL significantly decreased MitoTimer red:green ratio in Control flies (main effect of TEMPOL: F_1,76_=7.79, p=0.0066, three-way ANOVA). N=5-10 per group. **(F)** Representative projection images of MitoTimer-labeled DAergic projections to the FSB. Accompanying quantification of the MitoTimer red:green ratios showed that co-treatment of H_2_O_2_ (3%, 2d) with TEMPOL (3mM, 2d) significantly reduced H_2_O_2_-induced increases in MitoTimer red:green ratio in male RNAi flies (p=0.0024, unpaired t-test). Scale bar=25μm. N=7-8 per group. *p<0.05, **p<0.01, ***p<0.001; all data are plotted as Mean±SEM.

**Figure S5.**
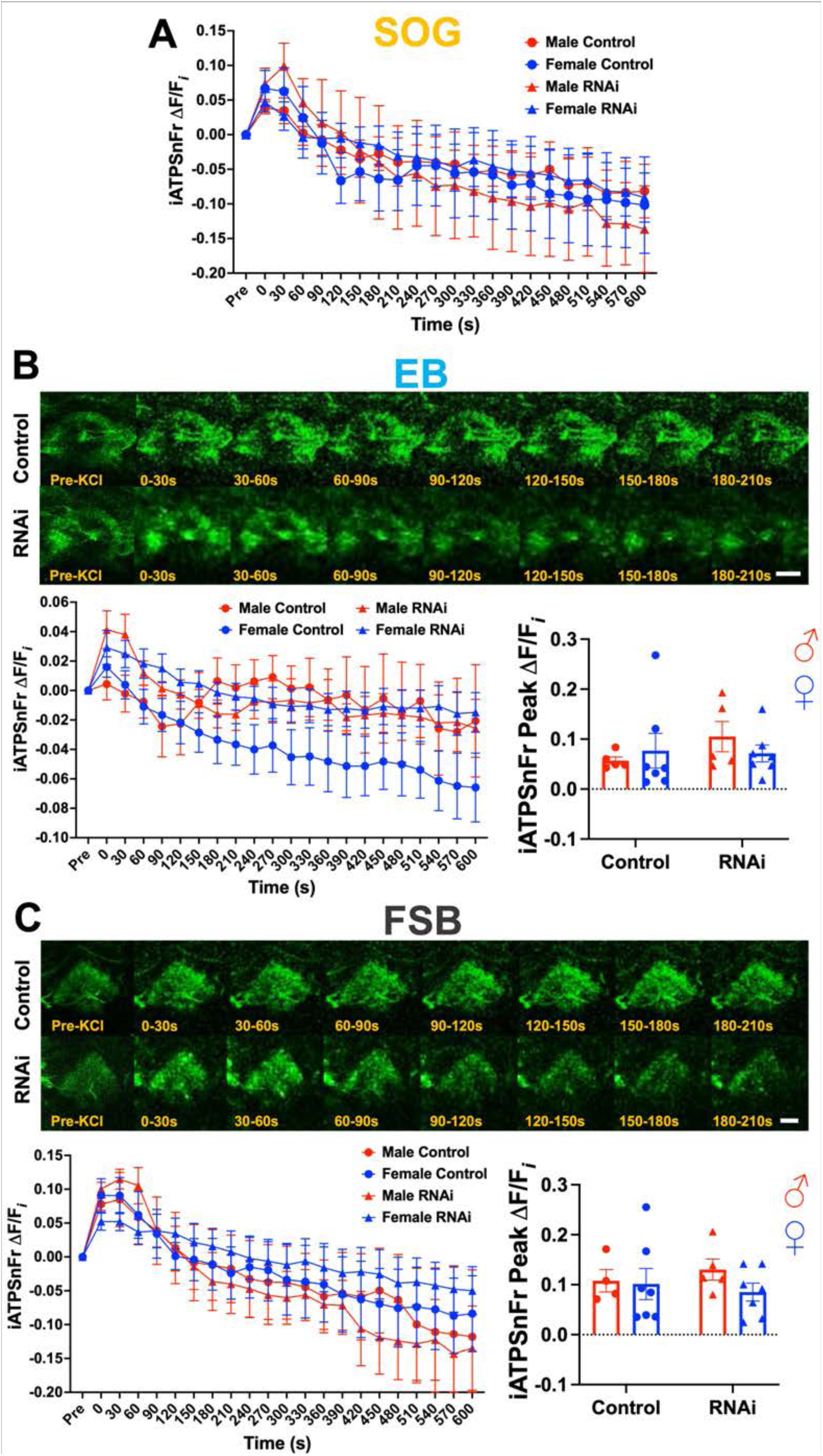
Neuronal depolarization increases intracellular ATP levels in DAergic projections to the EB and FSB, related to Figure 4. **(A)** Quantification of TH-driven iATPSnFr fluorescence changes (ΔF/F*_i_*) in the SOG before and after 40mM KCl-induced depolarization. **(B)** Representative images of TH-driven iATPSnFR fluorescence in DAergic projections to the EB before and after 40mM KCl-induced depolarization. There was no significant effect of DA neuron dVGLUT RNAi on iATPSnFR ΔF/F*_i_* in response to 40mM KCl-induced depolarization (p>0.05, two-way ANOVA). Scale bar=25μm. N=5-7 per group. **(C)** Representative projection images of TH-driven iATPSnFR fluorescence in DAergic projections to the FSB before and after 40mM KCl-induced depolarization. There was no effect of dVGLUT RNAi on iATPSnFR ΔF/F*_i_* during 40mM KCl depolarization (p>0.05, two-way ANOVA). Scale bar=25μm. N=4-7 per group. All data are plotted as Mean±SEM.

**Figure S6.**
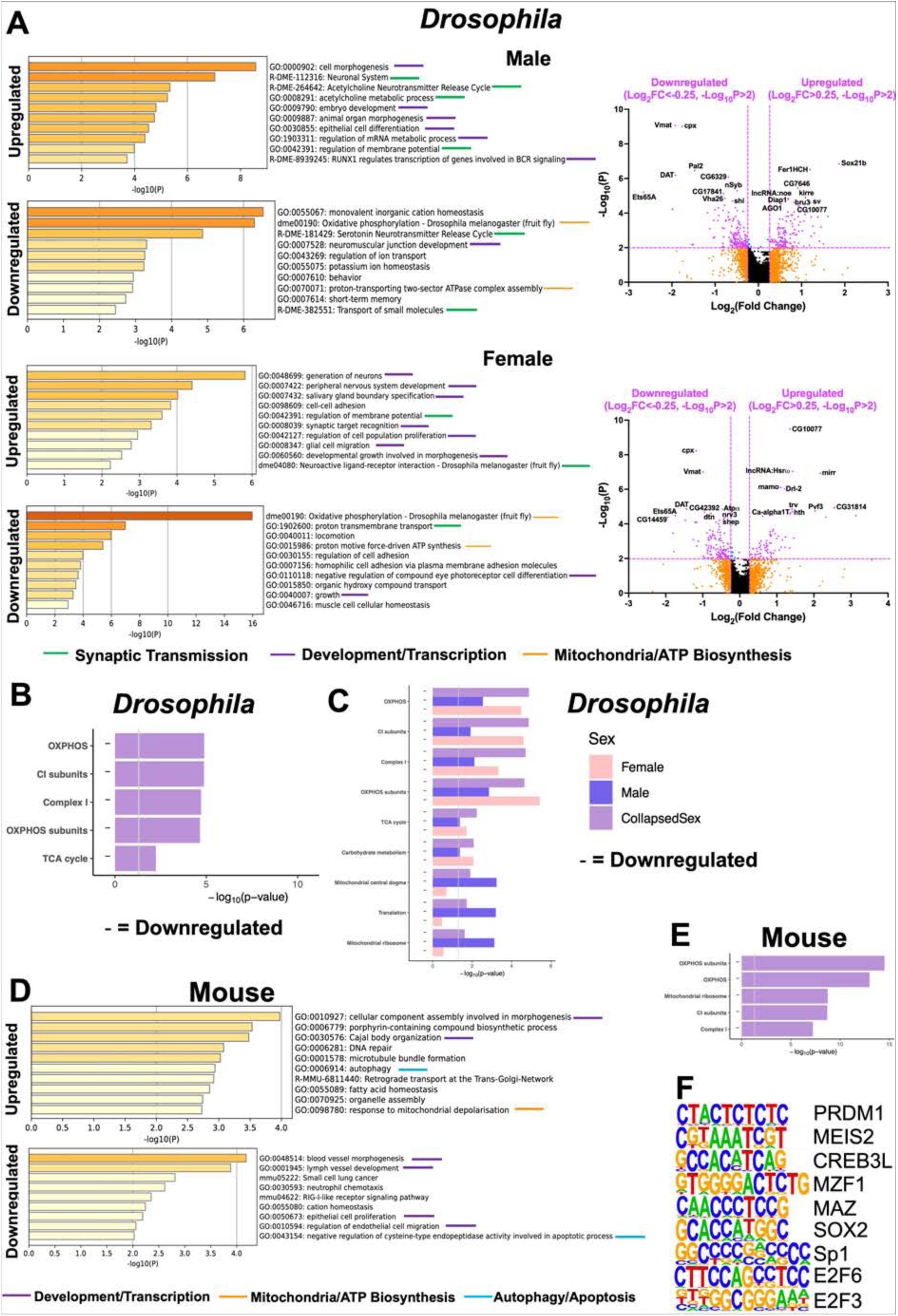
Sex-specific and mitochondria-focused RNAseq analyses, related to Figure 5. **(A)** Gene ontology (GO) enrichment analyses of RNAseq data and volcano plots showing differential expression of genes in dVGLUT^+^ DA neurons compared to dVGLUT^−^ DA neurons in male and female *Drosophila*. GO analyses showed similar pathway enrichments in male and female TH^+^/dVGLUT^+^ DA neurons, with stronger upregulation of synaptic transmission pathway genes in males and stronger downregulation of oxidative phosphorylation pathway genes in females. **(B)** Mitochondria-focused analyses showed genes involved in oxidative phosphorylation and mitochondrial respiratory complex I were downregulated in *Drosophila* dVGLUT^+^ DA neurons. **(C)** Downregulation of mitochondrial genes was especially prominent in female flies. **(D)** GO analyses of mouse RNAseq data showing upregulated and downregulated pathways based on differential gene expression in TH^+^/VGLUT2^+^ DA neurons. **(E)** Genes involved in oxidative phosphorylation and mitochondria respiratory complex I were downregulated in mouse TH^+^/VGLUT2^+^ DA neurons, analogous to *Drosophila*. **(F)** Hypergeometric Optimization of Motif EnRichment (HOMER) analysis identified differentially enriched transcription factors and their respective motifs in mouse TH^+^/VGLUT2^+^ DA neurons.

**Figure S7.**
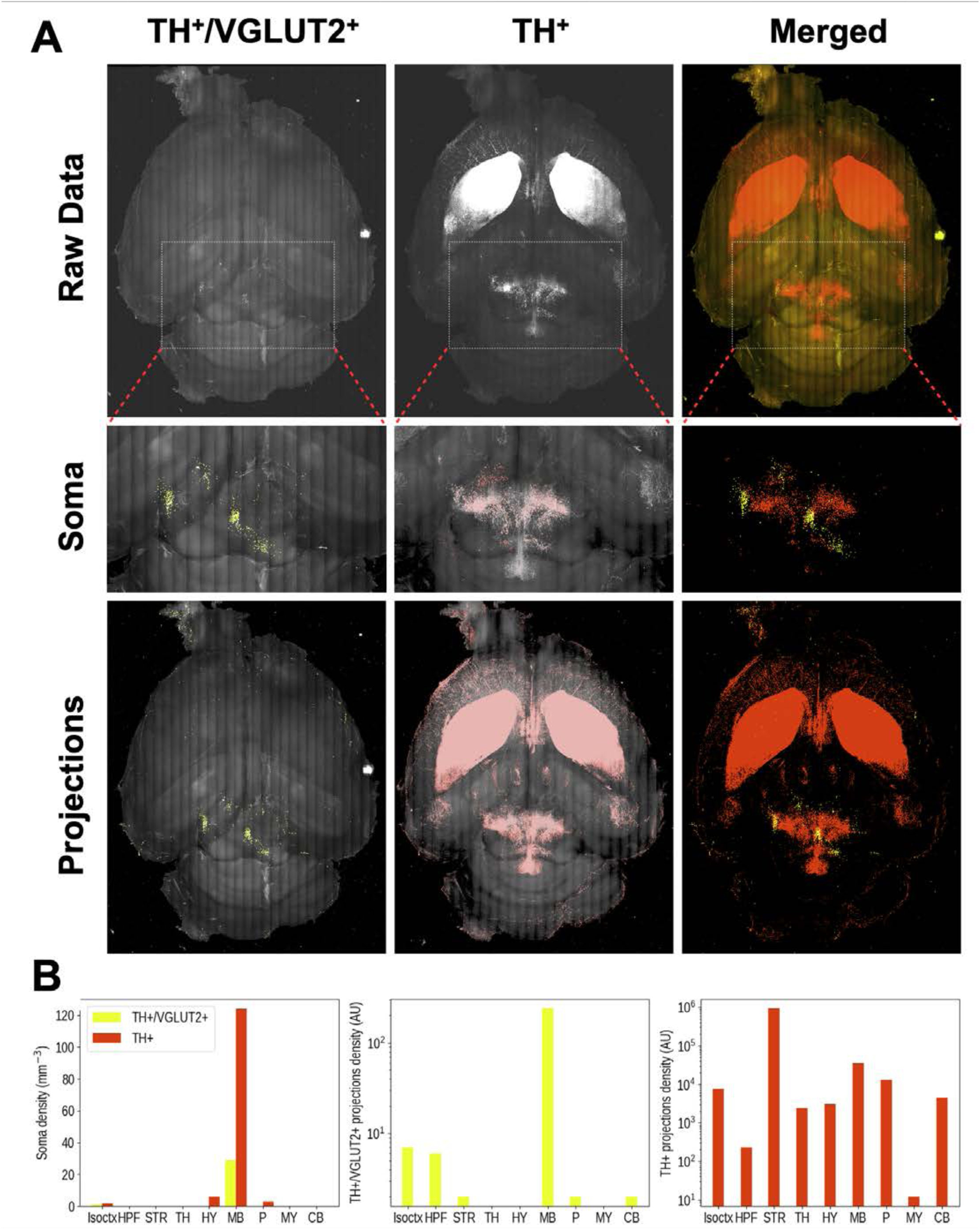
Intersectional genetic imaging of TH^+^/VGLUT2^+^ DA neurons in mouse brain via INTRSECT2.0, related to Figure 5, Video S1, Video S2. **(A, B)** INTRSECT2.0 intersectional genetic labeling of the TH^+^/VGLUT2^+^ subpopulation of VTA DA neurons in whole cleared mouse brain via multicolor ribbon scanning confocal microscopy. TH^+^/VGLUT2^+^ DA neurons were EYFP-labeled (in yellow), while TH^+^-only DA neurons were mCherry-labeled (in red). **(A) Top Panel:** A 2D projection overview of the raw imaging data according to the individual TH^+^/VGLUT2^+^ and TH+-only channels alongside the merged channel. The white box indicates the midbrain region. **Middle Panel:** Magnified views of the midbrain, highlighting INTRSECT2.0-labeled cell bodies. **Bottom Panel:** 2D projection images highlighting neuronal projections of the INTERSECT2.0-labeled cell bodies throughout the brain. **(B) Left panel:** Quantification of INTRSECT2.0-labeled TH^+^/VGLUT2^+^ (yellow bars) and TH^+^-only (red bars) cell bodies. **Middle panel:** Quantification of TH^+^/VGLUT2^+^ (yellow bars) projections according to brain region. **Right panel:** Quantification of TH^+^ (red bars) projections according to brain region. Abbreviations: Isoctx (Isocortex), HPF (hippocampal formation), STR (striatum), TH (thalamus), HY (hypothalamus), MB (midbrain), P (Pons), MY (myelencephalon), CB (cerebellum).

**Figure S8.**
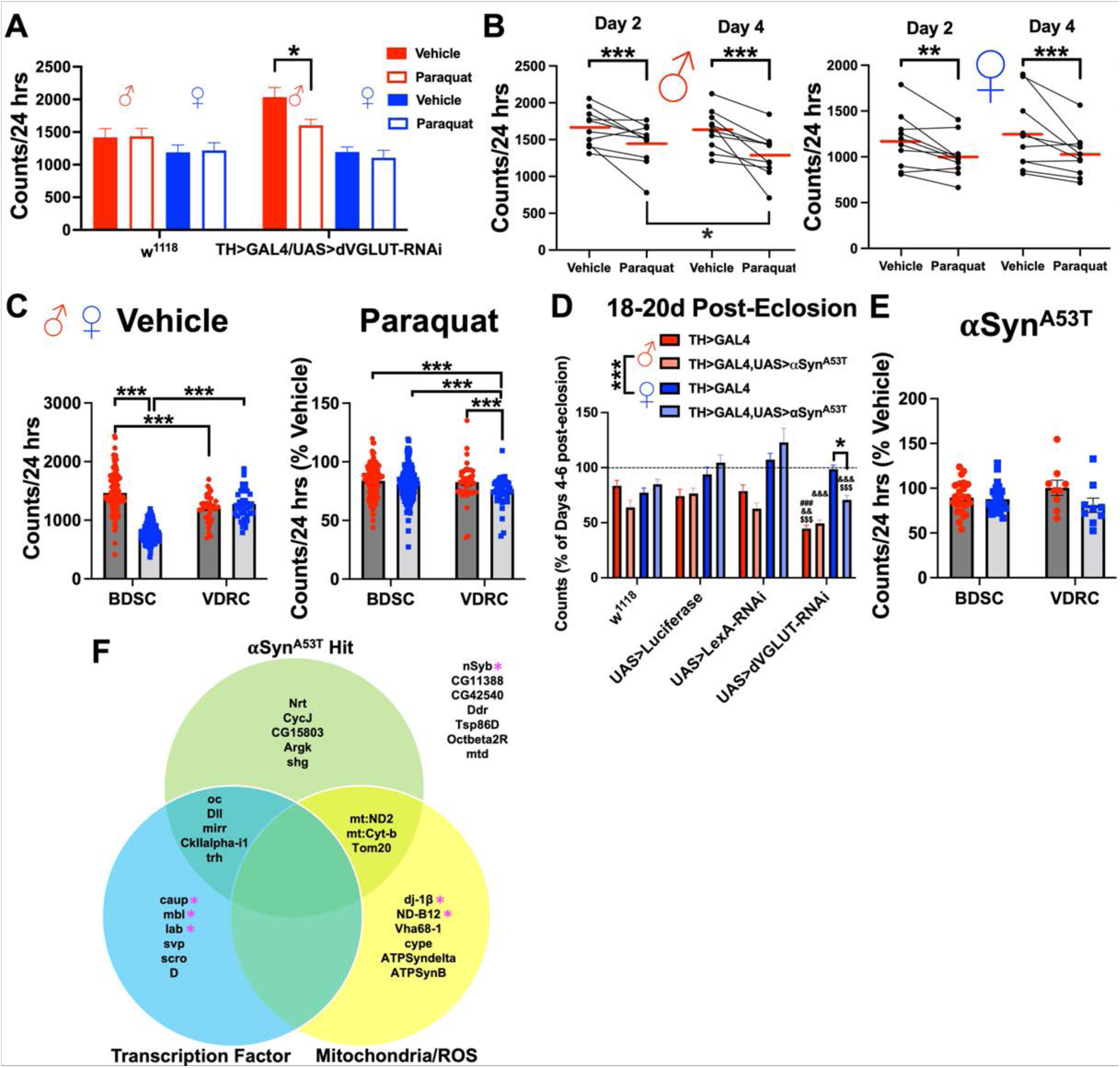
Genetic screen control experiments and survival curves, related to Figure 6. **(A)** Locomotion in adult male and female control w^1118^ flies (N=16-21 flies per group) compared to TH-driven dVGLUT RNAi (TH>GAL4/UAS>dVGLUT-RNAi) flies (N=15-16 flies per group) during days 2-4 of either vehicle or 10mM paraquat exposure. A sample size of 16 flies per group was sufficient to detect locomotor vulnerability to paraquat (p=0.048, Bonferroni multiple comparisons test). **(B)** Locomotion of the 10 control lines tested at day 2 versus day 4 of either vehicle or 10mM paraquat exposure; the respective mean locomotor counts over a 24h period are displayed for each line. There was a significant decrease in locomotion at day 4 of paraquat exposure compared to day 2 in male control lines (p=0.031, Bonferroni multiple comparisons test), but not in female control lines (p>0.05, Bonferroni multiple comparisons test). N=10 control lines per group, 15-33 flies per genotype. **(C)** Comparison of locomotion during days 2-4 of paraquat or vehicle exposure of all analyzed TH-driven Bloomington *Drosophila* Stock Center (BDSC) RNAi lines and Vienna *Drosophila* Resource Center (VDRC) RNAi lines. There was a significant effect of fly repository source (Vehicle: F_1,366_=10.32, p=0.0014; Paraquat: F_1,366_=7.53, p=0.0064, three-way ANOVA), but no interaction between line source and presence of RNAi (p>0.05, three-way ANOVA). N=39-138 RNAi lines per group, 3-37 flies per each line. **(D)** Locomotion of adult TH-driven dVGLUT RNAi flies (18-20d post-eclosion) alongside three control lines ± TH-driven αSyn^A53T^ expression. Results are represented relative to early adult locomotion (4-6d post-eclosion). Males were more vulnerable to age-related decreases in locomotion than females (F_1,398_=93.61, p<0.001, three-way ANOVA). A three-way interaction was also observed (F_3,398_=5.75, p<0.001). DA neuron dVGLUT RNAi knockdown rendered male flies more vulnerable to age-related locomotor decreases, while female dVGLUT RNAi flies only became vulnerable with αSyn^A53T^ overexpression in DA neurons. *p<0.05, ^###^p<0.001 versus w^1118^, ^&&^p<0.01 versus UAS>Luciferase, ^&&&^p<0.001 versus UAS>Luciferase, ^$$$^p<0.001 versus UAS>LexA-RNAi (Bonferroni multiple comparisons test). N=15-43 flies per group. **(E)** Locomotion between RNAi fly lines in response to TH-driven αSyn^A53T^ expression. There were no significant effects of fly repository source on locomotor response to DA neuron αSyn^A53T^ expression in either control or RNAi lines (p>0.05, two-way ANOVA). N=9-24 lines per group, 12-39 flies per line. **(F)** Venn diagram of candidate hits tested in the secondary αSyn^A53T^ screen; the screen candidates are divided according to their outcome in the secondary screen and their biological functions (*i.e*., transcription factors, mitochondrial function/ROS). Asterisks denote significant hits in the secondary αSyn^A53T^ screen after correcting for fly repository source (BDSC, VDRC) but prior to false discovery rate adjustment (p<0.05, q>0.05). *p<0.05, **p<0.01, ***p<0.001; all data are plotted as Mean±SEM.

## References

1. Dorsey, E.R., Sherer, T., Okun, M.S., and Bloem, B.R. (2018). The Emerging Evidence of the Parkinson Pandemic. Journal of Parkinson’s disease 8, S3–s8. 10.3233/jpd-181474.

2. Sulzer, D., and Surmeier, D.J. (2013). Neuronal vulnerability, pathogenesis, and Parkinson’s disease. Movement disorders : official journal of the Movement Disorder Society 28, 715–724. 10.1002/mds.25187.

3. Surmeier, D.J., Obeso, J.A., and Halliday, G.M. (2017). Selective neuronal vulnerability in Parkinson disease. Nat Rev Neurosci 18, 101–113. 10.1038/nrn.2016.178.

4. Hirsch, E., Graybiel, A.M., and Agid, Y.A. (1988). Melanized dopaminergic neurons are differentially susceptible to degeneration in Parkinson’s disease. Nature 334, 345–348. 10.1038/334345a0.

5. Damier, P., Hirsch, E.C., Agid, Y., and Graybiel, A.M. (1999). The substantia nigra of the human brain. II. Patterns of loss of dopamine-containing neurons in Parkinson’s disease. Brain 122 ( *Pt 8*), 1437–1448.

6. Schneider, J.S., Yuwiler, A., and Markham, C.H. (1987). Selective loss of subpopulations of ventral mesencephalic dopaminergic neurons in the monkey following exposure to MPTP. Brain Res 411, 144–150.

7. Jackson-Lewis, V., Jakowec, M., Burke, R.E., and Przedborski, S. (1995). Time course and morphology of dopaminergic neuronal death caused by the neurotoxin 1-methyl-4-phenyl-1,2,3,6-tetrahydropyridine. Neurodegeneration : a journal for neurodegenerative disorders, neuroprotection, and neuroregeneration 4, 257–269.

8. Steinkellner, T., Zell, V., Farino, Z.J., Sonders, M.S., Villeneuve, M., Freyberg, R.J., Przedborski, S., Lu, W., Freyberg, Z., and Hnasko, T.S. (2018). Role for VGLUT2 in selective vulnerability of midbrain dopamine neurons. J Clin Invest 128, 774–788. 10.1172/jci95795.

9. Alberico, S.L., Cassell, M.D., and Narayanan, N.S. (2015). The Vulnerable Ventral Tegmental Area in Parkinson’s Disease. Basal Ganglia 5, 51–55. 10.1016/j.baga.2015.06.001.

10. Wong, K.K., Muller, M.L., Kuwabara, H., Studenski, S.A., and Bohnen, N.I. (2012). Gender differences in nigrostriatal dopaminergic innervation are present at young-to-middle but not at older age in normal adults. J Clin Neurosci 19, 183–184. 10.1016/j.jocn.2011.05.013.

11. Jurado-Coronel, J.C., Cabezas, R., Ávila Rodríguez, M.F., Echeverria, V., García-Segura, L.M., and Barreto, G.E. (2018). Sex differences in Parkinson’s disease: Features on clinical symptoms, treatment outcome, sexual hormones and genetics. Frontiers in neuroendocrinology 50, 18–30. 10.1016/j.yfrne.2017.09.002.

12. Haaxma, C.A., Bloem, B.R., Borm, G.F., Oyen, W.J., Leenders, K.L., Eshuis, S., Booij, J., Dluzen, D.E., and Horstink, M.W. (2007). Gender differences in Parkinson’s disease. J Neurol Neurosurg Psychiatry 78, 819–824. 10.1136/jnnp.2006.103788.

13. Kaasinen, V., Joutsa, J., Noponen, T., Johansson, J., and Seppanen, M. (2015). Effects of aging and gender on striatal and extrastriatal [123I]FP-CIT binding in Parkinson’s disease. Neurobiol Aging 36, 1757–1763. 10.1016/j.neurobiolaging.2015.01.016.

14. Wooten, G.F., Currie, L.J., Bovbjerg, V.E., Lee, J.K., and Patrie, J. (2004). Are men at greater risk for Parkinson’s disease than women? J Neurol Neurosurg Psychiatry 75, 637–639. 10.1136/jnnp.2003.020982.

15. Baldereschi, M., Di Carlo, A., Rocca, W.A., Vanni, P., Maggi, S., Perissinotto, E., Grigoletto, F., Amaducci, L., and Inzitari, D. (2000). Parkinson’s disease and parkinsonism in a longitudinal study: two-fold higher incidence in men. ILSA Working Group. Italian Longitudinal Study on Aging. Neurology 55, 1358–1363. 10.1212/wnl.55.9.1358.

16. Lubomski, M., Louise Rushworth, R., Lee, W., Bertram, K.L., and Williams, D.R. (2014). Sex differences in Parkinson’s disease. Journal of clinical neuroscience : official journal of the Neurosurgical Society of Australasia 21, 1503–1506. 10.1016/j.jocn.2013.12.016.

17. Swerdlow, R.H., Parker, W.D., Currie, L.J., Bennett, J.P., Harrison, M.B., Trugman, J.M., and Wooten, G.F. (2001). Gender ratio differences between Parkinson’s disease patients and their affected relatives. Parkinsonism Relat Disord 7, 129–133. 10.1016/s1353-8020(00)00029-8.

18. Willis, A.W., Roberts, E., Beck, J.C., Fiske, B., Ross, W., Savica, R., Van Den Eeden, S.K., Tanner, C.M., Marras, C., and Parkinson’s Foundation, P.G. (2022). Incidence of Parkinson disease in North America. NPJ Parkinsons Dis 8, 170. 10.1038/s41531-022-00410-y.

19. De Miranda, B.R., Fazzari, M., Rocha, E.M., Castro, S., and Greenamyre, J.T. (2019). Sex differences in rotenone sensitivity reflect the male-to-female ratio in human Parkinson’s disease incidence. Toxicol Sci. 10.1093/toxsci/kfz082.

20. Dluzen, D.E., McDermott, J.L., and Liu, B. (1996). Estrogen as a neuroprotectant against MPTP-induced neurotoxicity in C57/B1 mice. Neurotoxicol Teratol 18, 603–606.

21. Miller, D.B., Ali, S.F., O’Callaghan, J.P., and Laws, S.C. (1998). The impact of gender and estrogen on striatal dopaminergic neurotoxicity. Ann N Y Acad Sci 844, 153–165.

22. Adamson, A., Buck, S.A., Freyberg, Z., and De Miranda, B.R. (2022). Sex Differences in Dopaminergic Vulnerability to Environmental Toxicants - Implications for Parkinson’s Disease. Curr Environ Health Rep. 10.1007/s40572-022-00380-6.

23. Poulin, J.F., Caronia, G., Hofer, C., Cui, Q., Helm, B., Ramakrishnan, C., Chan, C.S., Dombeck, D.A., Deisseroth, K., and Awatramani, R. (2018). Mapping projections of molecularly defined dopamine neuron subtypes using intersectional genetic approaches. Nat Neurosci 21, 1260–1271. 10.1038/s41593-018-0203-4.

24. Yamaguchi, T., Wang, H.L., Li, X., Ng, T.H., and Morales, M. (2011). Mesocorticolimbic glutamatergic pathway. J Neurosci 31, 8476–8490. 10.1523/JNEUROSCI.1598-11.2011.

25. Yamaguchi, T., Wang, H.L., and Morales, M. (2013). Glutamate neurons in the substantia nigra compacta and retrorubral field. Eur J Neurosci 38, 3602–3610. 10.1111/ejn.12359.

26. Yamaguchi, T., Qi, J., Wang, H.L., Zhang, S., and Morales, M. (2015). Glutamatergic and dopaminergic neurons in the mouse ventral tegmental area. The European journal of neuroscience 41, 760–772. 10.1111/ejn.12818.

27. Mingote, S., Amsellem, A., Kempf, A., Rayport, S., and Chuhma, N. (2019). Dopamine-glutamate neuron projections to the nucleus accumbens medial shell and behavioral switching. Neurochemistry international 129, 104482. 10.1016/j.neuint.2019.104482.

28. Chuhma, N., Mingote, S., Yetnikoff, L., Kalmbach, A., Ma, T., Ztaou, S., Sienna, A.C., Tepler, S., Poulin, J.F., Ansorge, M., et al. (2018). Dopamine neuron glutamate cotransmission evokes a delayed excitation in lateral dorsal striatal cholinergic interneurons. eLife 7. 10.7554/eLife.39786.

29. Maingay, M., Romero-Ramos, M., Carta, M., and Kirik, D. (2006). Ventral tegmental area dopamine neurons are resistant to human mutant alpha-synuclein overexpression. Neurobiology of disease 23, 522–532. 10.1016/j.nbd.2006.04.007.

30. Betarbet, R., Sherer, T.B., MacKenzie, G., Garcia-Osuna, M., Panov, A.V., and Greenamyre, J.T. (2000). Chronic systemic pesticide exposure reproduces features of Parkinson’s disease. Nat Neurosci 3, 1301–1306. 10.1038/81834.

31. Gibb, W.R., and Lees, A.J. (1991). Anatomy, pigmentation, ventral and dorsal subpopulations of the substantia nigra, and differential cell death in Parkinson’s disease. J Neurol Neurosurg Psychiatry 54, 388–396. 10.1136/jnnp.54.5.388.

32. Herrero, M.T., Hirsch, E.C., Kastner, A., Ruberg, M., Luquin, M.R., Laguna, J., Javoy-Agid, F., Obeso, J.A., and Agid, Y. (1993). Does neuromelanin contribute to the vulnerability of catecholaminergic neurons in monkeys intoxicated with MPTP? Neuroscience 56, 499–511. 10.1016/0306-4522(93)90349-k.

33. Shen, H., Marino, R.A.M., McDevitt, R.A., Bi, G.H., Chen, K., Madeo, G., Lee, P.T., Liang, Y., De Biase, L.M., Su, T.P., et al. (2018). Genetic deletion of vesicular glutamate transporter in dopamine neurons increases vulnerability to MPTP-induced neurotoxicity in mice. Proc Natl Acad Sci U S A 115, E11532–e11541. 10.1073/pnas.1800886115.

34. Dal Bo, G., Berube-Carriere, N., Mendez, J.A., Leo, D., Riad, M., Descarries, L., Levesque, D., and Trudeau, L.E. (2008). Enhanced glutamatergic phenotype of mesencephalic dopamine neurons after neonatal 6-hydroxydopamine lesion. Neuroscience 156, 59–70. 10.1016/j.neuroscience.2008.07.032.

35. Steinkellner, T., Conrad, W.S., Kovacs, I., Rissman, R.A., Lee, E.B., Trojanowski, J.Q., Freyberg, Z., Roy, S., Luk, K.C., Lee, V.M., and Hnasko, T.S. (2022). Dopamine neurons exhibit emergent glutamatergic identity in Parkinson’s disease. Brain 145, 879–886. 10.1093/brain/awab373.

36. Buck, S.A., Steinkellner, T., Aslanoglou, D., Villeneuve, M., Bhatte, S.H., Childers, V.C., Rubin, S.A., De Miranda, B.R., O’Leary, E.I., Neureiter, E.G., et al. (2021). Vesicular glutamate transporter modulates sex differences in dopamine neuron vulnerability to age-related neurodegeneration. Aging Cell 20, e13365. 10.1111/acel.13365.

37. Goldman, S.M. (2014). Environmental toxins and Parkinson’s disease. Annu Rev Pharmacol Toxicol 54, 141–164. 10.1146/annurev-pharmtox-011613-135937.

38. Tanner, C.M., Ross, G.W., Jewell, S.A., Hauser, R.A., Jankovic, J., Factor, S.A., Bressman, S., Deligtisch, A., Marras, C., Lyons, K.E., et al. (2009). Occupation and risk of parkinsonism: a multicenter case-control study. Archives of neurology 66, 1106–1113. 10.1001/archneurol.2009.195.

39. McCormack, A.L., Thiruchelvam, M., Manning-Bog, A.B., Thiffault, C., Langston, J.W., Cory-Slechta, D.A., and Di Monte, D.A. (2002). Environmental risk factors and Parkinson’s disease: selective degeneration of nigral dopaminergic neurons caused by the herbicide paraquat. Neurobiology of disease 10, 119–127.

40. Buck, S.A., Erickson-Oberg, M.Q., Bhatte, S.H., McKellar, C.D., Ramanathan, V.P., Rubin, S.A., and Freyberg, Z. (2022). Roles of VGLUT2 and Dopamine/Glutamate Co-Transmission in Selective Vulnerability to Dopamine Neurodegeneration. ACS chemical neuroscience 13, 187–193. 10.1021/acschemneuro.1c00741.

41. Castello, P.R., Drechsel, D.A., and Patel, M. (2007). Mitochondria are a major source of paraquat-induced reactive oxygen species production in the brain. J Biol Chem 282, 14186–14193. 10.1074/jbc.M700827200.

42. Cassar, M., Issa, A.R., Riemensperger, T., Petitgas, C., Rival, T., Coulom, H., Iche-Torres, M., Han, K.A., and Birman, S. (2015). A dopamine receptor contributes to paraquat-induced neurotoxicity in Drosophila. Hum Mol Genet 24, 197–212. 10.1093/hmg/ddu430.

43. Rappold, P.M., Cui, M., Chesser, A.S., Tibbett, J., Grima, J.C., Duan, L., Sen, N., Javitch, J.A., and Tieu, K. (2011). Paraquat neurotoxicity is mediated by the dopamine transporter and organic cation transporter-3. Proc Natl Acad Sci U S A 108, 20766–20771. 10.1073/pnas.1115141108.

44. Aryal, B., and Lee, Y. (2019). Disease model organism for Parkinson disease: Drosophila melanogaster. BMB Rep 52, 250–258. 10.5483/BMBRep.2019.52.4.204.

45. Martin, C.A., Barajas, A., Lawless, G., Lawal, H.O., Assani, K., Lumintang, Y.P., Nunez, V., and Krantz, D.E. (2014). Synergistic effects on dopamine cell death in a Drosophila model of chronic toxin exposure. Neurotoxicology 44, 344–351. 10.1016/j.neuro.2014.08.005.

46. Chaudhuri, A., Bowling, K., Funderburk, C., Lawal, H., Inamdar, A., Wang, Z., and O’Donnell, J.M. (2007). Interaction of genetic and environmental factors in a Drosophila parkinsonism model. J Neurosci 27, 2457–2467. 10.1523/jneurosci.4239-06.2007.

47. Mao, Z., and Davis, R.L. (2009). Eight different types of dopaminergic neurons innervate the Drosophila mushroom body neuropil: anatomical and physiological heterogeneity. Frontiers in neural circuits 3, 5. 10.3389/neuro.04.005.2009.

48. Ma, D., Herndon, N., Le, J.Q., Abruzzi, K.C., Zinn, K., and Rosbash, M. (2023). Neural connectivity molecules best identify the heterogeneous clock and dopaminergic cell types in the Drosophila adult brain. Sci Adv 9, eade8500. 10.1126/sciadv.ade8500.

49. Lawal, H.O., Chang, H.Y., Terrell, A.N., Brooks, E.S., Pulido, D., Simon, A.F., and Krantz, D.E. (2010). The Drosophila vesicular monoamine transporter reduces pesticide-induced loss of dopaminergic neurons. Neurobiology of disease 40, 102–112. 10.1016/j.nbd.2010.05.008.

50. Cohen, G.M., and d’Arcy Doherty, M. (1987). Free radical mediated cell toxicity by redox cycling chemicals. Br J Cancer Suppl 8, 46–52.

51. Tryphena, K.P., Nikhil, U.S., Pinjala, P., Srivastava, S., Singh, S.B., and Khatri, D.K. (2023). Mitochondrial Complex I as a Pathologic and Therapeutic Target for Parkinson’s Disease. ACS Chem Neurosci. 10.1021/acschemneuro.2c00819.

52. Grunewald, A., Kumar, K.R., and Sue, C.M. (2019). New insights into the complex role of mitochondria in Parkinson’s disease. Prog Neurobiol 177, 73–93. 10.1016/j.pneurobio.2018.09.003.

53. Hernandez, G., Thornton, C., Stotland, A., Lui, D., Sin, J., Ramil, J., Magee, N., Andres, A., Quarato, G., Carreira, R.S., et al. (2013). MitoTimer: a novel tool for monitoring mitochondrial turnover. Autophagy 9, 1852–1861. 10.4161/auto.26501.

54. Laker, R.C., Xu, P., Ryall, K.A., Sujkowski, A., Kenwood, B.M., Chain, K.H., Zhang, M., Royal, M.A., Hoehn, K.L., Driscoll, M., et al. (2014). A novel MitoTimer reporter gene for mitochondrial content, structure, stress, and damage in vivo. J Biol Chem 289, 12005–12015. 10.1074/jbc.M113.530527.

55. Nassel, D.R., and Elekes, K. (1992). Aminergic neurons in the brain of blowflies and Drosophila: dopamine- and tyrosine hydroxylase-immunoreactive neurons and their relationship with putative histaminergic neurons. Cell Tissue Res 267, 147–167. 10.1007/bf00318701.

56. Marella, S., Mann, K., and Scott, K. (2012). Dopaminergic modulation of sucrose acceptance behavior in Drosophila. Neuron 73, 941–950. 10.1016/j.neuron.2011.12.032 [doi] S0896-6273(12)00082-7 [pii].

57. Trisal, S., Aranha, M., Chodankar, A., VijayRaghavan, K., and Ramaswami, M. (2022). A Drosophila Circuit for Habituation Override. J Neurosci 42, 2930–2941. 10.1523/JNEUROSCI.1842-21.2022.

58. Kahsai, L., Carlsson, M.A., Winther, A.M., and Nassel, D.R. (2012). Distribution of metabotropic receptors of serotonin, dopamine, GABA, glutamate, and short neuropeptide F in the central complex of Drosophila. Neuroscience 208, 11–26. 10.1016/j.neuroscience.2012.02.007.

59. Kottler, B., Faville, R., Bridi, J.C., and Hirth, F. (2019). Inverse Control of Turning Behavior by Dopamine D1 Receptor Signaling in Columnar and Ring Neurons of the Central Complex in Drosophila. Curr Biol 29, 567–577 e566. 10.1016/j.cub.2019.01.017.

60. Dumitrescu, E., Copeland, J.M., and Venton, B.J. (2023). Parkin Knockdown Modulates Dopamine Release in the Central Complex, but Not the Mushroom Body Heel, of Aging Drosophila. ACS chemical neuroscience 14, 198–208. 10.1021/acschemneuro.2c00277.

61. Frighetto, G., Zordan, M.A., Castiello, U., Megighian, A., and Martin, J.R. (2022). Dopamine Modulation of Drosophila Ellipsoid Body Neurons, a Nod to the Mammalian Basal Ganglia. Frontiers in physiology 13, 849142. 10.3389/fphys.2022.849142.

62. Roy, Z., Bansal, R., Siddiqui, L., and Chaudhary, N. (2023). Understanding the Role of Free Radicals and Antioxidant Enzymes in Human Diseases. Curr Pharm Biotechnol 24, 1265–1276. 10.2174/1389201024666221121160822.

63. Mailloux, R.J. (2021). An update on methods and approaches for interrogating mitochondrial reactive oxygen species production. Redox biology 45, 102044. 10.1016/j.redox.2021.102044.

64. Wilcox, C.S. (2010). Effects of tempol and redox-cycling nitroxides in models of oxidative stress. Pharmacol Ther 126, 119–145. 10.1016/j.pharmthera.2010.01.003.

65. van Hameren, G., Campbell, G., Deck, M., Berthelot, J., Gautier, B., Quintana, P., Chrast, R., and Tricaud, N. (2019). In vivo real-time dynamics of ATP and ROS production in axonal mitochondria show decoupling in mouse models of peripheral neuropathies. Acta Neuropathol Commun 7, 86. 10.1186/s40478-019-0740-4.

66. Murphy, M.P. (2009). How mitochondria produce reactive oxygen species. The Biochemical journal 417, 1–13. 10.1042/bj20081386.

67. Lobas, M.A., Tao, R., Nagai, J., Kronschlager, M.T., Borden, P.M., Marvin, J.S., Looger, L.L., and Khakh, B.S. (2019). A genetically encoded single-wavelength sensor for imaging cytosolic and cell surface ATP. Nat Commun 10, 711. 10.1038/s41467-019-08441-5.

68. Mann, K., Deny, S., Ganguli, S., and Clandinin, T.R. (2021). Coupling of activity, metabolism and behaviour across the Drosophila brain. Nature 593, 244–248. 10.1038/s41586-021-03497-0.

69. Treiber, C.D., Dempsey, G., and Waddell, S. (2023). Transcriptomics defines extensive heterogeneity and alternative operating modes of a dopaminergic system. Manuscript in preparation.

70. Aguilar, J.I., Dunn, M., Mingote, S., Karam, C.S., Farino, Z.J., Sonders, M.S., Choi, S.J., Grygoruk, A., Zhang, Y., Cela, C., et al. (2017). Neuronal Depolarization Drives Increased Dopamine Synaptic Vesicle Loading via VGLUT. Neuron 95, 1074–1088.e1077. 10.1016/j.neuron.2017.07.038.

71. Rath, S., Sharma, R., Gupta, R., Ast, T., Chan, C., Durham, T.J., Goodman, R.P., Grabarek, Z., Haas, M.E., Hung, W.H.W., et al. (2021). MitoCarta3.0: an updated mitochondrial proteome now with sub-organelle localization and pathway annotations. Nucleic Acids Res 49, D1541–D1547. 10.1093/nar/gkaa1011.

72. Fenno, L.E., Mattis, J., Ramakrishnan, C., and Deisseroth, K. (2017). A Guide to Creating and Testing New INTRSECT Constructs. Curr Protoc Neurosci 80, 4 39 31-34 39 24. 10.1002/cpns.30.

73. Wang, Q., Ding, S.L., Li, Y., Royall, J., Feng, D., Lesnar, P., Graddis, N., Naeemi, M., Facer, B., Ho, A., et al. (2020). The Allen Mouse Brain Common Coordinate Framework: A 3D Reference Atlas. Cell 181, 936–953.e920. 10.1016/j.cell.2020.04.007.

74. He, K., Zhang, X., Ren, S., and Sun, J. (2016). Deep residual learning for image recognition. pp. 770–778.

75. Berg, S., Kutra, D., Kroeger, T., Straehle, C.N., Kausler, B.X., Haubold, C., Schiegg, M., Ales, J., Beier, T., Rudy, M., et al. (2019). ilastik: interactive machine learning for (bio)image analysis. Nat Methods 16, 1226–1232. 10.1038/s41592-019-0582-9.

76. Mingote, S., Chuhma, N., Kusnoor, S.V., Field, B., Deutch, A.Y., and Rayport, S. (2015). Functional Connectome Analysis of Dopamine Neuron Glutamatergic Connections in Forebrain Regions. J Neurosci 35, 16259–16271. 10.1523/jneurosci.1674-15.2015.

77. Perkins, L.A., Holderbaum, L., Tao, R., Hu, Y., Sopko, R., McCall, K., Yang-Zhou, D., Flockhart, I., Binari, R., Shim, H.S., et al. (2015). The Transgenic RNAi Project at Harvard Medical School: Resources and Validation. Genetics 201, 843–852. 10.1534/genetics.115.180208.

78. Hague, S., Rogaeva, E., Hernandez, D., Gulick, C., Singleton, A., Hanson, M., Johnson, J., Weiser, R., Gallardo, M., Ravina, B., et al. (2003). Early-onset Parkinson’s disease caused by a compound heterozygous DJ-1 mutation. Ann Neurol 54, 271–274. 10.1002/ana.10663.

79. Di Maio, R., Barrett, P.J., Hoffman, E.K., Barrett, C.W., Zharikov, A., Borah, A., Hu, X., McCoy, J., Chu, C.T., Burton, E.A., et al. (2016). alpha-Synuclein binds to TOM20 and inhibits mitochondrial protein import in Parkinson’s disease. Science translational medicine 8, 342ra378. 10.1126/scitranslmed.aaf3634.

80. Varga, S.J., Qi, C., Podolsky, E., and Lee, D. (2014). A new Drosophila model to study the interaction between genetic and environmental factors in Parkinson’s disease. Brain Res 1583, 277–286. 10.1016/j.brainres.2014.08.021.

81. Abul Khair, S.B., Dhanushkodi, N.R., Ardah, M.T., Chen, W., Yang, Y., and Haque, M.E. (2018). Silencing of Glucocerebrosidase Gene in Drosophila Enhances the Aggregation of Parkinson’s Disease Associated alpha-Synuclein Mutant A53T and Affects Locomotor Activity. Front Neurosci 12, 81. 10.3389/fnins.2018.00081.

82. Menzies, F.M., Yenisetti, S.C., and Min, K.T. (2005). Roles of Drosophila DJ-1 in survival of dopaminergic neurons and oxidative stress. Curr Biol 15, 1578–1582. 10.1016/j.cub.2005.07.036.

83. Lavara-Culebras, E., and Paricio, N. (2007). Drosophila DJ-1 mutants are sensitive to oxidative stress and show reduced lifespan and motor deficits. Gene 400, 158–165. 10.1016/j.gene.2007.06.013.

84. Blanco, J., Pandey, R., Wasser, M., and Udolph, G. (2011). Orthodenticle is necessary for survival of a cluster of clonally related dopaminergic neurons in the Drosophila larval and adult brain. Neural Dev 6, 34. 10.1186/1749-8104-6-34.

85. Kwon, H.J., Heo, J.Y., Shim, J.H., Park, J.H., Seo, K.S., Ryu, M.J., Han, J.S., Shong, M., Son, J.H., and Kweon, G.R. (2011). DJ-1 mediates paraquat-induced dopaminergic neuronal cell death. Toxicol Lett 202, 85–92. 10.1016/j.toxlet.2011.01.018.

86. Yamazaki, D., Maeyama, Y., and Tabata, T. (2023). Combinatory actions of co-transmitters in dopaminergic systems modulate Drosophila olfactory memories. J Neurosci. 10.1523/jneurosci.2152-22.2023.

87. Chen, N., Zhang, Y., Rivera-Rodriguez, E.J., Yu, A.D., Hobin, M., Rosbash, M., and Griffith, L.C. (2023). Widespread posttranscriptional regulation of cotransmission. Sci Adv 9, eadg9836. 10.1126/sciadv.adg9836.

88. Dvořáček, J., Bednářová, A., Krishnan, N., and Kodrík, D. (2022). Dopaminergic mushroom body neurons in Drosophila: Flexibility of neuron identity in a model organism? Neurosci Biobehav Rev 135, 104570. 10.1016/j.neubiorev.2022.104570.

89. Takemura, S.Y., Aso, Y., Hige, T., Wong, A., Lu, Z., Xu, C.S., Rivlin, P.K., Hess, H., Zhao, T., Parag, T., et al. (2017). A connectome of a learning and memory center in the adult Drosophila brain. eLife 6. 10.7554/eLife.26975.

90. Svensson, E., Apergis-Schoute, J., Burnstock, G., Nusbaum, M.P., Parker, D., and Schioth, H.B. (2018). General Principles of Neuronal Co-transmission: Insights From Multiple Model Systems. Frontiers in neural circuits 12, 117. 10.3389/fncir.2018.00117.

91. Hnasko, T.S., and Edwards, R.H. (2012). Neurotransmitter corelease: mechanism and physiological role. Annual review of physiology 74, 225–243. 10.1146/annurev-physiol-020911-153315.

92. Aso, Y., Herb, A., Ogueta, M., Siwanowicz, I., Templier, T., Friedrich, A.B., Ito, K., Scholz, H., and Tanimoto, H. (2012). Three dopamine pathways induce aversive odor memories with different stability. PLoS Genet 8, e1002768. 10.1371/journal.pgen.1002768.

93. Aso, Y., Siwanowicz, I., Bracker, L., Ito, K., Kitamoto, T., and Tanimoto, H. (2010). Specific dopaminergic neurons for the formation of labile aversive memory. Curr Biol 20, 1445–1451. 10.1016/j.cub.2010.06.048.

94. Otto, N., Pleijzier, M.W., Morgan, I.C., Edmondson-Stait, A.J., Heinz, K.J., Stark, I., Dempsey, G., Ito, M., Kapoor, I., Hsu, J., et al. (2020). Input Connectivity Reveals Additional Heterogeneity of Dopaminergic Reinforcement in Drosophila. Curr Biol 30, 3200–3211.e3208. 10.1016/j.cub.2020.05.077.

95. Li, F., Lindsey, J.W., Marin, E.C., Otto, N., Dreher, M., Dempsey, G., Stark, I., Bates, A.S., Pleijzier, M.W., Schlegel, P., et al. (2020). The connectome of the adult Drosophila mushroom body provides insights into function. Elife 9. 10.7554/eLife.62576.

96. Burke, C.J., Huetteroth, W., Owald, D., Perisse, E., Krashes, M.J., Das, G., Gohl, D., Silies, M., Certel, S., and Waddell, S. (2012). Layered reward signalling through octopamine and dopamine in Drosophila. Nature 492, 433–437. 10.1038/nature11614.

97. Liu, C., Placais, P.Y., Yamagata, N., Pfeiffer, B.D., Aso, Y., Friedrich, A.B., Siwanowicz, I., Rubin, G.M., Preat, T., and Tanimoto, H. (2012). A subset of dopamine neurons signals reward for odour memory in Drosophila. Nature 488, 512–516. 10.1038/nature11304.

98. Buck, S.A., Miranda, B.R., Logan, R.W., Fish, K.N., Greenamyre, J.T., and Freyberg, Z. (2021). VGLUT2 is a determinant of dopamine neuron resilience in a rotenone model of dopamine neurodegeneration. J Neurosci. 10.1523/jneurosci.2770-20.2021.

99. Reeve, A., Simcox, E., and Turnbull, D. (2014). Ageing and Parkinson’s disease: why is advancing age the biggest risk factor? Ageing Res Rev 14, 19–30. 10.1016/j.arr.2014.01.004.

100. Kenchappa, R.S., Diwakar, L., Annepu, J., and Ravindranath, V. (2004). Estrogen and neuroprotection: higher constitutive expression of glutaredoxin in female mice offers protection against MPTP-mediated neurodegeneration. FASEB J 18, 1102–1104. 10.1096/fj.03-1075fje.

101. Ciron, C., Zheng, L., Bobela, W., Knott, G.W., Leone, T.C., Kelly, D.P., and Schneider, B.L. (2015). PGC-1alpha activity in nigral dopamine neurons determines vulnerability to alpha-synuclein. Acta Neuropathol Commun 3, 16. 10.1186/s40478-015-0200-8.

102. Misiak, M., Beyer, C., and Arnold, S. (2010). Gender-specific role of mitochondria in the vulnerability of 6-hydroxydopamine-treated mesencephalic neurons. Biochim Biophys Acta 1797, 1178–1188. 10.1016/j.bbabio.2010.04.009.

103. Lee, J., Pinares-Garcia, P., Loke, H., Ham, S., Vilain, E., and Harley, V.R. (2019). Sex-specific neuroprotection by inhibition of the Y-chromosome gene, SRY, in experimental Parkinson’s disease. Proc Natl Acad Sci U S A 116, 16577–16582. 10.1073/pnas.1900406116.

104. Pacelli, C., Giguere, N., Bourque, M.J., Levesque, M., Slack, R.S., and Trudeau, L.E. (2015). Elevated Mitochondrial Bioenergetics and Axonal Arborization Size Are Key Contributors to the Vulnerability of Dopamine Neurons. Curr Biol 25, 2349–2360. 10.1016/j.cub.2015.07.050.

105. German, D.C., Manaye, K.F., Sonsalla, P.K., and Brooks, B.A. (1992). Midbrain dopaminergic cell loss in Parkinson’s disease and MPTP-induced parkinsonism: sparing of calbindin-D28k-containing cells. Ann N Y Acad Sci 648, 42–62. 10.1111/j.1749-6632.1992.tb24523.x.

106. Jung, S., Chung, Y., Lee, Y., Lee, Y., Cho, J.W., Shin, E.J., Kim, H.C., and Oh, Y.J. (2019). Buffering of cytosolic calcium plays a neuroprotective role by preserving the autophagy-lysosome pathway during MPP(+)-induced neuronal death. Cell Death Discov 5, 130. 10.1038/s41420-019-0210-6.

107. Liang, C.L., Sinton, C.M., Sonsalla, P.K., and German, D.C. (1996). Midbrain dopaminergic neurons in the mouse that contain calbindin-D28k exhibit reduced vulnerability to MPTP-induced neurodegeneration. Neurodegeneration 5, 313–318. 10.1006/neur.1996.0042.

108. Ricke, K.M., Pass, T., Kimoloi, S., Fahrmann, K., Jungst, C., Schauss, A., Baris, O.R., Aradjanski, M., Trifunovic, A., Eriksson Faelker, T.M., et al. (2020). Mitochondrial Dysfunction Combined with High Calcium Load Leads to Impaired Antioxidant Defense Underlying the Selective Loss of Nigral Dopaminergic Neurons. J Neurosci 40, 1975–1986. 10.1523/JNEUROSCI.1345-19.2019.

109. Kolen, B., Borghans, B., Kortzak, D., Lugo, V., Hannack, C., Guzman, R.E., Ullah, G., and Fahlke, C. (2023). Vesicular glutamate transporters are H(+)-anion exchangers that operate at variable stoichiometry. Nat Commun 14, 2723. 10.1038/s41467-023-38340-9.

110. Blakely, R.D., and Edwards, R.H. (2012). Vesicular and plasma membrane transporters for neurotransmitters. Cold Spring Harb Perspect Biol 4. 10.1101/cshperspect.a005595.

111. Grace, A.A., and Bunney, B.S. (1984). The control of firing pattern in nigral dopamine neurons: burst firing. J Neurosci 4, 2877–2890.

112. Hnasko, T.S., Hjelmstad, G.O., Fields, H.L., and Edwards, R.H. (2012). Ventral tegmental area glutamate neurons: electrophysiological properties and projections. J Neurosci 32, 15076–15085. 10.1523/jneurosci.3128-12.2012.

113. Chuhma, N., Choi, W.Y., Mingote, S., and Rayport, S. (2009). Dopamine neuron glutamate cotransmission: frequency-dependent modulation in the mesoventromedial projection. Neuroscience 164, 1068–1083. 10.1016/j.neuroscience.2009.08.057.

114. Groen, C.M., Podratz, J.L., Pathoulas, J., Staff, N., and Windebank, A.J. (2022). Genetic Reduction of Mitochondria Complex I Subunits is Protective against Cisplatin-Induced Neurotoxicity in Drosophila. J Neurosci 42, 922–937. 10.1523/jneurosci.1479-20.2021.

115. Picard, M., McEwen, B.S., Epel, E.S., and Sandi, C. (2018). An energetic view of stress: Focus on mitochondria. Frontiers in neuroendocrinology 49, 72–85. 10.1016/j.yfrne.2018.01.001.

116. La Manno, G., Gyllborg, D., Codeluppi, S., Nishimura, K., Salto, C., Zeisel, A., Borm, L.E., Stott, S.R.W., Toledo, E.M., Villaescusa, J.C., et al. (2016). Molecular Diversity of Midbrain Development in Mouse, Human, and Stem Cells. Cell 167, 566–580 e519. 10.1016/j.cell.2016.09.027.

117. Phillips, R.A., 3rd, Tuscher, J.J., Black, S.L., Andraka, E., Fitzgerald, N.D., Ianov, L., and Day, J.J. (2022). An atlas of transcriptionally defined cell populations in the rat ventral tegmental area. Cell Rep 39, 110616. 10.1016/j.celrep.2022.110616.

118. Sim, H., Lee, J.E., Yoo, H.M., Cho, S., Lee, H., Baek, A., Kim, J., Seo, H., Kweon, M.N., Kim, H.G., et al. (2020). Iroquois Homeobox Protein 2 Identified as a Potential Biomarker for Parkinson’s Disease. Int J Mol Sci 21. 10.3390/ijms21103455.

119. Tiklova, K., Gillberg, L., Volakakis, N., Lunden-Miguel, H., Dahl, L., Serrano, G.E., Adler, C.H., Beach, T.G., and Perlmann, T. (2021). Disease Duration Influences Gene Expression in Neuromelanin-Positive Cells From Parkinson’s Disease Patients. Front Mol Neurosci 14, 763777. 10.3389/fnmol.2021.763777.

120. Teh, C.H., Loh, C.C., Lam, K.K., Loo, J.M., Yan, T., and Lim, T.M. (2007). Neuronal PAS domain protein 1 regulates tyrosine hydroxylase level in dopaminergic neurons. J Neurosci Res 85, 1762–1773. 10.1002/jnr.21312.

121. Doitsidou, M., Flames, N., Topalidou, I., Abe, N., Felton, T., Remesal, L., Popovitchenko, T., Mann, R., Chalfie, M., and Hobert, O. (2013). A combinatorial regulatory signature controls terminal differentiation of the dopaminergic nervous system in C. elegans. Genes Dev 27, 1391–1405. 10.1101/gad.217224.113.

122. Yee, C.L., Wang, Y., Anderson, S., Ekker, M., and Rubenstein, J.L. (2009). Arcuate nucleus expression of NKX2.1 and DLX and lineages expressing these transcription factors in neuropeptide Y(+), proopiomelanocortin(+), and tyrosine hydroxylase(+) neurons in neonatal and adult mice. J Comp Neurol 517, 37–50. 10.1002/cne.22132.

123. Garritsen, O., van Battum, E.Y., Grossouw, L.M., and Pasterkamp, R.J. (2023). Development, wiring and function of dopamine neuron subtypes. Nat Rev Neurosci 24, 134–152. 10.1038/s41583-022-00669-3.

124. Di Salvio, M., Di Giovannantonio, L.G., Omodei, D., Acampora, D., and Simeone, A. (2010). Otx2 expression is restricted to dopaminergic neurons of the ventral tegmental area in the adult brain. Int J Dev Biol 54, 939–945. 10.1387/ijdb.092974ms.

125. Panman, L., Papathanou, M., Laguna, A., Oosterveen, T., Volakakis, N., Acampora, D., Kurtsdotter, I., Yoshitake, T., Kehr, J., Joodmardi, E., et al. (2014). Sox6 and Otx2 control the specification of substantia nigra and ventral tegmental area dopamine neurons. Cell Rep 8, 1018–1025. 10.1016/j.celrep.2014.07.016.

126. Di Giovannantonio, L.G., Di Salvio, M., Acampora, D., Prakash, N., Wurst, W., and Simeone, A. (2013). Otx2 selectively controls the neurogenesis of specific neuronal subtypes of the ventral tegmental area and compensates En1-dependent neuronal loss and MPTP vulnerability. Dev Biol 373, 176–183. 10.1016/j.ydbio.2012.10.022.

127. Reyes, S., Fu, Y., Double, K.L., Cottam, V., Thompson, L.H., Kirik, D., Paxinos, G., Watson, C., Cooper, H.M., and Halliday, G.M. (2013). Trophic factors differentiate dopamine neurons vulnerable to Parkinson’s disease. Neurobiol Aging 34, 873–886. 10.1016/j.neurobiolaging.2012.07.019.

128. Tang, X., Jiao, L., Zheng, M., Yan, Y., Nie, Q., Wu, T., Wan, X., Zhang, G., Li, Y., Wu, S., et al. (2018). Tau Deficiency Down-Regulated Transcription Factor Orthodenticle Homeobox 2 Expression in the Dopaminergic Neurons in Ventral Tegmental Area and Caused No Obvious Motor Deficits in Mice. Neuroscience 373, 52–59. 10.1016/j.neuroscience.2018.01.002.

129. Chung, C.Y., Licznerski, P., Alavian, K.N., Simeone, A., Lin, Z., Martin, E., Vance, J., and Isacson, O. (2010). The transcription factor orthodenticle homeobox 2 influences axonal projections and vulnerability of midbrain dopaminergic neurons. Brain 133, 2022–2031. 10.1093/brain/awq142.

130. Di Salvio, M., Di Giovannantonio, L.G., Acampora, D., Prosperi, R., Omodei, D., Prakash, N., Wurst, W., and Simeone, A. (2010). Otx2 controls neuron subtype identity in ventral tegmental area and antagonizes vulnerability to MPTP. Nat Neurosci 13, 1481–1488. 10.1038/nn.2661.

131. Lopert, P., and Patel, M. (2014). Brain mitochondria from DJ-1 knockout mice show increased respiration-dependent hydrogen peroxide consumption. Redox Biol 2, 667–672. 10.1016/j.redox.2014.04.010.

132. Zondler, L., Miller-Fleming, L., Repici, M., Goncalves, S., Tenreiro, S., Rosado-Ramos, R., Betzer, C., Straatman, K.R., Jensen, P.H., Giorgini, F., and Outeiro, T.F. (2014). DJ-1 interactions with alpha-synuclein attenuate aggregation and cellular toxicity in models of Parkinson’s disease. Cell Death Dis 5, e1350. 10.1038/cddis.2014.307.

133. Jimenez-Harrison, D., Huseby, C.J., Hoffman, C.N., Sher, S., Snyder, D., Seal, B., Yuan, C., Fu, H., Wysocki, V., Giorgini, F., and Kuret, J. (2023). DJ-1 Molecular Chaperone Activity Depresses Tau Aggregation Propensity through Interaction with Monomers. Biochemistry 62, 976–988. 10.1021/acs.biochem.2c00581.

134. Bonifati, V., Rizzu, P., van Baren, M.J., Schaap, O., Breedveld, G.J., Krieger, E., Dekker, M.C., Squitieri, F., Ibanez, P., Joosse, M., et al. (2003). Mutations in the DJ-1 gene associated with autosomal recessive early-onset parkinsonism. Science 299, 256–259. 10.1126/science.1077209.

135. Guzman, J.N., Sanchez-Padilla, J., Wokosin, D., Kondapalli, J., Ilijic, E., Schumacker, P.T., and Surmeier, D.J. (2010). Oxidant stress evoked by pacemaking in dopaminergic neurons is attenuated by DJ-1. Nature 468, 696–700. 10.1038/nature09536.

136. Hao, L.Y., Giasson, B.I., and Bonini, N.M. (2010). DJ-1 is critical for mitochondrial function and rescues PINK1 loss of function. Proc Natl Acad Sci U S A 107, 9747–9752. 10.1073/pnas.0911175107.

137. Lin, Z., Chen, C., Yang, D., Ding, J., Wang, G., and Ren, H. (2021). DJ-1 inhibits microglial activation and protects dopaminergic neurons in vitro and in vivo through interacting with microglial p65. Cell Death Dis 12, 715. 10.1038/s41419-021-04002-1.

138. Franco, R., and Cidlowski, J.A. (2009). Apoptosis and glutathione: beyond an antioxidant. Cell death and differentiation 16, 1303–1314. 10.1038/cdd.2009.107.

139. Hastings, T.G., Lewis, D.A., and Zigmond, M.J. (1996). Role of oxidation in the neurotoxic effects of intrastriatal dopamine injections. Proc Natl Acad Sci U S A 93, 1956–1961. 10.1073/pnas.93.5.1956.

140. Friggi-Grelin, F., Coulom, H., Meller, M., Gomez, D., Hirsh, J., and Birman, S. (2003). Targeted gene expression in Drosophila dopaminergic cells using regulatory sequences from tyrosine hydroxylase. J Neurobiol 54, 618–627. 10.1002/neu.10185 [doi].

141. Berry, J.A., Cervantes-Sandoval, I., Chakraborty, M., and Davis, R.L. (2015). Sleep Facilitates Memory by Blocking Dopamine Neuron-Mediated Forgetting. Cell 161, 1656–1667. 10.1016/j.cell.2015.05.027.

142. Diao, F., Ironfield, H., Luan, H., Diao, F., Shropshire, W.C., Ewer, J., Marr, E., Potter, C.J., Landgraf, M., and White, B.H. (2015). Plug-and-play genetic access to drosophila cell types using exchangeable exon cassettes. Cell reports 10, 1410–1421. 10.1016/j.celrep.2015.01.059.

143. Sherer, L.M., Catudio Garrett, E., Morgan, H.R., Brewer, E.D., Sirrs, L.A., Shearin, H.K., Williams, J.L., McCabe, B.D., Stowers, R.S., and Certel, S.J. (2020). Octopamine neuron dependent aggression requires dVGLUT from dual-transmitting neurons. PLoS Genet 16, e1008609. 10.1371/journal.pgen.1008609.

144. Choi, B.J., Imlach, W.L., Jiao, W., Wolfram, V., Wu, Y., Grbic, M., Cela, C., Baines, R.A., Nitabach, M.N., and McCabe, B.D. (2014). Miniature neurotransmission regulates Drosophila synaptic structural maturation. Neuron 82, 618–634. 10.1016/j.neuron.2014.03.012.

145. Aguilar, J.I., Dunn, M., Mingote, S., Karam, C.S., Farino, Z.J., Sonders, M.S., Choi, S.J., Grygoruk, A., Zhang, Y., Cela, C., et al. (2017). Neuronal Depolarization Drives Increased Dopamine Synaptic Vesicle Loading via VGLUT. Neuron 95, 1074–1088 e1077. 10.1016/j.neuron.2017.07.038.

146. Shearin, H.K., Macdonald, I.S., Spector, L.P., and Stowers, R.S. (2014). Hexameric GFP and mCherry reporters for the Drosophila GAL4, Q, and LexA transcription systems. Genetics 196, 951–960. 10.1534/genetics.113.161141.

147. Long, X., Colonell, J., Wong, A.M., Singer, R.H., and Lionnet, T. (2017). Quantitative mRNA imaging throughout the entire Drosophila brain. Nat Methods 14, 703–706. 10.1038/nmeth.4309.

148. Hirose, F., Ohshima, N., Kwon, E.J., Yoshida, H., and Yamaguchi, M. (2002). Drosophila Mi-2 negatively regulates dDREF by inhibiting its DNA-binding activity. Molecular and cellular biology 22, 5182–5193. 10.1128/mcb.22.14.5182-5193.2002.

149. Freyberg, Z., Sonders, M.S., Aguilar, J.I., Hiranita, T., Karam, C.S., Flores, J., Pizzo, A.B., Zhang, Y., Farino, Z.J., Chen, A., et al. (2016). Mechanisms of amphetamine action illuminated through optical monitoring of dopamine synaptic vesicles in Drosophila brain. Nature communications 7, 10652 10.1038/ncomms10652.

150. Schindelin, J., Arganda-Carreras, I., Frise, E., Kaynig, V., Longair, M., Pietzsch, T., Preibisch, S., Rueden, C., Saalfeld, S., Schmid, B., et al. (2012). Fiji: an open-source platform for biological-image analysis. Nat Methods 9, 676–682. 10.1038/nmeth.2019.

151. Croset, V., Treiber, C.D., and Waddell, S. (2018). Cellular diversity in the Drosophila midbrain revealed by single-cell transcriptomics. eLife 7. 10.7554/eLife.34550.

152. Park, A., Croset, V., Otto, N., Agarwal, D., Treiber, C.D., Meschi, E., Sims, D., and Waddell, S. (2022). Gliotransmission of D-serine promotes thirst-directed behaviors in Drosophila. Curr Biol 32, 3952–3970.e3958. 10.1016/j.cub.2022.07.038.

153. Thurmond, J., Goodman, J.L., Strelets, V.B., Attrill, H., Gramates, L.S., Marygold, S.J., Matthews, B.B., Millburn, G., Antonazzo, G., Trovisco, V., et al. (2019). FlyBase 2.0: the next generation. Nucleic acids research 47, D759–d765. 10.1093/nar/gky1003.

154. Stuart, T., Butler, A., Hoffman, P., Hafemeister, C., Papalexi, E., Mauck, W.M., 3rd, Hao, Y., Stoeckius, M., Smibert, P., and Satija, R. (2019). Comprehensive Integration of Single-Cell Data. Cell 177, 1888–1902.e1821. 10.1016/j.cell.2019.05.031.

155. Kilkenny, C., Browne, W.J., Cuthill, I.C., Emerson, M., and Altman, D.G. (2010). Improving bioscience research reporting: the ARRIVE guidelines for reporting animal research. PLoS Biol 8, e1000412. 10.1371/journal.pbio.1000412.

156. Muntifering, M., Castranova, D., Gibson, G.A., Meyer, E., Kofron, M., and Watson, A.M. (2018). Clearing for Deep Tissue Imaging. Current protocols in cytometry 86, e38. 10.1002/cpcy.38.

157. Susaki, E.A., Tainaka, K., Perrin, D., Yukinaga, H., Kuno, A., and Ueda, H.R. (2015). Advanced CUBIC protocols for whole-brain and whole-body clearing and imaging. Nature protocols 10, 1709–1727. 10.1038/nprot.2015.085.

158. Watson, A.M., Rose, A.H., Gibson, G.A., Gardner, C.L., Sun, C., Reed, D.S., Lam, L.K.M., St Croix, C.M., Strick, P.L., Klimstra, W.B., and Watkins, S.C. (2017). Ribbon scanning confocal for high-speed high-resolution volume imaging of brain. PLoS One 12, e0180486. 10.1371/journal.pone.0180486.

159. Sofroniew, N., Lambert, T., Evans, K., Nunez-Iglesias, J., Bokota, G., Winston, P., Peña-Castellanos, G., Yamauchi, K., Bussonnier, M., Doncila Pop, D., et al. (2022). napari: a multi-dimensional image viewer for Python (v0.4.17rc8). Zenodo. 10.5281/zenodo.7276432.

160. Tyson, A.L., Vélez-Fort, M., Rousseau, C.V., Cossell, L., Tsitoura, C., Lenzi, S.C., Obenhaus, H.A., Claudi, F., Branco, T., and Margrie, T.W. (2022). Accurate determination of marker location within whole-brain microscopy images. Sci Rep 12, 867. 10.1038/s41598-021-04676-9.

161. Tyson, A.L., Rousseau, C.V., Niedworok, C.J., Keshavarzi, S., Tsitoura, C., Cossell, L., Strom, M., and Margrie, T.W. (2021). A deep learning algorithm for 3D cell detection in whole mouse brain image datasets. PLoS computational biology 17, e1009074. 10.1371/journal.pcbi.1009074.

162. Howard, J., and Gugger, S. (2020). Fastai: A Layered API for Deep Learning. Information 11, 108.

163. Rocklin, M. (2015). Dask: Parallel computation with blocked algorithms and task scheduling. (SciPy Austin, TX), pp. 136.

164. Gopinath, A., Badov, M., Francis, M., Shaw, G., Collins, A., Miller, D.R., Hansen, C.A., Mackie, P., Tansey, M.G., Dagra, A., et al. (2021). TNFα increases tyrosine hydroxylase expression in human monocytes. NPJ Parkinsons Dis 7, 62. 10.1038/s41531-021-00201-x.

165. Weisel, F.J., Zuccarino-Catania, G.V., Chikina, M., and Shlomchik, M.J. (2016). A Temporal Switch in the Germinal Center Determines Differential Output of Memory B and Plasma Cells. Immunity 44, 116–130. 10.1016/j.immuni.2015.12.004.

166. Love, M.I., Huber, W., and Anders, S. (2014). Moderated estimation of fold change and dispersion for RNA-seq data with DESeq2. Genome Biol 15, 550. 10.1186/s13059-014-0550-8.

167. Hao, Y., Hao, S., Andersen-Nissen, E., Mauck, W.M., 3rd, Zheng, S., Butler, A., Lee, M.J., Wilk, A.J., Darby, C., Zager, M., et al. (2021). Integrated analysis of multimodal single-cell data. Cell 184, 3573–3587 e3529. 10.1016/j.cell.2021.04.048.

168. Stuart, T., Butler, A., Hoffman, P., Hafemeister, C., Papalexi, E., Mauck, W.M., 3rd, Hao, Y., Stoeckius, M., Smibert, P., and Satija, R. (2019). Comprehensive Integration of Single-Cell Data. Cell 177, 1888–1902 e1821. 10.1016/j.cell.2019.05.031.

169. Seney, M.L., Huo, Z., Cahill, K., French, L., Puralewski, R., Zhang, J., Logan, R.W., Tseng, G., Lewis, D.A., and Sibille, E. (2018). Opposite Molecular Signatures of Depression in Men and Women. Biol Psychiatry 84, 18–27. 10.1016/j.biopsych.2018.01.017.

170. Zhou, Y., Zhou, B., Pache, L., Chang, M., Khodabakhshi, A.H., Tanaseichuk, O., Benner, C., and Chanda, S.K. (2019). Metascape provides a biologist-oriented resource for the analysis of systems-level datasets. Nat Commun 10, 1523. 10.1038/s41467-019-09234-6.

171. Korotkevich, G., Sukhov, V., Budin, N., Shpak, B., Artyomov, M.N., and Sergushichev, A. (2021). Fast gene set enrichment analysis. bioRxiv, 060012. 10.1101/060012.

172. Heinz, S., Benner, C., Spann, N., Bertolino, E., Lin, Y.C., Laslo, P., Cheng, J.X., Murre, C., Singh, H., and Glass, C.K. (2010). Simple combinations of lineage-determining transcription factors prime cis-regulatory elements required for macrophage and B cell identities. Mol Cell 38, 576–589. 10.1016/j.molcel.2010.05.004.

173. Vissers, J.H., Manning, S.A., Kulkarni, A., and Harvey, K.F. (2016). A Drosophila RNAi library modulates Hippo pathway-dependent tissue growth. Nat Commun 7, 10368. 10.1038/ncomms10368.

